# Biobank-scale genotyping of Robertsonian translocations reveals hidden structural variation on the human acrocentric chromosomes

**DOI:** 10.64898/2026.03.08.710242

**Authors:** Arang Rhie, Juhyun Kim, Francisco Rodriguez-Algarra, Steven J. Solar, Sergey Koren, Dmitry Antipov, Caralynn M. Wilczewski, George L. Maxwell, Jennifer Gerton, Justin Paschall, Tamara Potapova, Tyra G. Wolfsberg, Sumeeta Singh, Sandra O. del Castillo del Rio, Human Pangenome Reference Consortium, Clesson Turner, Vardhman K. Rakyan, Adam M. Phillippy

## Abstract

Balanced Robertsonian translocation (ROB) is the most common chromosomal rearrangement in humans, with an estimated occurrence of 1 in 800 in newborn studies. Carriers are at increased risk of cancer and often diagnosed at fertility clinics after facing recurrent miscarriages, infertility, or aneuploid offspring. Genotyping carriers with DNA sequencing has been challenging because of gaps and misrepresentation of the translocation fusion site in the human reference genome. Only recently, telomere-to-telomere (T2T) human genomes successfully revealed sequences of the acrocentric short arms, including the most common ROB fusion site. A ROB results in loss of two ribosomal DNA (rDNA) arrays and its adjacent distal sequences, including the highly conserved distal junction (DJ). Here, we present a new method to type ROB carriers directly from short and long sequencing reads by estimating DJ copy number. We demonstrate that our method successfully genotypes ROBs using a reference-free approach or alignments to either T2T-CHM13v2 or GRCh38. Applying this method to a cohort of healthy newborns and family members (n=4,172) as well as the UK Biobank (n=490,416), we find candidate ROBs at a frequency consistent with the previously reported incidence (0.11-0.12%). In addition to ROB carriers, we report the frequency of one DJ loss (9, 2.8-3.4%) or gain (11+, 8.4-9.3%) from the two cohorts and the 1000 Genomes Project (n=3,202), and characterize the underlying structural variation in near-T2T genome assemblies from the Human Pangenome Reference Consortium. Importantly, we provide the first screening method for ROB carriers using low-coverage sequencing data, enhancing its clinical applicability and enabling new studies of structural variation on the acrocentric chromosomes.

## INTRODUCTION

Robertsonian translocation is a type of chromosomal fusion involving acrocentric or telocentric chromosomes. In humans, a Robertsonian chromosome (ROB) occurs at a rate of 1 in 800 individuals^1^, with the most common form of translocation found between chromosome 14 and either chromosome 13 or 21, accounting for 75% and 10% of cases, respectively^2,3^. If a person carries a ROB with two normal chromosomes to pair, it is considered a “balanced translocation (ROB carrier)” as opposed to monosomy or trisomy. For example, one fused ROB between 14 and 21 is still able to pair with the unaffected copies of 14 and 21 during meiosis. However, such carriers are at increased risk of childhood leukemia, non-Hodgkin lymphoma and breast cancer^4^. The carriers are also at increased risk to produce gametes with an unbalanced karyotype, often leading to spontaneous miscarriage, infertility, and uniparental disomy or trisomy in the offspring^5^. After meiosis of a cell containing a single ROB between chromosomes A and B the resulting haploid gamete can carry six possible configurations: A, B, AB, rob(AB), rob(AB)+A, rob(AB)+B. Only AB and rob(AB) are balanced, whereas the other configurations result in either a monosomy or trisomy. This explains both the subfertility of ROB carriers as well as their increased risk of children with trisomy. Homozygous ROB carriers are rare but possible if both parents transmit the same ROB.^6^ The extra 21q copy from an inherited ROB has been reported to cause 2-4% of Down syndrome cases^7^. Similarly but less frequently, an additional 13q copy transmitted from a ROB causes Patau syndrome.^8^ Therefore, early carrier identification is essential to enable risk-stratified cancer surveillance, timely clinical intervention, and informed reproductive counseling.

The exact mechanism and breakpoint of the fusion site forming the most common type of ROB remained unknown until recently. In the past, the ROB fusion breakpoint was reported to be localized in HSat3^9^, and later corrected between HSat1 and the rDNA^10,11^. Nevertheless, sequence-level identification of ROB carriers has been hampered by the incomplete and misassembled acrocentric regions in the standard human reference genomes (both in hg19 and GRCh38). In 2022, the first complete telomere-to-telomere (T2T) human genome revealed the complex structure of the acrocentric p-arms, including regions with high sequence homology^12^. In a follow-up study, using human pangenome assemblies^13^, frequent inter-chromosomal exchange was inferred within so-called pseudo-homolog regions (PHR)^14^. A particular PHR on the proximal short arms of Chrs. 13, 14, and 21 was suggested to mediate ROB translocations^15^, which was later confirmed by the T2T sequencing and assembly of three ROB cell lines^16^. Such ROB formation results in the loss of one fusion site and sequences distal to it, which includes the rDNA gene array on each chromosome and the highly conserved distal junction (DJ) sequence of each rDNA array^14,16,17^.

The most widely used methods for identifying ROB carriers have been karyotyping, fluorescence in situ hybridization (FISH)^18^ and GTG-banding, which are typically performed only after clinical indications, such as infertility or recurrent pregnancy loss, have been identified. However, these methods require obtaining live cells, processing, imaging, and analysis by trained cytogeneticists, making them incompatible with high-throughput whole-genome sequencing (WGS) workflows or retrospective analyses of extracted and biobanked DNA samples. More recently, a proprietary microarray based chip^19^ and Hi-C methods^20^ have been developed to detect ROB carriers for retrospective screening. However, the lack of understanding of the exact ROB breakpoint and gaps in the prior human reference genomes (i.e. both GRCh38 and hg19), has limited the use of genotyping or sequencing data, including WGS reads, for the detection of ROB carriers^21^.

Recognizing that ROBs result in the predictable loss of acrocentric sequence along with the increasing large-scale cohorts sequenced with WGS available, we sought to explore methods to screen ROB carriers from such datasets. Here, we show three ways to detect ROBs from short-read sequencing data by using (1) T2T-CHM13/hs1 as an alternate reference for read alignment, (2) pre-existing read alignments mapped to GRCh38, and (3) DJ marker *k*-mers and their multiplicity directly obtained from sequencing reads. We benchmarked these methods to reference genomes CHM13 and HG002 as controls and three known ROB cell lines as cases^12,16^. Next, we applied the method on WGS in the Reverse Phenotyping Core (RPC) cohort (n=4,172)^22,23^ and the UK Biobank WGS cohort (n=490,444)^24^. We report that these methods identify potential ROB carriers in all cohorts at approximately the expected frequency of 1 in 800. Finally, we typed all samples from the 1000 Genomes Project (1KGP)^25^ and further examined samples with atypical copies of the DJ in the recent, near-T2T HPRC Release 2 assemblies^26^. We propose this method as a routine screening approach for identifying potential ROB carriers in any whole-genome sequencing dataset.

## RESULTS

### Detecting ROBs by the loss of rDNA distal junctions

We first identified the most stable, highly conserved region to target for copy number estimates. The most common form of ROB translocation is reported to have a breakpoint in the proximal short arm, between the rDNA and centromere^16^. Therefore, ROB carriers are expected to be missing two rDNA arrays and all sequences distal to these arrays (**Figure 1A**). Because of the difficulty in detecting the loss of rDNA units due to their highly variable copy number, we targeted the region immediately distal of the rDNA array, the DJ, which had been reported to be highly conserved in humans and expected to be in a single copy per chromosome^17^. Surveying the copy number estimates from the Simons Genome Diversity Project^12,27^, we confirmed that the DJ region was the only suitable region maintaining a stable 10 copy pattern across individuals regardless of ancestral background, suggesting the presence of a highly conserved, single DJ per rDNA array (**Figure 1B**). Thus, we hypothesized that two DJ copies will be lost following most Robertsonian translocations, resulting in eight DJs in the ROB carriers. Using this hypothesis, we developed a pipeline for read-mapping based copy number estimates utilizing a masked T2T-CHM13 reference (**Figure 1C, S1**). Specifically, we masked all but one copy of the DJ in the T2T-CHM13 reference and compared the coverage against the background coverage collected from the rest of the non-heterochromatic autosomal regions. In addition, we masked all but one copy of the 45S rDNAs and Robertsonian-associated PHRs, allowing these regions to be used as target regions for copy number estimation (**Table S1 and Figure S1-4**).

**Figure 1.**
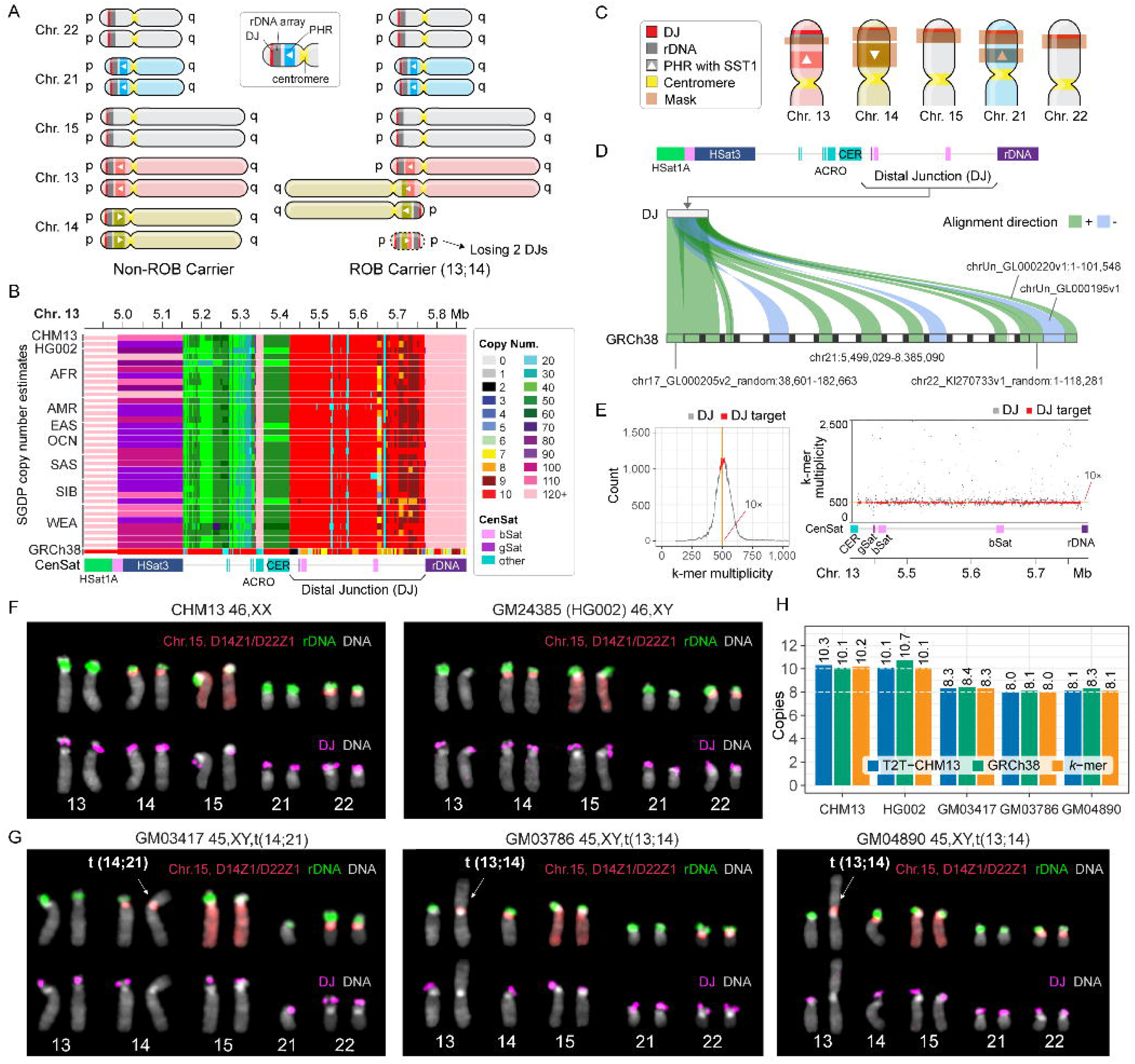
Target rDNA distal junction (DJ) for ROB carrier detection. (A) Schematic overview of the most common karyotype of ROB carriers. Chr. 14 pseudo-homolog region (PHR) has been identified in its inverted orientation compared to those on Chrs. 13 and 21, reported as the fusion site for the most common forms of ROBs. When a ROB is formed, one small segment carrying one PHR and two DJs gets lost. Example ideogram shows a t(13;14) ROB. (B) Copy number estimates of the DJ show conserved 10-20 copies across individuals from various ancestral backgrounds in the Simons Genome Diversity Project (SGDP). Copy numbers for GRCh38 (at bottom) indicate this region is mis-assembled in GRCh38. SGDP population labels: AFR, Africa; AMR, America; EAS, East Asia; OCN, Oceania; SAS, South Asia; SIB, Siberia; WEA, Western Eurasia. (C) Masked regions of the T2T-CHM13v2.0 acrocentric p-arms for copy number estimation of the DJ, rDNA, and PHR. Only one copy of each in Chr. 13 has been chosen as the target for copy number estimates. See also Figure S1-4 and Table S1. (D) Identifying DJ target region for copy number estimates for sequencing read alignments on GRCh38. Nine mis-assembled copies of the target were found on Chr. 21 surrounded by gaps, two copies on unplaced contigs and two on unlocalized contigs of Chrs. 17 and 22. See also Tables S2-4. (E) Reference free DJ copy number estimates. *K*-mers found in all five DJs in T2T-CHM13v2.0 were selected as “DJ *k*-mers”. Based on the *k*-mer copy number count in Illumina reads (left), *k*-mers with ∼10× coverage were selected as “DJ target *k*-mers”. The *k*-mer multiplicity (num. of times a k-mer is found in the entire read set, similar to read coverage) on CHM13 Chr. 13 shows a stable 10× coverage across the region for the DJ target *k*-mers. See also Figure S5. (F) Control cell lines CHM13 and GM24385 (HG002). All 10 acrocentric haplotypes show the presence of rDNA (green) and DJ (magenta) with FISH. (F, G) Whole chromosome painting of Chr. 15 and centromeric satellite 14/22 (CenSat D14Z1/D22Z1) probe was applied to distinguish each of the acrocentric chromosomes (red). (G) Previously reported ROB cell lines, in which the breakpoint has been identified as the PHR in de Lima *et al*.^16^ The ROB chromosomes show the absence of the rDNA (green) and DJ (magenta). (H) DJCounter confirms 10 DJs in samples shown in panel F and 8 DJs in panel G. See also Table S5.

To expand the pipeline on large-scale cohorts already mapped to GRCh38, we sought to look for the presence of the target DJ sequence in GRCh38. When mapping GRCh38 to each target sequence extracted from CHM13, we identified nine DJ copies on GRCh38 chr21 organized as a repeat array with some inverted and incomplete copies interspersed and surrounded with gaps (**Figure 1D**). This is an arrangement seen in no T2T-assembled genome since, and represents a clear mis-assembly in GRCh38. In addition, partial DJ sequences were found on two unplaced sequences (chrUn_GL000195v1 and chrUn_GL000220v1) and two unlocalized sequences (chr22_KI270733v1_random and chr17_GL000205v2_random), with high sequence identity (average 96.2%), and were included as target regions for quantifying DJ copy number in GRCh38 (**Tables S2-3** and **Figure S1**). Diverged pieces of DJs were found on Chrs. 4 and 10, similar to those observed in CHM13, however these were excluded from targeting due to their low sequence identity (<83.4%) and low coverage (<2.8%). Using these reference-specific DJ targets, we implemented three distinct modes that differ in their normalization methods and read-filtering strategies (**Table S4**).

In addition to the mapping-based method, we also developed a reference-free approach to reduce any potential mapping bias. To select target *k*-mers for the DJ, all *k*-mers (*k*=31) in T2T-CHM13v2 were evaluated for their copy number multiplicity observed in Illumina reads derived from CHM13 (**Figure S1**). Among the *k*-mers found at least once in each of the five DJs (Chrs. 13, 14, 15, 21 and 22), target *k*-mers were selected from the ∼10 copy peak to exclude highly repetitive or variable regions (**Figure 1E**). To estimate the copy number of the DJs for each new sample, all *k*-mers are collected directly from the FASTQ files. The two-copy peak is then extracted from the entire *k*-mer histogram and compared to the median multiplicity of the target DJ *k*-mers. Since this method operates directly on FASTQ files, it eliminates the need for a mapping process, thus reducing computational resource requirements.

To test the pipelines, we used Illumina reads from CHM13, HG002 (GM24385)—for which 10 DJs were confirmed^12,28^—and ROB cell lines GM03417, GM03786, and GM04890—for which 8 DJs and ROBs were confirmed by T2T assemblies and metaphase FISH by de Lima *et al.*^16^ (**Figure 1F,G**). We applied our mapping-based method using both GRCh38 and CHM13 reference genomes, utilizing BAM files aligned to each reference, as well as our *k*-mer-based method using raw sequencing reads. All three methods consistently reported ∼10 DJs for both CHM13 and HG002 and ∼8 DJs for the 3 known ROB cell lines, as expected (**Figure 1H and S5**). Regardless of the methods used, GM03417 showed elevated copies of the DJ (∼8.3), reflecting the cellular heterogeneity (**Table S5**). Roughly, 10% of the cells had an additional copy of Chr. 21 present in the cell-line^16^.

### One in 800 ROB candidates in newborn family cohort

The Reverse Phenotyping Core (RPC) cohort includes 1,325 families with healthy newborns, parents, and siblings (where available), totalling 4,187 samples. Blood samples were collected at the time of birth. Illumina short read data aligned to hg19 from Wilczewski *et al.*^23^ and Bodian *et al.*^22^ were extracted and re-aligned to the masked version of T2T-CHM13v2.0 and evaluated for the copy number of the DJs, PHRs and rDNAs (**Figure 2**). In total, we found 4 individuals with estimated 8 copies of the DJs (**Figure 2A**). This includes two parents and one mother-newborn (daughter) pair (**Table 1**). Counting the 3 out of 2,650 presumably unrelated parents, among which 2,612 were sequenced, yields an estimated ratio of 1 in 871-883 (0.11%), matching closely the expected 1 in 800 (0.125%) incidence^1^. Among the 2,612 parents, 2,277 (87.17%) appeared as having 10 estimated copies, comprising the majority of the individuals (**Figure 2A**). Interestingly, regardless of with or without the offspring, we found 3.34-3.45% individuals with 9 DJs, and 9.26-9.27% with more than 10 DJs, with 13 being the most copies observed (**Figure 2 and Table 1**). When comparing the number of samples classified in each DJ copy bin, no significant difference was observed between XX and XY individuals (χ^2^ test of independence, *p*>0.5). We observed a few more samples with 8 DJs in XX carriers (3 vs. 1), with one transmitted from the mother to daughter. Although the sample size is too small to draw firm conclusions, this observation is consistent with previous reports that ROBs are more frequently transmitted from mother to XX offspring^29^. When comparing the distribution of DJ copy number between parents and offspring, we observed a slight shift toward higher copy numbers in the parental genomes (Mann–Whitney U test, p = 3.7 × 10□□; **Figures 2B and S6**).

**Figure 2.**
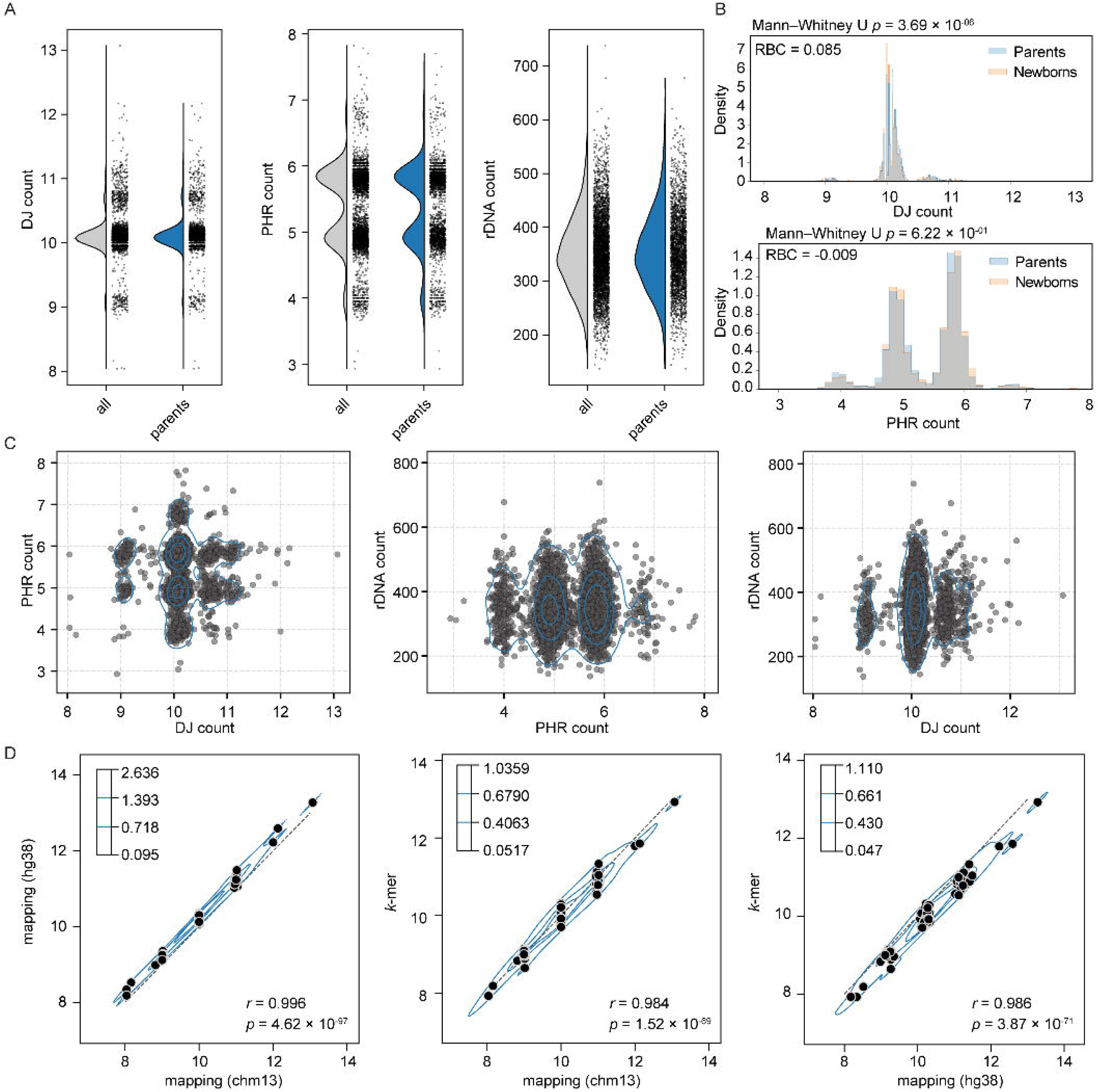
DJ, PHR, and rDNA copy number estimation using T2T-CHM13, GRCh38, and *k*-mer based methods for the RPC cohort. (A) Diploid copy number estimation of DJ, PHR, and rDNA in the RPC cohort using CHM13-based mapping approach. (B) Histogram of the probability density of DJ and PHR copy numbers across samples, stratified by parents and newborns. The area under each group’s histogram sums to one. A two-sided Mann–Whitney U test was performed using raw counts to compare parents and newborns, and the rank-biserial correlation (RBC) is reported as the effect size. See also Figure S6. (C) Each point represents a sample from the RPC cohort. No obvious correlations were observed between the DJ, PHR, and rDNA copy numbers. The data density is shown by the blue contour lines. (D) Comparison between DJ counts derived from mapping-based estimates (CHM13 and GRCh38) and the *k*-mer–based method using subsampled samples (n = 92), showing strong correlation. Density levels are shown with four contour levels. See also Figure S7.

**Table 1.** ROB candidates from the RPC cohort. Estimated num. of DJs, PHRs, and rDNAs are shown below with the categorized copy counts binned. ROB candidates are highlighted with a green background.

| Family | Sex | DJ | Bin | PHR | Bin | rDNA | Relationship |
| --- | --- | --- | --- | --- | --- | --- | --- |
| A | XX | 8.16 | 8 | 3.87 | 4 | 386.6 | Mother |
|  | XY | 10.09 | 10 | 4.85 | 5 | 433.8 | Father |
|  | XY | 8.88 | 9 | 4.71 | 5 | 415.6 | Child |
| B | XX | 10.2 | 10 | 6.00 | 6 | 338.3 | Mother |
|  | XY | 8.04 | 8 | 4.04 | 4 | 229.6 | Father |
|  | XX | 9.10 | 9 | 5.40 | 5 | 357.7 | Child-A |
|  | XX | 8.95 | 9 | 5.31 | 5 | 334.8 | Child-B |
| C | XX | 8.05 | 8 | 4.77 | 5 | 254.7 | Mother |
|  | XY | 9.04 | 9 | 4.85 | 5 | 325.6 | Father |
|  | XX | 8.04 | 8 | 5.80 | 6 | 316.2 | Child-A |
|  | XX | 9.15 | 9 | 4.90 | 5 | 424.9 | Child-B |

However, when comparing the number of PHRs, we observed a broader polymorphic copy number distribution compared to the DJs (**Figure 2A**). Initially, we expected that the majority of individuals would carry six copies of the PHRs as observed in CHM13 and HG002, and five copies for those with 8 DJs due to the ROB as reported in Guarracino *et al*.^14^ and de Lima *et al.*^16^. A little over half the samples were observed as 6 copies (52.3% in all and 56.6% among parents), still forming the largest frequency, followed by 5, 4 and 7 copies (39.3-35.6%, 6.2-5.5% and 2.0-2.3%, respectively for all and among parents; **Table S6**). Among the parental genomes, the portion of 6 PHR copies observed was slightly higher in samples carrying XX over XY (57.9% vs. 55.2%). For 5 and 4 copies, however, the observed frequency was slightly higher in XY carriers over XX (36.5% vs. 34.7% and 5.8% vs. 5.1%). Unlike the DJ, the distribution of PHR copy numbers was not significantly different between parents and their offsprings (Mann–Whitney U test, *p*>0.5. **Figure 2B**). The copy number of the PHRs in the 8 DJ samples also varied from 4 to 6 (**Table 1**). Distribution of the PHR copy number also showed no significant relationship with the DJ gains or losses and showed a very weak correlation when restricted to samples with 8 to 10 DJ copies (Pearson correlation, *r*=0.01, *p*=0.39), indicating there could be other polymorphic variations of the PHRs that are yet to be discovered (**Figure 2C**).

Next, we examined the changes in rDNA copy number associated with DJ loss or gain. The rDNA copy number varied widely among samples, ranging from 137 to 738 copies (**Figure 2A**). Unlike the DJ or PHR distribution, we observed a unimodal peak with a wide deviation (mean=352, sd=70.2) as previously reported for the UK Biobank cohort^30^. Again, the rDNA copy number distribution showed a very weak positive correlation for samples with 8 to 10 DJ copies (DJ < 10.4; Pearson correlation: r = 0.09, p = 8.1 × 10□□). However, for samples with 10 or more copies, copy number ranges were similar and less correlated (DJ >= 10.4, Pearson r = 0.12, p = 1.4 × 10□²). (**Figure 2C**). No significant correlation was observed between PHR copy number and rDNA copy numbers (p > 0.05) (**Figure 2C**).

We then evaluated whether our two alternative approaches—using GRCh38 as the reference or using marker *k*-mers—yielded high correlation with the CHM13 mapping-based approach. We selected samples representing each classified DJ copy number category, including all samples with 8 DJ copies and all samples with more than 12 DJ copies (n = 92 total). Among the GRCh38-based modes, we selected the “Fast-precise” mode (more details in **“<u>DJCounter as a tool for predicting DJ copies</u>”** section and **<u>Methods</u>**), which showed the highest correlation with CHM13-based estimates and the lowest residual sum of squares (RSS) when fitting DJ copy numbers derived from GRCh38 alignments (**Figure S7 and Table S4**). All three approaches showed similar distributions and strong correlations (Pearson correlation r² and *p*-values shown in **Figure 2D**), demonstrating the robustness and broad applicability of these methods across different data types.

### DJ and PHR copy number distribution in UK Biobank

To confirm the distribution of the DJ copy numbers and the PHRs in an independent cohort, we applied our analysis to the UK Biobank (n=490,444), for which CRAM files aligned to GRCh38 are available (Byrska-Bishop *et al.*, Methods^25^). To enable DJ and PHR copy number estimation in this large cohort, we applied the mapping-based approach of our pipeline to reduce computational demands (“pilot Fast mode” in **Methods and Figure S8**). Among the samples, we were able to calculate DJ and PHR counts for 490,369 and 490,374 samples, respectively. The distribution pattern and percent samples assigned to each discrete DJ and PHR count replicated what we observed in the RPC cohort, both when considering all individuals and when restricted to unrelated individuals (**Figure 3A**). In total, we identified 583 samples with 8 DJ copies, corresponding to a predicted ROB frequency of 1 in 841 (0.12%), again closely matching the expected 1 in 800. Unlike the RPC cohort, we observed some individuals with 7 DJ counts. More individuals with 3 PHR counts were observed, presumably due to the larger size of the cohort, with a frequency of 0.05% regardless of the sex chromosome composition, and the lowest PHR count observed 2, in a single sample. Again, we did not observe a statistically significant difference in the frequency of each discrete copy number bin between sex (χ^2^ test of independence, *p*>0.5, **Table 2**). Similar to the RPC cohort, we did not observe differences between sexes. We also confirmed that DJ counts were significantly (albeit slightly) higher not only in the parent group compared to the offspring (**Figure 3B**), but also in older groups compared to younger groups (**Figure S9**). Consistent with the RPC cohort, we did not observe differences in PHR distribution between parents and offspring (**Figure 3B**).

**Figure 3.**
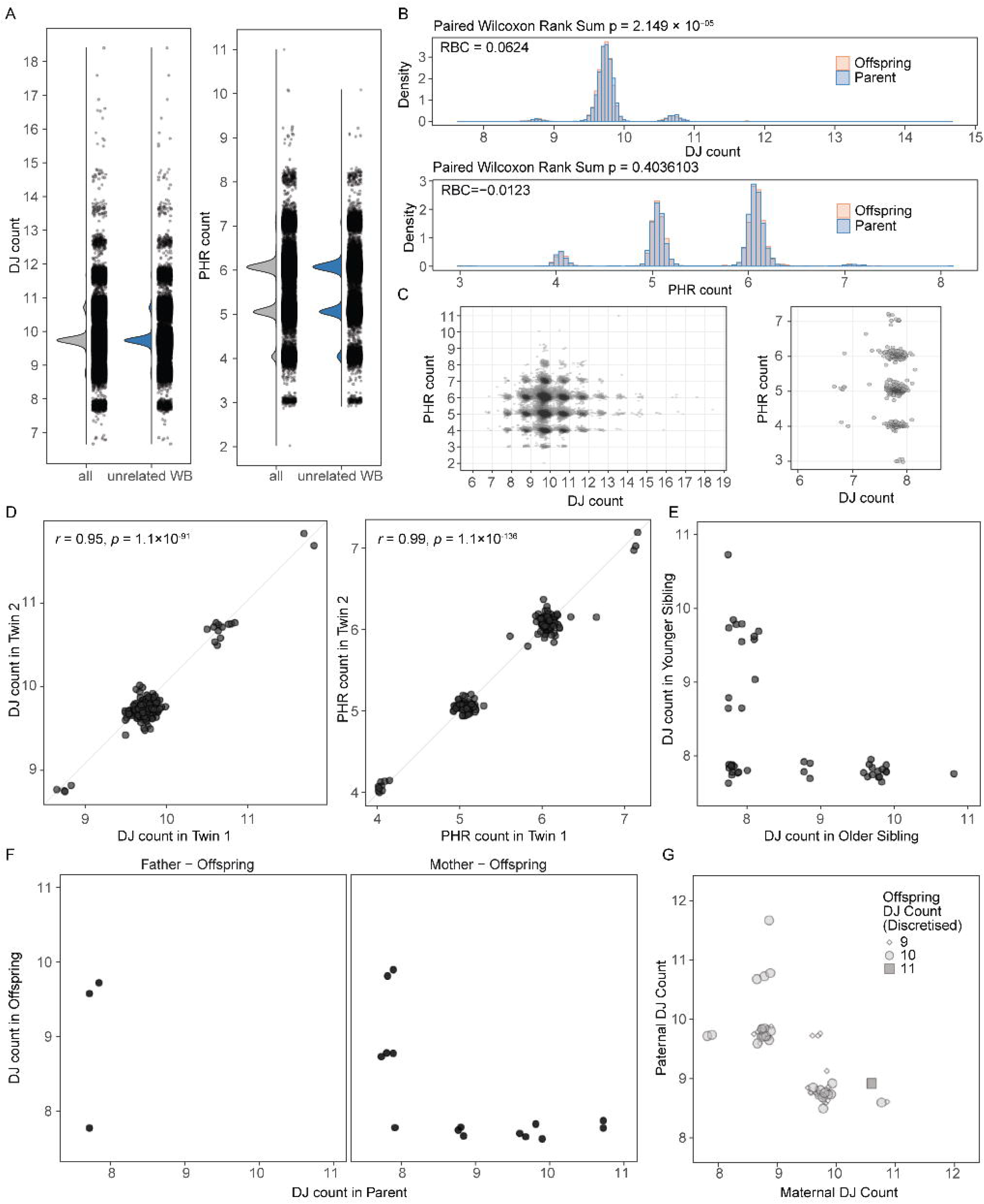
DJ and PHR counts in the UK Biobank. (A) DJ and PHR count distribution among all and unrelated white British (WB) samples. See also Figure S8. (B) Density distribution of DJ (top) and PHR (bottom) counts between the parent group and the offspring group. The RBC was calculated using the paired Wilcoxon Rank Sum test. See also Figure S9. (C) DJ copies as a function for PHR copy numbers. Left, for all DJs. Right, inset of the samples classified as 8 or lower. (D) DJ (left) and PHR (right) counts in monozygotic twins. The twins share the same copy number (Pearson’s r test). (E) DJ counts in siblings (except monozygotic twins) with one of the siblings determined to have 8 DJs. No particular inheritance patterns were found. (F) Inheritance pattern of all 8 DJ carriers found with father-offspring or mother-offspring relationships. No 8 DJ offspring with both parents were found in the cohort. (G) Inheritance pattern of the DJ in offspring among the available trios, with at least one member of the trio having 9 or fewer estimated DJs. No trio with 9 DJs in both parents was observed. See also Table S7.

**Table 2.** Copy number estimates of the DJs across the RPC cohort (n=4,187) and UK Biobank WGS cohort (n=490,369). Regardless of the cohorts, samples with 8 copies of DJs were observed at the lowest frequency (0.11-0.12%). The largest fraction of samples was observed with 10 DJs (87.17-88.70%), followed by 11 DJs (7.81-9.00%), 9 DJs (2.81-3.45%), and more than 11 DJs (0.26-0.56%), indicating loss of one or gains of one or more DJs are relatively common. See also Table S6.

| Cohort | Num. of DJs | Num. of samples | % | XX | % | XY | % |
| --- | --- | --- | --- | --- | --- | --- | --- |
| RPC | <8 | 0 | 0.00% | 0 | 0.00% | 0 | 0.00% |
|  | **8** | **4** | **0.10%** | **3** | **0.14%** | **1** | **0.05%** |
|  | 9 | 140 | 3.34% | 81 | 3.82% | 59 | 2.86% |
|  | 10 | 3,659 | 87.39% | 1,848 | 87.05% | 1,811 | 87.74% |
|  | 11 | 373 | 8.91% | 186 | 8.76% | 187 | 9.06% |
|  | >11 | 11 | 0.26% | 5 | 0.24% | 6 | 0.29% |
|  | Total | 4,187 | 100.00% | 2,123 | 100.00% | 2,064 | 100.00% |
| RPC<br>(Parents only) | <8 | 0 | 0.00% | 0 | 0.00% | 0 | 0.00% |
|  | **8** | **3** | **0.11%** | **2** | **0.15%** | **1** | **0.08%** |
|  | 9 | 90 | 3.45% | 51 | 3.91% | 39 | 2.98% |
|  | 10 | 2,277 | 87.17% | 1,135 | 87.04% | 1,142 | 87.31% |
|  | 11 | 235 | 9.00% | 113 | 8.67% | 122 | 9.33% |
|  | >11 | 7 | 0.27% | 3 | 0.23% | 4 | 0.31% |
|  | Total | 2,612 | 100.00% | 1,304 | 100.00% | 1,308 | 100.00% |
| UK Biobank | <8 | 9 | 0.002% | 6 | 0.002% | 3 | 0.00% |
|  | **8** | **583** | **0.12%** | **312** | **0.12%** | **271** | **0.12%** |
|  | 9 | 13,778 | 2.81% | 7,460 | 2.80% | 6,318 | 2.82% |
|  | 10 | 434,951 | 88.70% | 235,927 | 88.70% | 199,024 | 88.70% |
|  | 11 | 38,291 | 7.81% | 20,791 | 7.82% | 17,500 | 7.80% |
|  | >11 | 2,757 | 0.56% | 1,500 | 0.56% | 1,257 | 0.56% |
|  | Total | 490,369 | 100.00% | 265,996 | 100.00% | 224,373 | 100.00% |

With the increased number of samples carrying 8 DJ copies, we were able to more precisely evaluate the relationship between PHR and DJ copy number. The overall distribution mirrored that observed in the RPC cohort, with a broader PHR copy number distribution among individuals with 10 DJ copies, again presumably due to the larger sample size (**Figure 3C**). A higher proportion of individuals with 8 or fewer DJ copies had 5 PHR copies, as we initially hypothesized (54.4%, compared with 37.8% in individuals with 9 or more DJs). However, substantial individuals carried 3, 4, 6 or 7 copies of the PHRs (45.6%), suggesting other structural variations may be present altering the PHR counts. For example, prior HPRC assemblies showed 34.5% of the Chr. 14 haplotypes lacked the PHR (n=9 out of 26 haplotypes; de Lima *et al.*^16^, Extended Data Figure 9), supporting our observation of variable PHR copy numbers in the population.

We further confirmed that monozygotic twins have identical DJ and PHR copy numbers (**Figure 3D**), whereas dizygotic twins or siblings show differences in DJ copy number inheritance pattern (**Figure 3E**), confirming the precision of our method. Among samples carrying 8 DJs, 9 samples had either the maternal or paternal genome available. Among the 3 paternal samples with 8 DJs, we found 1 case where the offspring inherited the same pattern (**Figure 3F left**). Among the 6 maternal samples, 1 offspring had inherited 8 DJs (**Figure 3F right**). The copy number of the offspring matched one of the parents in most cases, or were at 10 copies in most cases when one of the parents had ≥11 copies (**Figure 3G and Table S7**). Three samples were found to have 9 DJs while both parental genomes had 10 DJs, suggesting a possible *de novo* loss of the DJ in the offspring. No cases were observed in which both parents had 9 DJs among the available trios.

### Uncovering structural variation of the DJ in near-T2T genomes

For detailed inspection of structural gains or losses in DJ copy number, we analyzed 1KGP samples both with and without near-T2T assemblies generated by the HPRC. The 1KGP samples have recently been re-sequenced using Illumina short reads (30×, n=3,202 among which n=2,504 are unrelated)^25^. A subset of the samples (n=226) have been long-read sequenced by the HPRC to build a reference pangenome from *de novo* assembled haplotypes. The cell lines were selected to broadly sample global genomic variation and were restricted to lines with a low passage count and normal karyotype to avoid artifacts from cell culture. These lines were assembled with Verkko^31^ using long reads, Hi-C and parental data when available^26^.

We downloaded the Illumina BAM files aligned to GRCh38 from IGSR^25^ and counted the DJ copy numbers using the GRCh38 and *k*-mer based methods. To assess compatibility between methods and across independent cohorts, we applied previously derived linear models for each GRCh38 mode, using subsampled RPC data to transform DJ counts obtained from GRCh38 alignment into the CHM13-based distribution. This model was subsequently used to transform DJ counts in an independent 1KGP cohort. Among the evaluated approaches, the “Fast-precise” mode demonstrated the lowest RSS (**Table S4**). Moreover, it was the only method for which the transformed 1KGP distribution was statistically indistinguishable from the original RPC distribution generated using the masked T2T-CHM13 method (**Figure 4A and S10**). Therefore, we chose to use the “Fast-precise transformed” DJ counts for further analyses (**Figure 4A**). We obtained a similar distribution as found in the RPC or UK Biobank samples, albeit with a slightly higher fraction of individuals binned in 8, 9 or 11+ DJ copies (**Figure 4A**, **S11 and Table S8**). Here, we also observed a pattern in which the parents have a distribution shifted slightly higher than the offspring, consistent with our findings from the RPC cohort and UK Biobank samples (**Figure 4A,B**).

**Figure 4.**
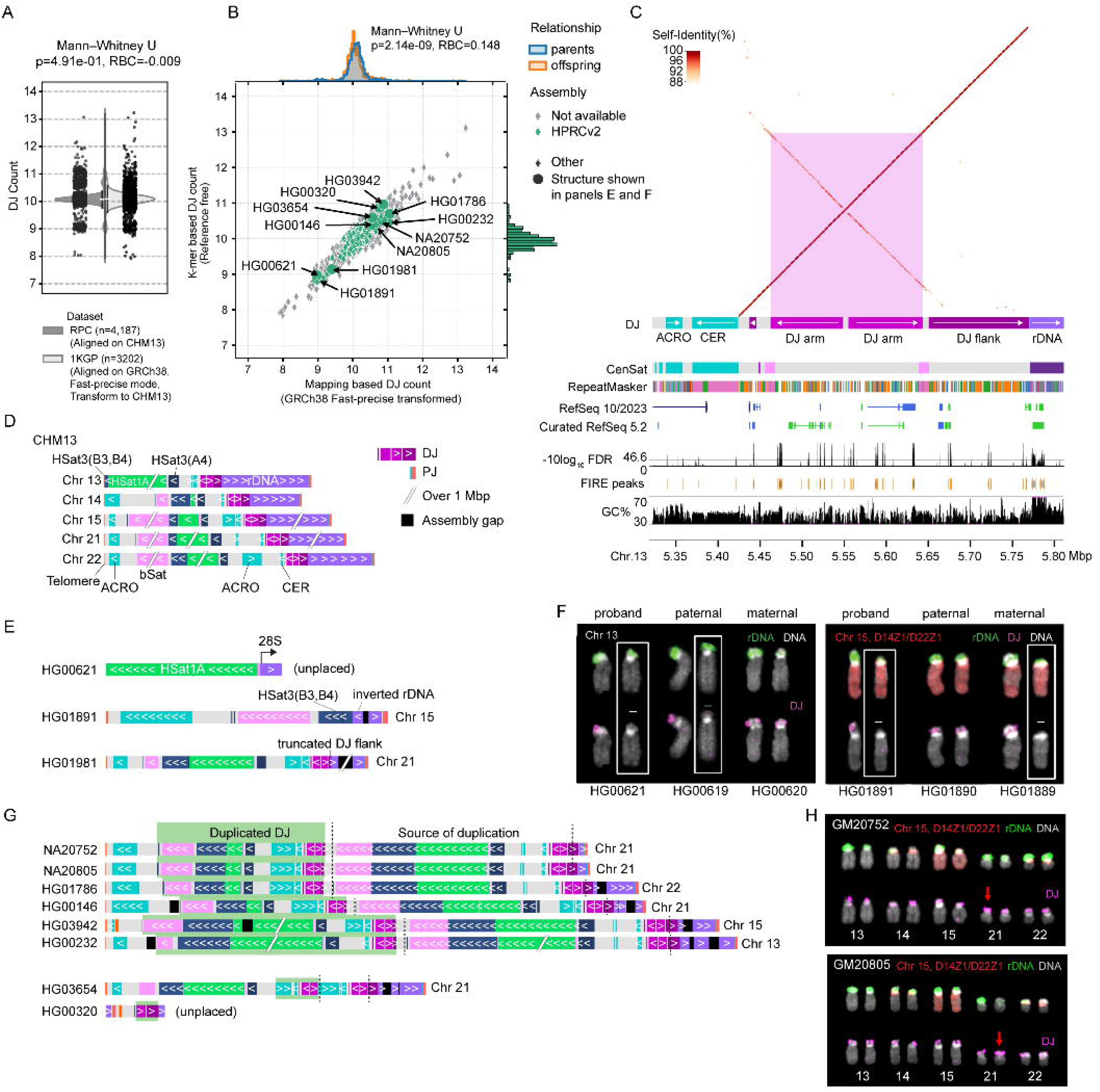
Polymorphism of the rDNA DJs and surrounding sequence on the short arms of the acrocentric chromosomes. (A) Distribution of DJ copy numbers in the RPC cohort, calculated using masked CHM13 as the ground truth, and in the 1KGP cohort, transformed from GRCh38 alignments (Fast-precise mode). See also Figure S10. (B) Distribution of DJ copy numbers across 1KGP samples (n = 3,202), with samples assembled by HPRC highlighted in green. The normalized density plot (top) shows a slight shift of parental samples toward higher values among those with ≥10 copies. The right marginal plot displays the distribution of samples with assemblies. See also Figure S11-12 and Tables S8-9. (C) Structure of the DJ and its functional components. The self-dotplot on top shows the palindromic arms inside the DJ, marked as DJ arm. The region flanking to the rDNA (DJ flank) shows a relatively unique pattern, with the beginning showing diverged inverted repeat shared at the beginning of the DJ (next to the CER). Tracks below show repeat, RefSeq gene and chromatin accessibility annotations. Inconsistent protein-coding (blue) and pseudogene (green) annotations reflect the lack of understanding of the genes in this region. Open chromatin accessibility pattern indicates possible functional activity. RefSeq 10/2023, RefSeq release of T2T-CHM13 from NCBI. Curated RefSeq 5.2, curated annotation from GRCh38^40^. FDR and FIRE peaks, fiber-seq inferred regulatory element peaks and false discovery rate in Phred scale, in which higher scores indicate better support^41^. (D) Distal sequences and repeats as observed in CHM13. All DJ sequences are intact. Chr. 13 does not have the sequence below the bSat unlike the other sequences. (C,D,E,G) See also Tables S10-11 and Figure S13-14. (E) Ideograms of cases with loss of DJs. HG00621 had an unplaced sequence containing the loss of DJ. Sequences lost include the beginning of the rDNA array down to the bSat, resulting in loss of sequences including the entire DJ, CER, ACRO and HSat3(A4). The rDNA array started at the large subunit (28S). HG01891 lost sequences including the HSat1A, with a rare rDNA inversion found at the beginning of the rDNA array. HG01981 had only a partial loss of the DJ flank. Sequences larger than 1 Mbp are reduced and re-scaled for clarity. (F) FISH with probes for the DJs (green) and rDNAs (magenta) and chromosome painting for Chr. 15 (red) on the 2 cases found to have 9 DJs. HG00621 exhibited the same DJ loss pattern as the paternal genome (chr13), while HG01891 shared the pattern observed in the maternal genome (chr15). FISH probes and chromosome painting follows Figure 1. (G) Ideograms of cases with gain of DJs. Six samples shared the same duplicated form of DJ (green box), sharing the same breakpoint at the DJ flank (dotted lines). The HSat1A block size varied, however the overall structure of the duplicated copy (green box) remained identical. The 2 samples at the bottom showed a unique structure, with one having the DJ palindromes duplicated and the inverted repeat part of the DJ flank. The last case shows a sequence within the rDNA array, with one rDNA array (left) having the PJ intact. This time, the additional DJ has only one of the DJ arms present with the DJ flank. See also Figure S15. (H) FISH for the two samples with 11 DJs. Because the DJs are too close to each other, and highly packed inside the heterochromatin, the duplicated DJs aren’t clearly distinguishable. Still, one haplotype (red arrow) exhibits stronger intensity compared to the other. The other haplotype of the same chromosome was assembled with a single DJ flanking the rDNA array. FISH probes and chromosome painting follows Figure 1.

Seven individuals in the 1KGP dataset were classified as having 8 DJs, suggesting possible ROB carriers (**Table S9**). All seven samples were identified within pedigreed families, allowing us to confirm the inheritance pattern. Among the four families, one family showed a potential ROB was inherited in the father-son relationship (HG01204-HG01206), and in the mother-daughter relationship (HG00651-HG00652). Two potential ROB carriers were found only in the parental genomes and not observed in their offspring. Interestingly, one offspring had 8.1-8.2 copies, while neither parent carried 8 DJ copies, indicative of a possible *de novo* ROB. Alternatively, given the parents are both classified as 9 copies, it is also possible that this child inherited two DJ-less acrocentric chromosomes. Unfortunately, these samples were not selected by the HPRC and so never underwent karyotyping. However, one sample, HG01204, was initially selected for sequencing by the HPRC and underwent G-banding for quality control. This revealed a karyotypic aberration involving a fusion between Chrs. 13 and 21 (**Figure S12**), which is an extremely rare type of a ROB chromosomal fusion with only a few case reports available to date (for example, Chen *et al.*^32^). HPRC ultimately excluded this sample due to its abnormal karyotype, but this example highlights the ability of our method to correctly identify a true ROB from a large biobank.

Since no 8 DJ cases were sequenced by the HPRC, we evaluated all remaining cases of abnormal DJ counts. Six samples with 9 predicted DJs and ten samples with 11 predicted DJs were included in the HPRCr2 assemblies, allowing us to further inspect the sequence-level structure of the DJs and the surrounding region. To confirm the entirety of the DJ structure, we divided the DJ target into 3 regions—two palindromic arms residing close to the CER repeat and the region flanking to the rDNA (DJarms and DJflank in **Figure 4C**). We searched for the DJ, rDNAs and other satellite repeats commonly found on the short arms of the acrocentric chromosomes (**Figure 4D and Table S10**). Next, we investigated the assembly graph to confirm that the distal sequences were unambiguously assembled and connected to the rDNA nodes (**Table S11**). HiFi or ONT reads were mapped to the junction to validate the loss of DJs.

Among the 6 samples with 9 DJ copies, 3 samples successfully assembled the junction with the loss of DJ (**Figure 4E** and **Figure S13**). In the first case, HG00621, the DJ was lost entirely, including the HSat3 (A4) distal to the DJ and the beginning of the rDNA array. The rDNA array started at the 28S subunit rather than the regular 5’ external transcribed spacer. FISH painting of the DJs for this family showed the loss of DJ was also observed in the paternal genome, HG00619 (**Figure 4F**). In the second case, HG01891, we found the DJ was also lost entirely, including part of HSat3 (**Figure 4E**). The beginning of the rDNA array was found inverted with a gap at the inversion breakpoint within the array. A local re-assembly of the inverted rDNA junction was confirmed with ONT UL reads, suggesting this is a true inversion of the rDNA (**Figure S14**). The chromosome painting showed the absence of DJ was also present in the maternal genome on Chr. 15 (**Figure 4F**). The third case, HG01981, had DJ counts of 9.1-9.4, indicating slightly higher counts for the samples grouped as 9 DJs (**Figure 4E**). We found the DJ was mostly intact except for the sequence flanking to the rDNA array on Chr. 21. Given that all 3 cases are showing non-overlapping breakpoints, we conclude that loss of 1 DJ is less common, however still transmissible within families.

When investigating the 8 samples with extra DJ junction assembled, we found 7 samples assembled the entire distal sequence to the telomere, including the duplicated DJ (**Figure 4G** and **Figure S15**). Within those samples, we found the second DJ was partially duplicated with the breakpoint residing internal to the flanking sequence, including distal sequences. Six samples shared a similar segmentally duplicated block spanning sequences down to the large beta satellite block. The DJ arm was truncated, with only the beginning 34 kb part being duplicated. This truncated DJ flank sequence is shared downstream the DJ, as an inverted repeat, with long-noncoding transcripts and pseudogenes. The duplicated DJ in NA20752 (GM20752) and NA20805 (GM20805) was not visible in FISH, likely due to their proximity at this scale (**Figure 4H**). Arguably, one haplotype of Chr. 21 showed stronger intensity for the DJ probes, suggestive for a possible duplication. The second case, found in HG03654, had a block duplicated but starting with a smaller portion of the DJ flank down to the ACRO repeat (**Figure 4G**). The last case, HG00320, was partially assembled, surrounded by rDNA sequences. In this sample, the partial DJ contained only one arm of the DJ palindromes, with the flanking sequence intact to the rDNA. Given that we see 6 cases with overlapping segmentally duplicated breakpoints ending at the same DJ flank sequence, we conclude this form is the more common form present among the extra DJ carriers.

### DJCounter as a tool for predicting DJ copies

Considering that large-scale sequencing studies predominantly rely on short reads, utilizing the supported GRCh38 reference genome with target DJ regions (**Tables S1, S3**) in most cases, we developed our method into a tool “DJCounter” to enable accurate detection of ROBs and determine the DJ copy numbers, all while maintaining reasonable computational demands (**Figure S1**).

DJCounter accepts either alignment files (BAM/CRAM aligned to CHM13 or GRCh38) or sequencing reads (FASTQ) as input files. When reads mapped to a supported reference are available, alignment-based analysis is recommended. In this method, read coverage is collected for the target DJ and autosomes. Three modes are provided that differ in duplicate-read filtering and background coverage estimation (**Table S4**). When reads are not aligned, or aligned to an unsupported reference, *k*-mer based analysis is recommended. For *k*-mer based analysis, the sample’s *k*-mers are intersected with our marker DJ *k*-mers to estimate the target k-mer multiplicity and normalized by the background two-copy multiplicity. In both analysis methods, the output is an estimated DJ copy number for a diploid genome. Normal samples typically have a value of approximately 10, whereas ROB carriers typically show values around eight. As a lightweight wrapper, DJCounter requires minimal computational resources, making it readily applicable to large-scale cohort studies.

Here, we summarize the various filtering and normalization strategies that were tested for benchmarking purposes (**Table S4**). Initially, we used the most conservative approach, normalizing DJ counts based on read depth from alignments to masked T2T-CHM13 (“High-resolution” mode). As this mode is performed at base-resolution, it required substantial wall-clock time and was not suitable to apply to the UK Biobank dataset. To address this limitation, we tested an alternative “Fast” mode that utilizes read counts directly from the BAM index file. While doing so, we refined the DJ targets (**Table S3**) that were missing in the “pilot-Fast” mode (**Table S2**). Although computationally efficient, this method does not filter out secondary, supplementary or duplicated read alignments, resulting in inconsistent results (batch effects) depending on how the BAM file was generated. The last mode, “Fast-precise” filters out the aforementioned reads prior to estimating DJ counts and uses all filtered reads mapped to the autosomes and normalizes by the total size of the autosomes.

We benchmarked the computational requirements of the three modes for GRCh38 and the *k*-mer based method across different sequencing depths. We subsampled the Illumina NovaSeq reads for HG002^33^ to 1×, 5×, 10×, 20×, 30×, 40×, and 50×. As expected, the “Fast” mode was the most computationally efficient option, requiring substantially less CPU time and maximum peak memory, making it suitable for large cohort studies (**Figure 5A-C and S16**). The “Fast-precise” mode required slightly more computational resources than the “Fast” mode but offered the advantage of minimizing batch effects by relying exclusively on primary alignments. Although the k-mer based method requires higher computational resources compared with alignment-based approaches, it remains more efficient when starting from raw reads avoiding the additional 4.5-200 CPU hours of read mapping, depending on coverage (**Figure S16**). Therefore, the k-mer based method is preferable when unaligned reads or alignments to an unsupported reference are available. The *k*-mer based method operates on Meryl^34^, which can trade shorter CPU wall time for higher memory. The “Fast-precise” mode shows higher DJ count estimates (+0.7), however provides robust results across different sequencing coverages, even at 1× (**Figure 5D**). The *k*-mer based method yields estimates closer to the expected 10 copies at > 10× coverage but becomes unstable below 10× (**Figure 5D**). Nevertheless, these results indicate that both mapping (>5x) and *k*-mer based (>10x) methods are applicable in clinical settings even for low-coverage sequencing data.

**Figure 5.**
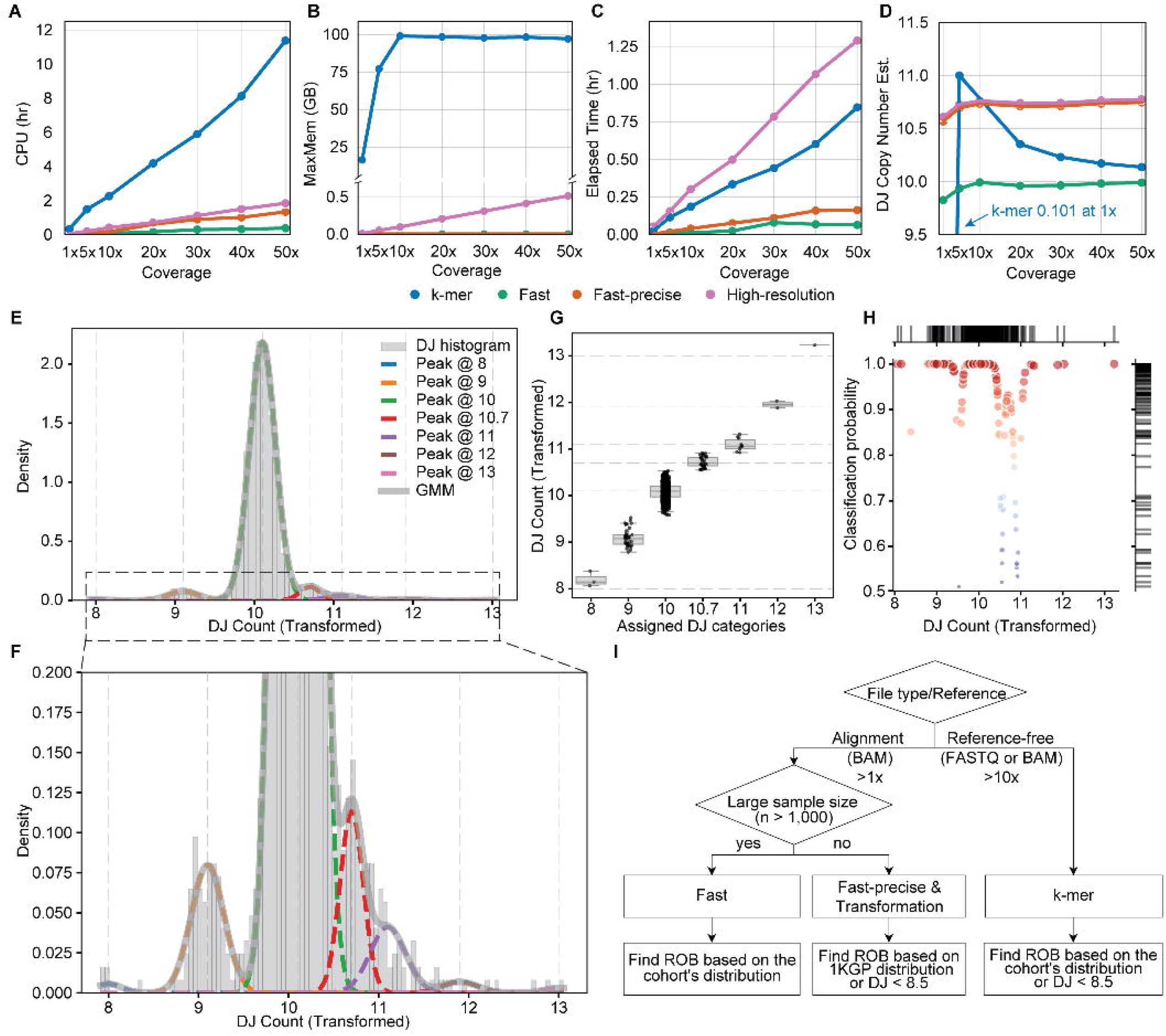
Computational resources required for DJ estimation using GRCh38 alignment–based and *k*-mer approaches, and DJ classification using a GMM trained on 1KGP distribution. (A-C) Computational resources required for DJ calculation across methods, including total CPU time, maximum memory usage (MaxMem) and Elapsed time (wall-clock). See also Table S4 and Figure S16. (D) DJ copy number estimation using different approaches, methods, and sequencing coverages. HG002 short read data was used and subsampled to varying depths. (E) Gaussian mixture model (GMM) fitting on the training set subsampled from 1KGP (n = 2,402) and a (F) Zoomed-in view. See also Figure S17. (G) DJ classification using the trained GMM on the independent test subset of 1KGP (n = 800). See also Figure S18. (H) Transformed DJ copy numbers with corresponding classification probabilities for each DJ category. The color scale corresponds to the y-axis values, with higher values shown in red and lower values shown in blue. (I) Recommendations for using DJCounter based on input format and sample size. See also Figure S1,S16,S19.

A limitation of this approach is that it requires a large cohort to accurately estimate the distribution and identify ROB samples based on that distribution. Therefore, we developed a framework that operates at a single-sample level, enabling calculation of DJ copy number directly from GRCh38 alignments and identifies ROB candidates using a reference density derived from the 1KGP cohort obtained with the “Fast-precise” mode. We constructed a Gaussian mixture model (GMM) to transform DJ copy number estimates using subsamples from the 1KGP cohort (training set, n = 2,402) and validated the model on the remaining samples (test set, n = 800). This framework identifies candidate ROB samples and classifies each sample into a discrete copy number bin, even when only a single sample is available. Due to the uneven density across peaks, we fixed the mean of each component and constrained peak positions to align between the RPC and 1KGP cohorts (**Figure S17**). An additional intermediate component between 10 and 11 copies was included to reflect the partially duplicated DJ we observed (**Figure 5F**) as a separate category (**Figure 5E-F**). Using this model, we could assign a discrete integer bin for each sample in both the 1KGP test set (**Figure 5G**) and the RPC cohort (**Figure S18**), along with posterior classification probabilities. The classification probabilities indicated high confidence (> 0.5) for all samples tested, with the majority of the test set showing strong support (> 0.9), including ROB samples (**Figure 5H**).

For optimal performance, we recommend different modes in DJCounter under different scenarios (**Figure 5I**). For studies with large sample sizes (n > 1,000) with no batch effects expected, where the cohort-specific distribution can be reliably estimated and reads are aligned to GRCh38, we recommend running the “Fast” mode and inspecting the overall distribution to identify potential ROB carriers. However, applying a strict cutoff of 8 copies may be challenging, as the Fast mode includes duplicates and supplementary reads, which can shift the overall distribution. In contrast, for smaller cohorts (n < 1,000) or datasets with potential batch effects, where the underlying distribution cannot be reliably estimated, the “Fast-precise” mode is preferred. In this setting, the classification approach should be applied to DJ estimates generated using the Fast-precise mode, with duplicate reads properly removed or marked in the alignment files. We do not recommend downsampling when alignments are already available to avoid additional overhead in computation. The *k*-mer based method is reference-free and does not require the alignment step. Therefore, it is recommended when rapid diagnosis is needed without alignments.

To assess the applicability of DJCounter, we applied our recommended three methods (**Figure 5I**) to long-read datasets (ONT and HiFi) from HG002 and the three ROB samples, and compared the results with DJ estimates from Illumina data (**Figure S19**). Mapping-based DJ counting showed slightly different estimates across sequencing platforms, with the most accurate values observed for Illumina data when using CHM13 as the reference. Overall, the counts were consistently higher in HiFi and ONT compared to Illumina in all samples. Counts were also higher when using GRCh38 as the reference, likely reflecting mapping biases from sequencing technologies and aligners. In contrast, *k*-mer based results showed minimal differences between Illumina and HiFi, while ONT exhibited lower DJ counts, likely due to lower per-base read accuracy affecting *k*-mer sets. Despite these differences, all methods and sequencing platforms consistently distinguish between normal and ROB samples, which differ by approximately two copies.

## DISCUSSION

Here we used recently completed T2T and near-T2T assemblies along with FISH validation to understand the structure and conservation of the short arms of human acrocentric chromosomes, particularly surrounding the rDNA arrays. Using this information, we developed a new method for genotyping the rDNA distal junction and other features of the short arms to predict the presence of Robertsonian chromosomes. Importantly, our method can be successfully applied to short-read sequencing datasets (>5x), enabling its application to large-scale biobanks, as demonstrated here using the RPC, UK Biobank, and 1KGP datasets.

Given the concordance between the observed frequency and the previously reported ROB prevalence, as well as the observed inheritance patterns, we propose that a DJ copy number of 8 or less is a reliable marker for screening ROB carriers. We also looked into the frequency distribution of the PHR on the proximal short arms of Chrs. 13, 14, and 21, where one copy is expected to be lost after ROB formation. However, a wider range of PHR copy numbers was observed, indicating the polymorphic nature of this sequence, and with no significant association with DJ copy number. Thus, in contrast to the DJ, the copy number of the PHRs does not appear to be a reliable marker of ROBs. Prior work points to the SST1 macrosatellite and an enrichment of PRDM9 motifs within the ROB-associated PHR as possible recombination hotspots, both explaining the higher prevalence of 13;14 and 14;21 ROBs, as well as the high degree of structural variation^16^. Cataloging structural variants and their implications across the acrocentric short arms is an area of future work.

We also found a lower number of DJs in the older group than in the younger group, with the younger group showing counts closer to the expected normal count (∼10), across two independent datasets. We observed a similar pattern in the parent group compared with the child group, with the child group also showing copy numbers closer to the expected count of 10. One possible explanation is the accumulation of structural variants with age in the acrocentric short arms, which undergo more frequent sequence exchange than other chromosomes^14,35^.

The DJ and the structure of the sequences distal to the DJ on the acrocentric short arms are nearly impossible to genotype without near complete genome assemblies built from long reads. The near-T2T HPRCr2 assemblies allowed us to understand the loss and gain of the DJs, revealing previously uncharacterized structural variation in this region. The 9 DJ cases found in the 1KGP set were sporadic, with the DJ-less junction sequence not found in the RPC cohort. The gain of DJs was more frequently observed, with blocks of neighboring sequences duplicated alongside. The recurrent duplication found in 6 out of 8 cases across the 1KGP populations indicate that this is the most common form of duplication, regardless of ancestral background.

Originally, we expected the copy number of the DJs to be found as 10 in most samples, with only a handful of 8 DJs, as an indication for ROBs. However, our genotyping results from the two large cohorts showed a greater variability in the copy number, especially in the 9 or 11 range. Using the learned frequency of the 9 DJs we observed, we estimate a rough false positive rate for having two DJ-less haplotypes could be up to one third of the 8 DJ cases (**Methods**). Alternatively, it is also possible to miss a true ROB carrier when the individual carries the duplicated DJ in one of the non-ROB chromosomes, leading to a rough false negative rate of 7.2%. Copy number analysis alone is limited to distinguish a true ROB carrier from these alternative scenarios. Therefore, we suggest using our tool as a screening method to identify potential ROB carriers, followed by karyotype confirmation. This approach will be particularly useful for screening large cohorts to prioritize samples for confirmatory cytogenetic testing.

Because our method works with WGS data, it can be used to identify Robertsonian translocation carriers in large clinical cohorts, including retrospectively from existing data.^36–38^ A positive DJCounter result could be confirmed by karyotyping, providing a sensitive approach for identifying ROBs that may have been unrecognized contributors to disease progression. Because balanced ROB carriers are phenotypically normal at birth, most remain undiagnosed until clinical implications arise, such as leukemia, lymphoma, and other cancers,^4^ which has hindered characterization of ROB biology. Aside from the fertility complications, the mechanistic link between ROB carriage and these risks remain poorly characterized. Even more so, the health implications of DJ and rDNA array copy number variants is unknown. We further characterized complex structural variation involving different DJ copy numbers across personalized genomes at scale, providing opportunities to uncover previously unrecognized associations. The DJ includes important regulatory elements that have been shown with nucleolar function.^39^ Thus, disruption of the DJ through either duplication or deletion may affect the transcription and function of the adjacent rDNA array. Future studies, aided by the methods and results presented here, are now needed to understand the clinical implications of structural variation on the short arms of the acrocentric chromosomes.

### Limitations

To validate the four 8 DJ individuals we identified in the RPC cohort are ROB carriers, we attempted to contact them to request participation. Despite our efforts, all individuals were unable to be recontacted or declined to participate. Additionally, due to the extensive review processes required to re-contact ROB candidates within the UK Biobank, we instead genotyped the DJ copy numbers in the widely studied 1KGP lymphoblastoid cell lines. The structural variation in the DJ region may contain variants specific to the cell line as reported in the 1KGP dataset.^25^

## Supporting information

Supplemental list of authors

## RESOURCE AVAILABILITY

### Lead contact

Further information and requests for resources should be directed to and will be fulfilled by the lead contact, Arang Rhie.

### Materials availability

This study did not generate new unique reagents or sequencing data.

### Data and code availability

DJCounter is available on GitHub (https://github.com/marbl/DJCounter). The initial DJ counting method for the masked T2T-CHM13 version is available on GitHub (https://github.com/arangrhie/T2T-Ref). The acrocentric haplotypes presented in this study from the Verkko assemblies are available on Zenodo (https://doi.org/10.5281/zenodo.18895340) along with the acrocentric satellite annotation. Original imaging data underlying this manuscript can be accessed from the Stowers Original Data Repository at https://www.stowers.org/research/publications/libpb-2613.

### Consortia

Human Pangenome Reference Consortium (affiliations in Supplemental list of authors) Derek Albracht, Ivan A. Alexandrov, Jamie Allen, Alawi A. Alsheikh-Ali, Nicolas Altemose, Casey Andrews, Dmitry Antipov, Lucinda Antonacci-Fulton, Alexander Arguello, Mobin Asri, Marcelo Ayllon, Jennifer R. Balacco, Floris P. Barthel, Edward A. Belter Jr, Halle D. Bender, Andrew P. Blair, Davide Bolognini, Katherine E. Bonini, Christina Boucher, Guillaume Bourque, Silvia Buonaiuto, Shuo Cao, Andrew Carroll, Ann M. Mc Cartney, Monika Cechova, Mark J.P. Chaisson, Pi-Chuan Chang, Xian Chang, Jitender Cheema, Haoyu Cheng, Claudio Ciofi, Hiram Clawson, Sarah Cody, Vincenza Colonna, Holland C. Conwell, Robert Cook-Deegan, Mark Diekhans, Maria Angela Diroma, Daniel Doerr, Zheng Dong, Danilo Dubocanin, Richard Durbin, Jana Ebler, Evan E. Eichler, Jordan M. Eizenga, Parsa Eskandar, Eddie Ferro, Anna-Sophie Fiston-Lavier, Sarah M. Ford, Willard W. Ford, Giulio Formenti, Adam Frankish, Mallory A. Freeberg, Qichen Fu, Stephanie M. Fullerton, Robert S. Fulton, Shenghan Gao, Yan Gao, Gage H. Garcia, Obed A. Garcia, Joshua M.V. Gardner, Shilpa Garg, Erik Garrison, Nanibaa’ A. Garrison, John E. Garza, Margarita Geleta, Mohammadmersad Ghorbani, Tina A. Graves-Lindsay, Richard E. Green, Carol W. Greider, Cristian Groza, Bida Gu, Andrea Guarracino, Melissa Gymrek, Maximilian Haeussler, Leanne Haggerty, Ira M. Hall, Nancy F. Hansen, Yue Hao, Mohammad Amiruddin Hashmi, David Haussler, Prajna Hebbar, Peter Heringer, Glenn Hickey, Todd L. Hillaker, S. Nakib Hossain, Neng Huang, Sarah E. Hunt, Toby Hunt, Alexander G. Ioannidis, Nafiseh Jafarzadeh, Nivesh Jain, Erich D. Jarvis, Maryam Jehangir, Juan Jiang, Eimear E. Kenny, Juhyun Kim, Bonhwang Koo, Sergey Koren, Milinn Kremitzki, Charles H. Langley, Ben Langmead, Heather A. Lawson, Daofeng Li, Heng Li, Ronghan Li, Wen-Wei Liao, Jiadong Lin, Tianjie Liu, Glennis A. Logsdon, Ryan Lorig-Roach, Jonathan LoTempio Jr, Hailey Loucks, Jane E. Loveland, Jianguo Lu, Shuangjia Lu, Julian K. Lucas, Walfred Ma, Juan F. Macias-Velasco, Kateryna D. Makova, Maximillian G. Marin, Christopher Markovic, Tobias Marschall, Franco L. Marsico, Fergal J. Martin, Mira Mastoras, Capucine Mayoud, Brandy McNulty, Jack A. Medico, Julian M. Menendez, Karen H. Miga, Anna Minkina, Matthew W. Mitchell, Saswat K. Mohanty, Younes Mokrab, Jean Monlong, Shabir Moosa, Avelina Moreno-Ochando, Shinichi Morishita, Jonathan M. Mudge, Katherine M. Munson, Njagi Mwaniki, Nasna Nassir, Chiara Natali, Shloka Negi, Lingbin Ni, Adam M. Novak, Faith Okamoto, Keisuke K. Oshima, Pilar N. Ossorio, Chie Owa, Sadye Paez, Benedict Paten, Clelia Peano, Adam M. Phillippy, Brandon D. Pickett, Laura Pignata, Nadia Pisanti, David Porubsky, Pjotr Prins, Timofey Prodanov, Anandi Radhakrishnan, T. Rhyker Ranallo-Benavidez, Brian J. Raney, Mikko Rautiainen, Alessandro Raveane, Andreas Rechtsteiner, Luyao Ren, Arang Rhie, Fedor Ryabov, Samuel Sacco, Farnaz Salehi, Michael C. Schatz, Laura B. Scheinfeldt, Aarushi Sehgal, William E. Seligmann, Mahsa Shabani, Kishwar Shafin, Shadi Shahatit, Ruhollah Shemirani, Vikram S. Shivakumar, Swati Sinha, Jouni Sirén, Linnéa Smeds, Steven J. Solar, Marco Sollitto, Nicole Soranzo, Andrew B. Stergachis, Marie-Marthe Suner, Yoshihiko Suzuki, Arda Söylev, Ahmad Abou Tayoun, Jack A.S. Tierney, Chad Tomlinson, Francesca Floriana Tricomi, Mohammed Uddin, Matteo Tommaso Ungaro, Rahul Varki, Flavia Villani, Ivo Violich, Mitchell R. Vollger, Brian P. Walenz, Charles Wang, Lisa E. Wang, Ting Wang, Aaron M. Wenger, Conor V. Whelan, Zilan Xin, Zheng Xu, Kai Ye, DongAhn Yoo, Wenjin Zhang, Ying Zhou, Xiaoyu Zhuo, Giulia Zunino

## ACKNOWLEDGMENTS

A.R., J.K., S.J.S., S.K., D.A., A.M.P. were supported by National Human Genome Research Institute (NHGRI) ZIA HG200398. J.K. was supported by a grant of the Korea Health Technology R&D Project through the Korea Health Industry Development Institute (KHIDI), funded by the Ministry of Health & Welfare, Republic of Korea (grant number: RS-2022-KH131838). F.R-A and V.K.R. were supported by grants from the Biotechnology and Biological Sciences Research Council (BB/R00675X/1). F.R-A was additionally supported by a Barts Charity seed grant (G-002983). S.O.C.R was supported by a grant from the Medical Research Council (MR/W007045/1). C.T. and C.M.W. were supported by NHGRI ZIA HG200418-02. T.G.W. and S.S. were supported by NIH grant ZIC HG200345. Members of the Reverse Phenotyping Core are as follows: Felicia Akinwande, MBA, MSN, RN, Leslie G. Biesecker, MD, Jennifer Johnston, PhD, Justin Paschall, MS, Sumeeta Singh, MS, Clesson Turner, MD, Caralynn Wilczewski, PhD, ScM, and Tyra Wolfsberg, PhD. J.G. and T.P. are supported by the Stowers Institute for Medical Research and by NCI R01CA266339. T.P. is supported by NCI R50CA305001. We would like to acknowledge the NHGRI for funding the following grants supporting the creation of the human pangenome reference: U41HG010972, U01HG010971, U01HG013760, U01HG013755, U01HG013748, U01HG013744, R01HG011274, and the Human Pangenome Reference Consortium (BioProject ID: PRJNA730823). This research was supported in part by the Intramural Research Program of the National Institutes of Health (NIH). The contributions of the NIH authors are considered Works of the United States Government. The findings and conclusions presented in this paper are those of the authors and do not necessarily reflect the views of the NIH or the U.S. Department of Health and Human Services. This work utilized the computational resources of the NIH HPC Biowulf cluster (https://hpc.nih.gov). During the preparation of this work the authors used ChatGPT5.3 and Gemini to rephrase text for fluency. After using this tool, the authors reviewed and edited the content as needed and take full responsibility for the content of the published article.

## AUTHOR CONTRIBUTIONS

A.R. and J.K. conceived, developed DJCounter, conducted analysis for the RPC and 1KGP cohorts, and drafted the manuscript. C.M.W., G.L.M., J.P., T.G.W., S.S. and C.T. conceived analysis and administered the RPC dataset. F.R.-A., S.O.C.R and V.K.R. performed analysis on the UK Biobank cohort. A.R., S.J.S. and A.M.P. defined the target regions. A.R., S.J.S. S.K. and D.A. generated the HPRC Verkko assemblies and performed manual inspection to validate the DJ variation. T.P. and J.G. conducted metaphase FISH and chromosome painting. C.M.W. and G.L.M. re-contacted participants for validation. A.M.P. supervised the project and provided critical revisions. All authors reviewed and approved the final manuscript.

## Methods

Detailed methods are provided in the online version of this paper and include the following:

- KEY RESOURCES TABLE
- METHOD DETAILS

## STAR□METHODS

### Key resources table

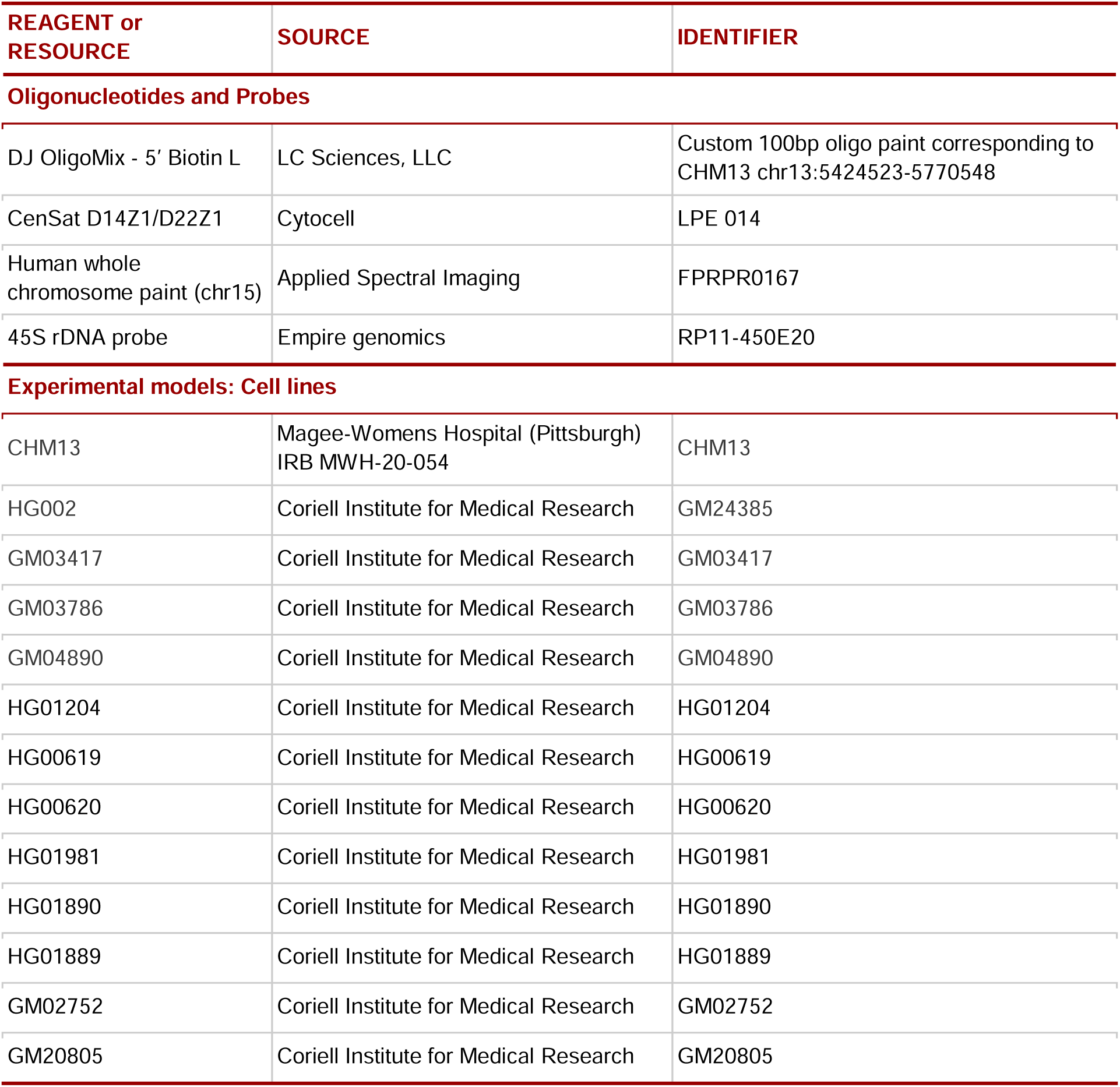
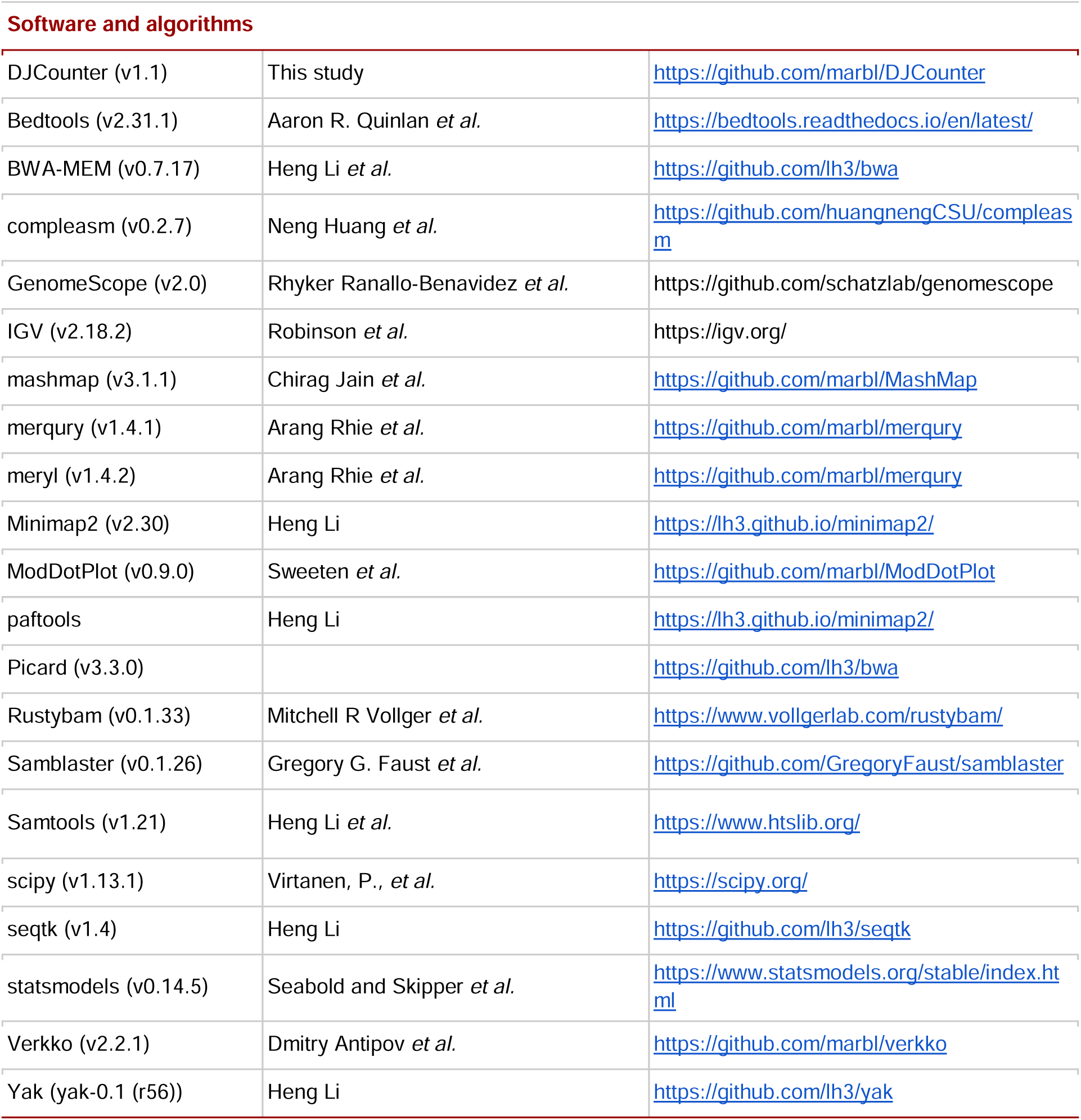

## METHOD DETAILS

### Samples and data

#### Control data

***For CHM13***, PCR-free Illumina Hi-Seq 2500 paired-end reads from Nurk *et al*.^12^ were used. The reads are available on SRA: SRX1009644.

***For HG002***, PCR-free Illumina NovaSeq paired-end reads from Baid *et al*.^33^ were used. The reads are available under google storage: https://console.cloud.google.com/storage/browser/brain-genomics-public/research/sequencing/fastq/novaseq/wgs_pcr_free/50x.

***For the three previously identified ROB samples***, PCR-free Illumina NovaSeq X paired-end reads from de Lima *et al*.^16^ were used. The reads are available under dbGap: phs003920.v1.p1.

#### Tested cohorts

##### Reverse Phenotyping Core cohort

The original Reverse Phenotyping Core (RPC) cohort data included in the present analysis consisted of BAM files per sample, originally collected as part of previously published newborn trio genomic sequencing study^22^ aligned to the hg19 reference genome. We therefore extracted all paired-end reads from the BAM files using samtools fastq and realigned them to either CHM13v2.0 or GRCh38.

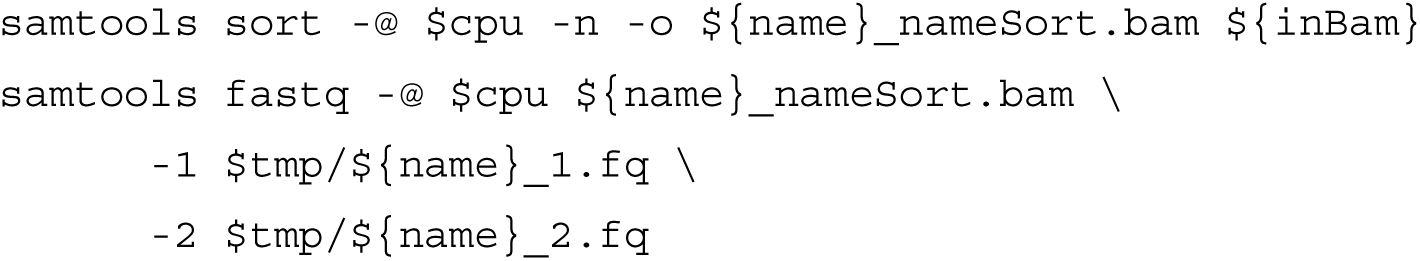

##### UK Biobank

To obtain DJ and PHR count estimates on the UK Biobank, GRCh38 alignment files in `cram` format stored in subfolders of the `/Bulk/GATK and GraphTyper WGS/Whole genome GATK CRAM files and indices [500k release]/` path were accessed through the UK Biobank Research Analysis Platform (UKB RAP, https://www.ukbiobank.ac.uk/use-our-data/research-analysis-platform). The Fast copy number estimation method described below was then executed via the UKB RAP application called swiss-army-knife (SAK) version 4.5.0, with local and remote scripts communicating through the `dxpy` python library.

##### 1KGP

The 1000 Genomes Project dataset 30x Illumina NovaSeq 6000 paired-end reads were obtained from Byrska-Bishop *et al*.^25^. BAM files aligned to GRCh38 were downloaded and processed locally following instructions on IGSR for the 2504 unrelated and the 698 samples: https://www.internationalgenome.org/data-portal/data-collection/1000genomes_30x.

### Target regions for DJ, PHR and rDNAs

Here we describe 3 features for copy number estimates of the distal junction (DJ), pseudo-homolog region (PHR) and the ribosomal DNA (rDNA).

#### Target regions in T2T-CHM13 and masked T2T-CHM13 reference

We used CHM13v2.0 as the reference genome, with the revised Cambridge Reference Sequence (rCRS) included for mitochondrial sequences. Centromeric Satellite (CenSat) repeat annotations from Altemose *et al*.^42^ were used to define target regions of interest. For identifying internal repeat structure, we used ModDotPlot v0.9.0^43^ to visualize sequence identity and its orientation.

***The DJ region*** spans ∼341 kbp upstream of the rDNA arrays in the short arms of the acrocentric chromosomes. We defined the DJ excluding the CER repeat as it was identified in the CenSat annotation from Nurk *et al*.^12^ and Altemose *et al*. 2023^42^. After comparing the DJs on T2T-CHM13v2.0 to each other, we found minor variations on each unit, however the majority stayed concordant (**Figure S2**). The DJs were all similar in terms of length (341,898-346,124 bps) with high identity (>95%), except for the bSat between the DJ arm and DJ flank (**Figure 5C**) and the variation found within the second DJ arm as reported before^44^.

***For the PHR***, we used the region as presented in Guarracino *et al*.^14^ and refined the boundaries to exclude the inverted repeat. The PHR on Chr. 13 is the longest, spanning 1,174,544 bps, while Chr. 14 and Chr. 21 have a shorter form, spanning 1,124,403 and 1,126,601 bps, respectively. We chose the longest form on Chr. 13 as the target for the PHR. The PHR consists of two regions outside the highly repetitive satellite repeats, which we refer to here as PHR arm1 and PHR arm2 (**Figure S3**).

***For the rDNA***, we compared the first rDNA units in each chromosomal array (**Figure S4**). Among them, the unit on Chr. 13 had the longest form and was therefore chosen.

The final target regions for the DJ, PHR arm1 and the first rDNA unit are on Chr. 13. We masked the rest of the units present elsewhere in the reference using bedtools maskfasta (v 2.31.1). Although not used in this study, we additionally masked all 5S rDNA units except the last copy on Chr. 1.

To account for differences in sex chromosome complement, we applied the sex chromosome complement reference (SCC) approach to ensure proper read mapping on the sex chromosomes. Specifically, we used a pseudoautosomal region (PAR)-masked Y reference for XY individuals and a Y-masked reference for XX individuals. **Table S1** contains the exact coordinates of the target and masked region used on T2T-CHM13v2.0.

#### Target regions on GRCh38

To establish target coordinates on GRCh38, we extracted the corresponding target sequences from CHM13 and aligned the GRCh38 reference genome (https://rcs.bu.edu/examples/bioinformatics/gatk/ref/) to each target sequence using Minimap2 (v2.30)^45^ with the -ax asm20 option. The resulting alignments were converted to PAF format using paftools.js (sam2paf). Regions shorter than 10 kbp were excluded from further analysis (**Tables S2-4**).

~~~
minimap2 -ax asm20 {target}.fasta Homo_sapiens_assembly38.fasta.gz
~~~

The PHR regions were defined as described above, resulting in a single location found on chr21:9722154-9796322 (+). Additional inspection was performed to refine the DJ region in GRCh38.

Initially, we evaluated the palindromic region of the DJ (ct_13_9) for counting the DJs. Before applying this approach to the larger cohort, we selected 50 samples from RPC and aligned the reads to GRCh38. Paired-end short reads were aligned to the GRCh38 reference using the same pipeline that we applied to RPC samples aligned to CHM13 (see “Read alignment on each reference” below). Next, we inspected the base-level sequencing depth across the DJ region by lifting over the alignments from GRCh38 to CHM13. This was done using the target and query coordinates in the PAF file, scaling the query’s start and end positions to match the full DJ coordinates.

Per–base-pair depth was calculated using samtools depth. Following manual inspection, a subregion within the original DJ target in CHM13 (chr13:5,566,000–5,600,000) was excluded due to copy number variable alignment behavior in GRCh38. The refined DJ target coordinates were subsequently lifted back to GRCh38 to generate a final BED file for copy number estimation (**Table S2**). We found that using this target produced the expected distribution compared to the samples distribution on T2T-CHM13, with the majority of samples exhibiting approximately 10 DJ copies. This BED file was used to assess DJ copy number variation across UK Biobank samples.

However, we later found that extending the DJ target region to include upstream and downstream of the initial ct_13_9 sequences encompassing the full DJ and its flanking regions resulted in a stronger correlation with the CHM13-based mapping results. Regions on chromosomes 4 and 10 with low sequence identity (<83.4%) and low coverage (<2.8%) were excluded from analysis. Therefore, this expanded region was selected as the final region of interest for DJ copy number estimation.

### Read alignment on each reference

#### CHM13

##### Pilot testing on control samples

We used Illumina short reads from the 3 identified ROB samples from de Lima *et al*^16^, and mapped them to the masked reference. As a control, we used Illumina short reads from CHM13 and HG002, which were previously identified to have 10 DJs from FiSH experimental data (Nurk *et al* and Supplementary Figure XVIII.59 in Yoo *et al*.)^12,46^.

##### RPC

We utilized the read coverage on chromosome X and Y in the original GRCh37 alignments to infer the sample’s biological sex. First, we confirmed the BAM file is sorted based on genomic coordinates. Next, we used samtools idxstats to calculate the number of mapped reads to each sex chromosome and collected the size of the sex chromosomes. The coverage for chromosomes X and Y was determined by dividing the total number of mapped reads by the total chromosome length for each respective chromosome. To infer the sample’s sex, we calculated the coverage ratio between chromosome X and chromosome Y (covX/covY). Samples with a ratio greater than 4 were classified as XX, while those with a ratio of 4 or less were classified as XY. Once the presence of Y was confirmed, the proper masked reference has been set to be used in the downstream analysis.

Short-read paired-end data from both the control samples and the RPC were aligned to the corresponding SCC reference using BWA (v0.7.17)^47^ and post-processed with samtools (v1.21)^48^ using fixmate, sort and markdup. Secondary alignments were removed with samtools view -F0×100 option.

~~~
bwa mem -t $cpu $ref $r1 $r2
samtools fixmate -m -@$cpu $out.sam $out.fix.bam
samtools sort -@$cpu -O bam -o $out.bam -T $out.tmp $tmp/$out.fix.bam
samtools markdup -r -@$cpu $out.bam $out.dedup.bam
~~~

#### GRCh38

To match with the method used in the UKBiobank and 1KGP, the paired-end reads from the six control samples and the 92 subsampled RPC samples were aligned to the reference genome using BWA-MEM (v0.7.17)^47^. The resulting SAM files were processed with SAMBLASTER (v0.1.26)^49^ to add mate tags and sort reads by query name, after which duplicate reads were marked using Picard *MarkDuplicates* (v3.3.0).

### Copy number estimate

#### Mapping based approaches

##### T2T-CHM13v2.0

Copy number estimates for target regions were calculated as the ratio of median sequencing depth in the target region to that of the autosomal background, excluding acrocentric short arms. Median depth for each target and background region was computed using samtools (v1.23)^48^ and bedtools (v2.31.1)^50^ with the following command:

~~~
samtools depth -@ $cpu -b $bed $bam | \
  awk ’{print $1,$2-1,$2,$3}’ OFS=’\t’ -| \
  bedtools map -a $bed -b - -c 4 -o median > $prefix.coverage_results.bed
~~~

For a given target *T*, the copy number (*C_NT_*) was calculated as:

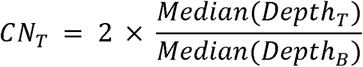

Where *Depth_T_* represents the depth per base-pair within the target region and *Depth_B_* represents the depth across the autosomal region.

##### GRCh38

Challenges in the normalization process arose from collapsed, misassembled, and fragmented target sequences in the acrocentric short arms, in particular the DJ regions in GRCh38, as well as from potential PCR-duplicate–like biases observed in reads mapped to these regions. To reduce computational bottlenecks and minimize the impact on the target copy number estimation, we evaluated four different approaches for calculating the target and background sequencing depth.

Starting from the pilot-Fast mode, we developed four approaches and benchmarked the 92 RPC samples and compared correlations of the distribution to the result from T2T-CHM13 (**Figure S7**).

***For the pilot Fast mode applied to the UK Biobank cohort,*** we used samtools (v1.20)^48^ to collect the number of reads mapped to the target and all regions. Specifically, we used `samtools view -c` aligned to the target (*RC_T_*, **Table S2** for the DJ target) and `samtools idxstats` to collect reads aligned to the autosomes by using num. of mapped reads − num. of unmapped reads. In this mode, no reads are filtered prior to calculation. Unlike the RPC samples, we learned `samtools depth` was creating a bottleneck and was too costly to run on all 490k samples on DNAnexus (1 hour vs. 2 minutes using samtools idxstats). Albeit a little shifted, we found a good correlation to the T2T-CHM13 based result for the 92 subsamples (Pearson’s correlation, r=0.981, p=57×10^-66^) between the two methods (**Figure S8**), and decided to move forward using this approach. The script was deployed on DNAnexus and collected copy counts of the DJ, rDNA, and PHR.

Copy number for each target was estimated using read count normalization. Read counts (*RC*) mapped to each target were normalized to autosomal coverage, which was calculated by subtracting unmapped read counts (*RU_A_*) from the total reads mapped to autosomes (*RC_A_*) and dividing by the total autosomal length (*Len_A_*). The normalized target RC was then used to infer copy number.

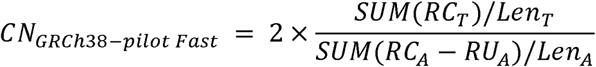

After a series of evaluations, we updated our method to include the DJ flanking region in the target as it yielded better copy number estimates. The following three modes (**Table S5**) use the target DJ region as shown in **Table S3**.

***The Fast mode*** estimates copy number using the number of reads aligned to the target region (*RC_T_*) calculated by `samtools view -c`, normalized by the original target length in CHM13v2.0 (*Len_T_*). Autosomal coverage is calculated by using the median of autosomal read coverage, defined as the total number of reads mapped to each autosome (*Len_C_*) divided by the length of the autosomal (*RC_C_*) reported by using `samtools idxstat`. In this mode, no reads are filtered prior to calculation.

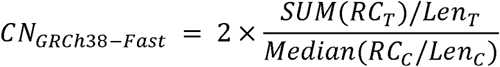

***In the Fast-precise mode,*** reads marked as duplicates, as well as secondary and supplementary alignments, were excluded from analysis. Target coverage was calculated as the total filtered read counts aligned to the target region (*FRC_T_*) (as reported by `samtools view -c -F 3332`) divided by the target length (*FRC_A_*). Background coverage was calculated as the total number of filtered reads aligned to the autosomes (*Len_T_*) divided by the total autosomal length (*Len_A_*).

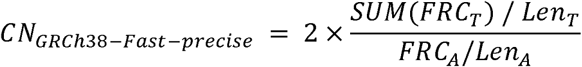

***The High-resolution mode*** calculates coverage at per-base resolution, consistent with the approach used for CHM13v2.0. The depth for each target region was computed using `samtools depth` (*Depth_r_*) and normalized by the target length (*Len_T_*). For background coverage, we used `samtools coverage` to obtain coverage metrics for each autosome (*Cov_c_*) and defined the background as the median autosomal coverage.

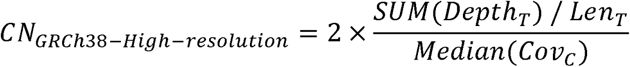

#### Reference-free k-mer based approach

***For identifying the DJ k-mers,*** 31-mers were collected with Meryl (v1.4.2)^34^ for each of the five DJ copies in T2T-CHM13v2.0. The DJ sequence was extracted with `samtools faidx`, and `meryl count k=31` was used to build the `dj.$chr.meryl` for each chromosome. Only *k*-mers present once in each chromosome were selected and intersected, to ensure we use *k*-mers present once in each DJ and in all 5 DJs. This yielded 178,731 potential target *k*-mers in `dj.meryl`.

~~~
meryl intersect [equal-to 1 dj.chr13.meryl] [equal-to 1 dj.chr14.meryl]
output dj.chr13_chr14.1.meryl
meryl intersect [equal-to 1 dj.chr15.meryl] [equal-to 1 dj.chr21.meryl]
output dj.chr15_chr21.1.meryl
meryl intersect [intersect dj.chr13_chr14.1.meryl dj.chr15_chr21.1.meryl]
[equal-to 1 dj.chr22.meryl] output dj.meryl
~~~

***For identifying the DJ target k-mers,*** the DJ *k*-mers were filtered one more time after projecting the k-mer multiplicity as found in the short reads. The 31-mer meryl DB was built with `meryl count k=31`, and the histogram was generated with `meryl histogram`. The 2 copy peak was inferred using GenomeScope 2.0^51^ with the following command line:

~~~
Rscript $tools/genomescope2.0/genomescope.R -i chm13.k31.hist \
~~~

-o chm13.k31.genomescope2 --fitted_hist --testing -k 31

The predicted haploid coverage (`kcov`) resulted in 49.9. The inspected 10 copy peak in CHM13 is therefore ∼50 x 10; resulting in ∼500x. This matched the peak distribution of the DJ *k*-mers when projected to the read spectrum (**Figure 1E**). The final selected DJ target *k*-mers were chosen falling between the range of 475-525x (±0.5 copy), to allow room for small variations or sequencing biases. This resulted in 52,227 distinct DJ target *k*-mers in `dj.target.meryl`.

~~~
meryl greater-than 475 [less-than 525 dj.chm13_count.meryl] \ output dj.target.meryl
~~~

***For copy number estimating the DJs in a given sample,*** we first collect 31-mers from the samples’ short reads. The 31-mer database is then queried to collect the k-mer multiplicity histogram of the DJ target *k*-mers. The median of the DJ target k-mers found in the reads is used as the *k*-mer frequency of the DJs, and the “2-copy peak” is inferred from the histogram of all 31-mers in the reads. The 2-copy peak is obtained using a script in Merqury (`kmerHistToPloidyDepth.jar`), which differentiates the slope of the *k*-mer histogram and reports the estimated ploidy and its peak and boundary by collecting the local maxima^34^.

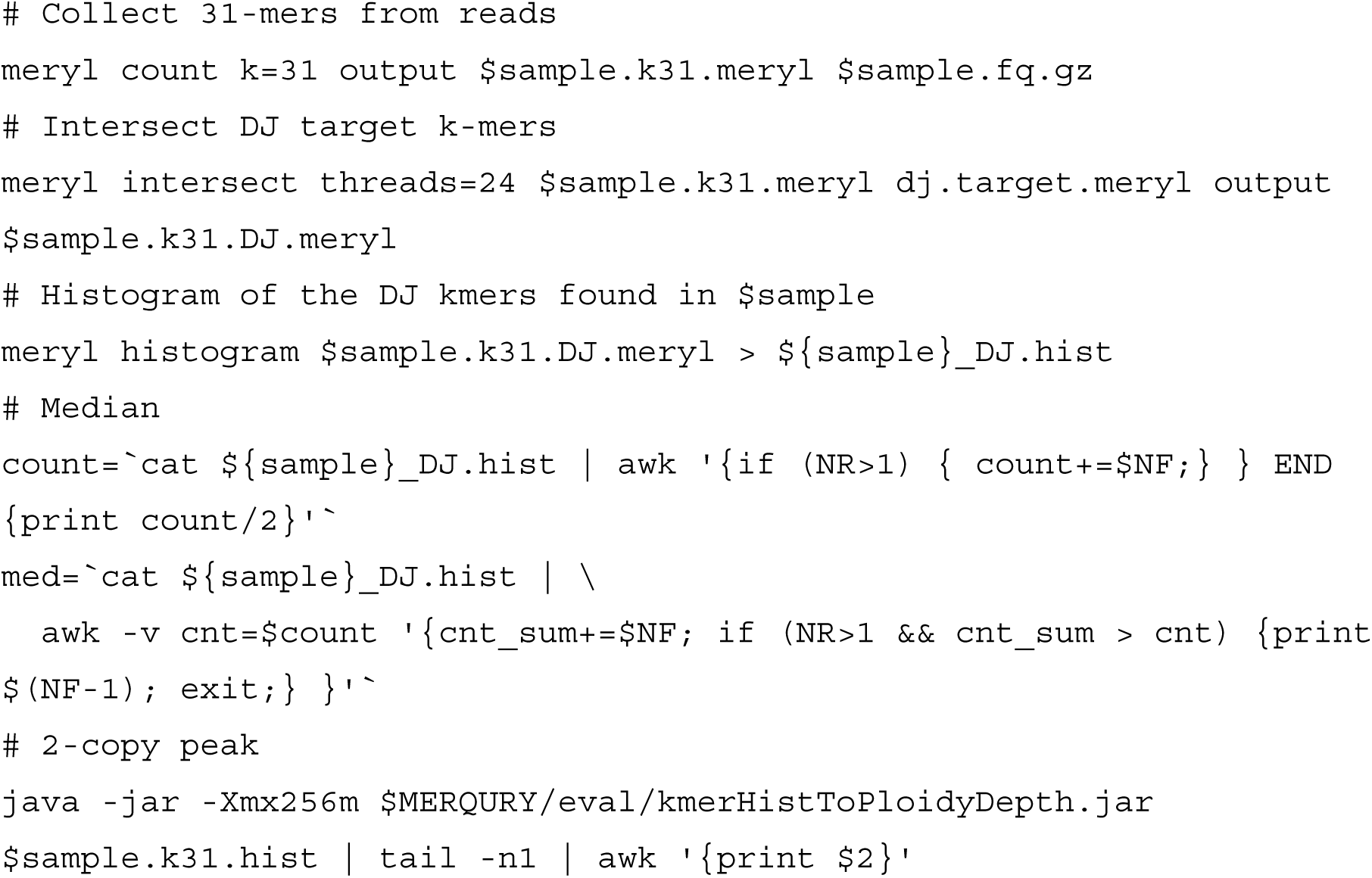

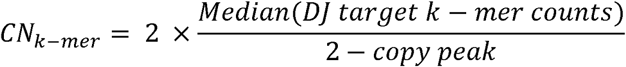

### Binning into counts

After the estimated copy numbers were collected for all samples, the distribution was manually inspected to discretize and bin the counts for further comparison in **Tables 2** and **S6**. In most cases, a simple rounding (cut-off at 0.5) matched the distribution dips for the CHM13 reference based and the k-mer based methods. The GRCh38 based counts however resulted in a shifted distribution, and therefore was manually adjusted to cut the binning boundaries to match the dip in the distribution. This motivated further development of a more sophisticated binning method for GRCh38 mapping based analysis, as described in the “GMM fitting and classification” section.

#### Chromosome spreads and Fluorescent In-Situ Hybridization (FISH) for rDNA and DJ

Human lymphoblastoid cell lines (LCL) and fibroblast cell lines were according to Coriell protocols. CHM13 immortalized using human telomerase reverse transcriptase (hTERT) were propagated in DMEM:F12 with L-glutamine supplemented with 1x non-essential amino acids (Gibco), 1x Insulin-Transferrin-Selenium (Gibco), 1mM sodium pyruvate (Gibco), and 10% FBS. For preparation of chromosome spreads, cultures were blocked in mitosis by the addition of Karyomax colcemid solution (0.1 µg/ml, Thermo Fisher) for 6-7h. Adherent fibroblast cells were collected by trypsinization. Cells were incubated in hypotonic 0.4% KCl solution for 12 min and prefixed by addition of methanol:acetic acid (3:1) fixative solution (1% total volume). Pre-fixed cells were spun down and then fixed in Methanol:Acetic acid (3:1). Chromosome spreads were dropped on a glass slide and incubated at 65°C overnight. Before hybridization, slides were treated with 0.1mg/ml RNAse A (Qiagen) in 2xSSC for 45 minutes at 37°C and dehydrated in a 70%, 80%, and 100% ethanol series for 2 minutes each. Slides were denatured in 70% deionized formamide/2X SSC solution pre-heated to 72°C for 1.2-1.5 min. Denaturation was stopped by immersing slides in 70%, 80%, and 100% ethanol series chilled to −20°C. Probes used in combinations were denatured in a hybridization buffer (Empire genomics) by heating to 80°C for 7 minutes before applying to denatured slides. Specimens were hybridized to the probes under a glass coverslip or HybriSlip hybridization cover (GRACE Biolabs) sealed with the rubber cement or Cytobond (SciGene) in a humidified chamber at 37°C for 48-72hours. After hybridization, slides were washed in 50% formamide/2X SSC 3 times for 5 minutes per wash at 45°C, then in 1x SSC solution at 45°C for 5 minutes twice and at room temperature once. For biotin detection, slides were incubated with streptavidin conjugated to Cy5 (Thermo) for 2-3 hours in PBS containing 0.1% Triton X-100 and 5% bovine serum albumin (BSA), and then washed 3 times for 5 minutes with PBS/0.1% Triton X-100. Slides were rinsed in water, air-dried in the dark, and mounted in Vectashield containing DAPI (Vector Laboratories). Z-stack confocal images were acquired on the Nikon TiE microscope equipped with 100x objective NA 1.45, Yokogawa CSU-W1 spinning disk, and Flash 4.0 sCMOS camera (Hamamatsu). Image processing was performed in FIJI.

### Pattern of DJs in the HPRC samples

#### HPRC Verkko assemblies

In parallel to the HPRCr2 assemblies, a Verkko v2.2.1^31^ assembly of each genome was made using automated scripts to download data from AWS, build parental k-mer databases, and run Verkko with both Hi-C and Trio data in parallel. Standardized post-processing and QC was performed with the same pipeline which used Yak (https://github.com/lh3/yak) to measure QV (k=31) and switch error rate, compleasm^52^ for gene completeness, and computed T2T contig and scaffold statistics using seqtk to identify the telomeric and gap sequences in the assemblies. For each sample where a trio and Hi-C was available, the assembly with the most T2T contigs was selected. In total, there were 59 trios and 159 Hi-C assemblies.

#### Chromosome assignment and orientation

Mashmap v3.1.1^53^ was used to map the assembly to CHM13v2.0. The contigs were assigned to chromosomes if they had mappings over 99% identity covering more than 10% of the reference chromosome in a given orientation (forward or reverse-complement). If a contig matched over 10% of more than one chromosome, multiple assignments were noted and the longest one was used. The XY pseudo-autosomal regions (PAR) and distal regions of the acrocentric were excluded from the covered fraction count as they recombine between individuals and cannot be accurately assigned via reference alignment. Therefore, these regions could only be named as a chromosome if they were assembled together with a unique region of the chromosome. Across all samples and assembly types (verkko-hi-c or verkko-trio+hi-c), there were 23,094 contigs totaling 2.11 Tbp with an average size of 91.58 Mbp which could be oriented and assigned. There were 338 contigs with ambiguous assignments with an average size of 5.7 Mbp and a total of 1.936 Gbp while 75,565 contigs could not be assigned with an average size of 0.12 Mbp totaling 8.73 Gbp.

In parallel, the homopolymer-compressed Verkko gfa were aligned using Mashmap (v3.1.1)^53^ to a compressed version of CHM13v2.0 to assign the individual nodes to a chromosome. A two pass assignment was performed. First, nodes with mappings over 99% identity and 5 Mbp matches to a chromosome were assigned to that chromosome. Second, if a connected component had no nodes assigned by the first criteria, nodes over 500 kbp were assigned. This identified connected components in the assembly graph belonging to each chromosome.

#### Repeat annotation of the acrocentric chromosomes

The acrocentric chromosomal short arms carry human satellite repeats (HSat) at and around the centromere, some specific to the acrocentric region forming a particular pattern^12,54^. To obtain an overview of the sequence content and confirm sequences belonging to the acrocentric short arms, particularly the distal and proximal sequences from the rDNA, satellite sequences in the assemblies were annotated based on a set of targeted sequence (Table S8).

In brief, the target repeat sequences were selected based on the CenSatv2.1 annotation available for T2T-CHM13v2.0 (https://s3-us-west-2.amazonaws.com/human-pangenomics/T2T/CHM13/assemblies/annotation/chm13v2.0_censat_v2.1.bed). Most were selected from chr13 or chrY, except for HSat1A, HSat3_A5, and HSat3_B3. For these three HSat repeats, the original consensus generated for classifying the satellite array subgroups were used as described by Altemose *et al*.^55^. The consensus sequences are available as fasta files on https://github.com/altemose/HSatReview/tree/main. The rDNA reference KY962518 was rotated to begin upstream of the 45S TSS, to include the 45S promoter region.

The target sequences were found using minimap2 (v2.28) in each assembly. Because most mappers are designed to find one best match for each query in the reference, using the assembly as the reference would yield only one result for each mapped target satellite. Thus, the reference and query sequences were flipped, mapping the assembly (query) to the targeted repeat sequence (reference) instead. This approach allows a more sensitive search of the target sequence in the assembly. Once all the alignments are collected in a PAF file, the reference and query coordinates were inverted with rustybam (v0.1.33)^56^ and converted to BED format. Alignment blocks within 500 bp were merged with BEDTools (v2.31.1)^50^ and formatted with designated colors and filtered to be over 2 kbp or 7–8 kbp to exclude excessive alignment matches to LINE elements.

Below are the command lines used:

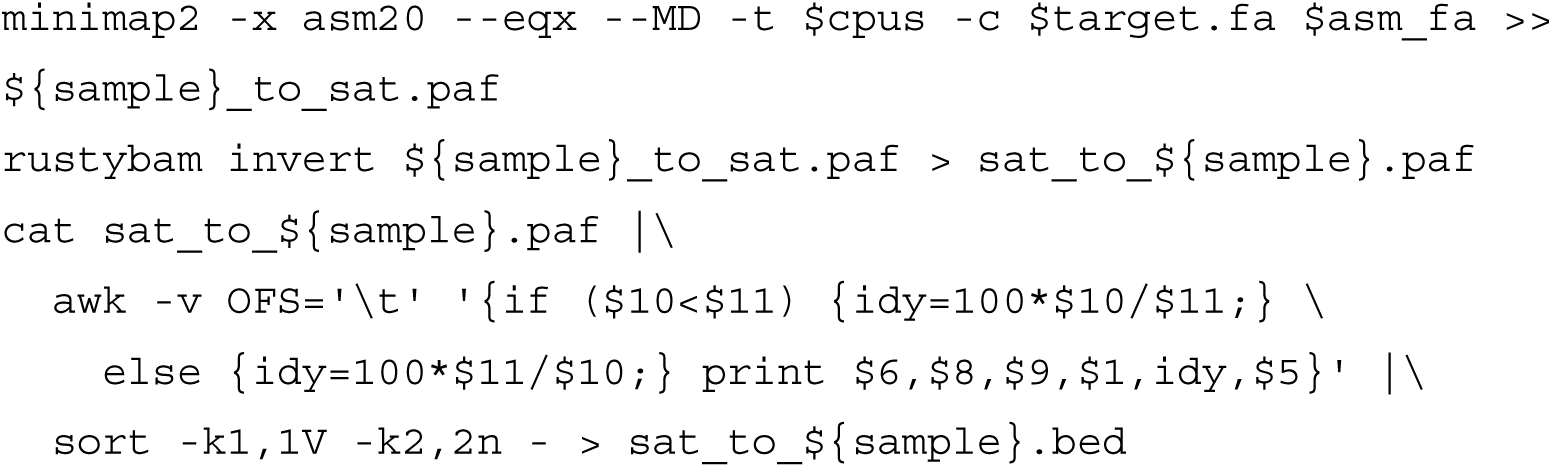

For each target satellite, merging and filtering was applied along with the color assignment. Telomere sequences and gaps were annotated with Seqtk (v1.4, https://github.com/lh3/seqtk) using tel -d5000 and gap -l 1 parameters.

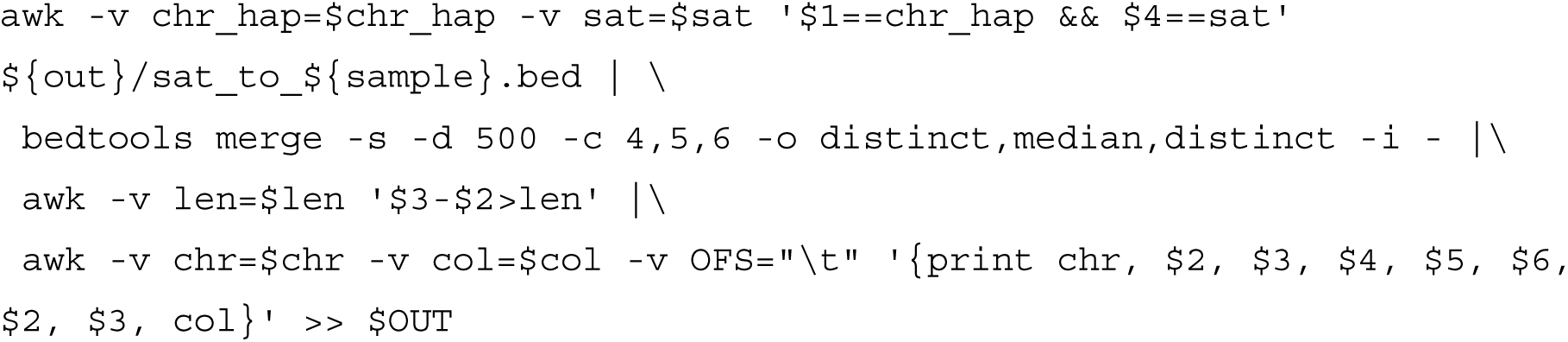

The final BED file was then concatenated and sorted for manual inspection and visualization on IGV (v2.18.2)^57^.

### Benchmarking and optimizing DJCounter for GRCh38

#### Subsampling and benchmarking computing resources

HG002 reads aligned to GRCh38 were subsampled to the desired sequencing depths (1×, 5×, 10×, 20×, 30×, 40×, and 50×) using samtools (v1.21)^48^ to evaluate the stability and computational requirements of each method. We ran DJCounter on the subsampled BAM files using the GRCh38-based modes and additionally extracted reads from the BAM files for analysis with the k-mer–based method.

To benchmark computational requirements of each DJ estimation mode, we extracted runtime and resource utilization metrics from SLURM job accounting records using sacct. For each job, we summarized: (i) Elapsed time as the maximum wall-clock duration across steps (hours); (ii) Total CPU time as the sum of CPU time across steps (hours); (iii) maxMem as the maximum resident memory usage (MaxRSS, GB); (iv) maxVmem as the maximum virtual memory usage (MaxVMSize, GB); (v) %CPU as the average CPU utilization relative to a single fully utilized core (100%); and (vi) AvgCPU as the mean CPU time per step (hours).

#### Linear transformation between GRCh38- and CHM13-derived DJ copy numbers

To determine which GRCh38 mapping mode exhibited the least cohort bias and to harmonize DJ copy number estimates derived from GRCh38 alignments with those obtained from CHM13, we fitted an ordinary least squares (OLS) linear regression model using the *statsmodels* package (v0.14.5)^58^. Specifically, DJ copy numbers calculated from CHM13 alignments of 92 RPC subsamples were modeled as a function of DJ copy numbers derived from GRCh38 alignments of the matched samples:

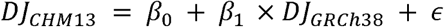

where *β_0_* represents the intercept, *β_1_* the slope, and *∈* the residual error term.

We fitted this model separately using DJ copy numbers generated from different GRCh38 modes (Fast, Fast-precise and High-resolution) and selected the Fast-precise mode because it yielded the lowest residual sum of squares (RSS) among all modes and showed no significant difference between the CHM13-aligned RPC DJ distribution and the transformed 1KGP DJ distribution.

To harmonize DJ copy numbers across cohorts, we applied the fitted Fast-precise regression model to transform GRCh38-derived DJ estimates to the CHM13 scale using the ‘predict’ function in *statsmodels*. The transformed values were subsequently used for GMM fitting and classification analyses

#### GMM fitting and classification

We used the transformed DJ copy numbers to split the 1KGP samples into a training set and a test set at a ratio of 0.75 and 0.25, respectively. When applying a standard GMM without fixed means, the model failed to identify peaks in sparsely populated regions and exhibited over-dispersion or component collapse, particularly for rare categories such as ROB samples with 8 copies. To address this issue, we performed GMM fitting with fixed component means and constrained standard deviations (0.1≤ σ ≤ 0.2).

Candidate peak locations were identified from the DJ copy number distributions of the RPC cohort aligned to CHM13 and the transformed 1KGP (GRCh38-aligned) cohort using the ‘find_peak’ function from the *scipy.signal* module (v1.13.1)^59^. Peaks that were consistent across cohorts were selected as 7(*k*) fixed component means (*u*): [8, 9.1, 10.1, 10.7, 11.1, 11.9, 13].

To quantify uncertainty in the estimated mixture parameters, we performed nonparametric bootstrap resampling on the training set. Using the DJ copy numbers from the training data, we generated B=1,000 bootstrap datasets by sampling observations with replacement. For each bootstrap replicate b, we fit a one-dimensional GMM with fixed means and constrained standard deviations using an expectation–maximization (EM) algorithm. For each replicate, we estimated the mixture weights (*ψ_kb_*) and standard deviations (σ_kb_) for each k. Model fitting was performed with a maximum of 500 iterations, and convergence was defined as a change in log-likelihood of less than 1×10e-10 times between successive iterations. Final parameter (*ψ_k_* and σ_k_) estimates were obtained by averaging the bootstrap estimates of weights and standard deviations.

For classification, we used the fitted GMM parameters to assign samples from the test set to discrete DJ copy number categories. For each sample, the unnormalized posterior density for each component was computed as *ψ_k_. norm.pdf (x_i_,u_k_,σ_k_)* and normalized across components to obtain posterior membership probabilities. Each sample was assigned to the category with the highest posterior probability (maximum a posteriori classification).

### False positive and false negative rate

#### False positive rate

We assume ROB carriers have 8 DJ copies based on observations from prior studies. However, as shown in our study, we found 1 DJ-less haplotypes (9 DJs) across different cohorts at a frequency of 3.1%. Assuming the 10 acrocentric chromosomes are independent, this gives us a rough estimate that each acrocentric chromosome can lack a DJ with probability of 1 - (1 - 0.031)^1/10^ ≈ 0.31%. As such, the probability of a deletion of DJs is 0.0031, the probability that exactly 2 of them have a deletion is given by

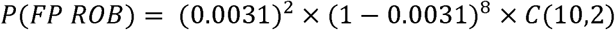

This corresponds to approximately 0.04% of all samples. This could consist up to ⅓ of detected 0.12% 8 DJ cases.

#### False negative rate

Given the high frequency we observed for one or more DJ gains (∼8.9%), it is also possible that a ROB carrier can be misclassified as having 9 copies when one of the non-fused acrocentric haplotype has the duplicated DJ present. Similarly as above, we can roughly estimate the probability for having a duplicated DJ in each acrocentric chromosome as 1 - (1 - 0.089)^1/10^ ≈ 0.0093. Thus, the chance that a ROB carrier will have duplicated DJ in at least one of the other acrocentric chromosomes and thus more than 8 DJ can be estimated as

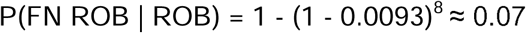

As a result, we expect that roughly 7% of ROBs can be missed by using 8 DJs as the sole marker for identifying ROB carriers.

Copy number analysis alone cannot distinguish between a true ROB carrier and this alternative scenario. To determine whether samples with 8 copies carry a genuine fusion chromosome, additional analysis such as karyotyping or Hi-C are required. Given that we did not observe any offspring with both parents having 9 DJs in the UK Biobank dataset, we estimate the chances might be even lower. We believe a more dedicated study on familial participants with identified 8 DJ offspring and 9 DJ parents may answer this question.

