## Supplemental Figures and Tables for "Biobank-scale genotyping of Robertsonian translocations reveals hidden structural variation on the human acrocentric chromosomes"

### Supplementary Figures

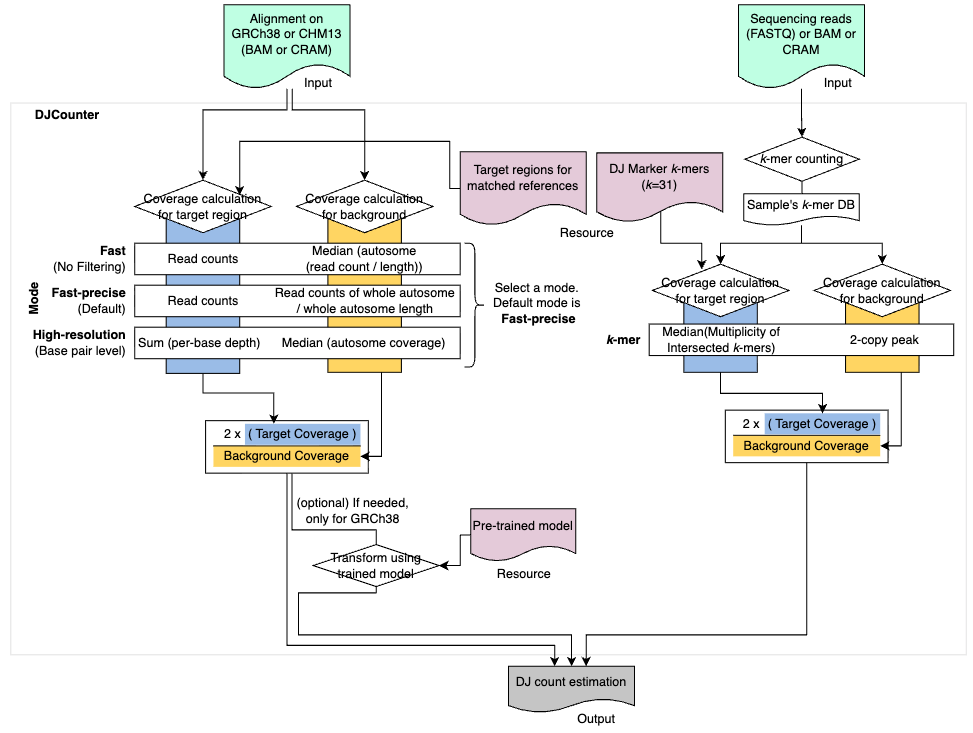

##### Figure S1. Workflow of DJCounter, related to Figure 1 and 5I.

Inputs are indicated in green and consist of either alignment files (BAM or CRAM aligned on CHM13 or GRCh38) or sequencing read files (FASTQ). Several resources are provided within DJCounter as shown in pink in the center of the workflow. On the left, different modes for estimating DJ counts from alignment files are shown. These modes use different approaches to calculate coverage, with or without duplicate-read filtering, and can either count reads or calculate base-pair depth and coverage. On the right, the k-mer mode is illustrated. The first step is to build a k-mer database for the sample using meryl and k-mer size of 31. This database is then used to identify the two-copy peak to use as background coverage and the median multiplicity of the intersecting k-mers between the sample k-mers and the pre-built k-mers from the DJ region as target coverage. For the alignment-based approach using GRCh38, with small sample size (<1000), we recommend transforming the DJ count using a pre-trained model to adjust it to a CHM13-equivalent count. The output is a DJ count (float) for the sample.

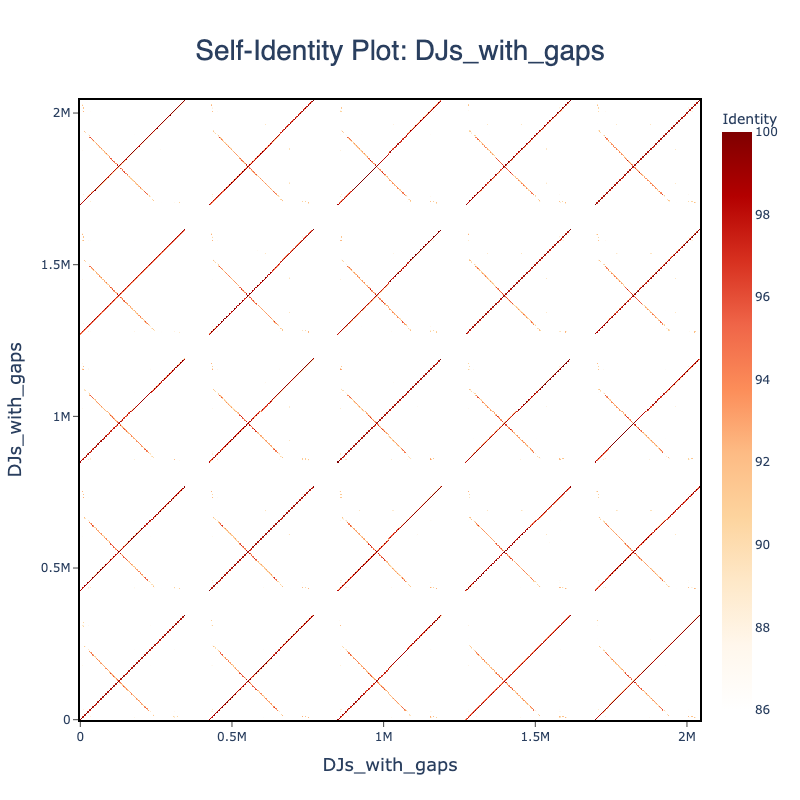
A

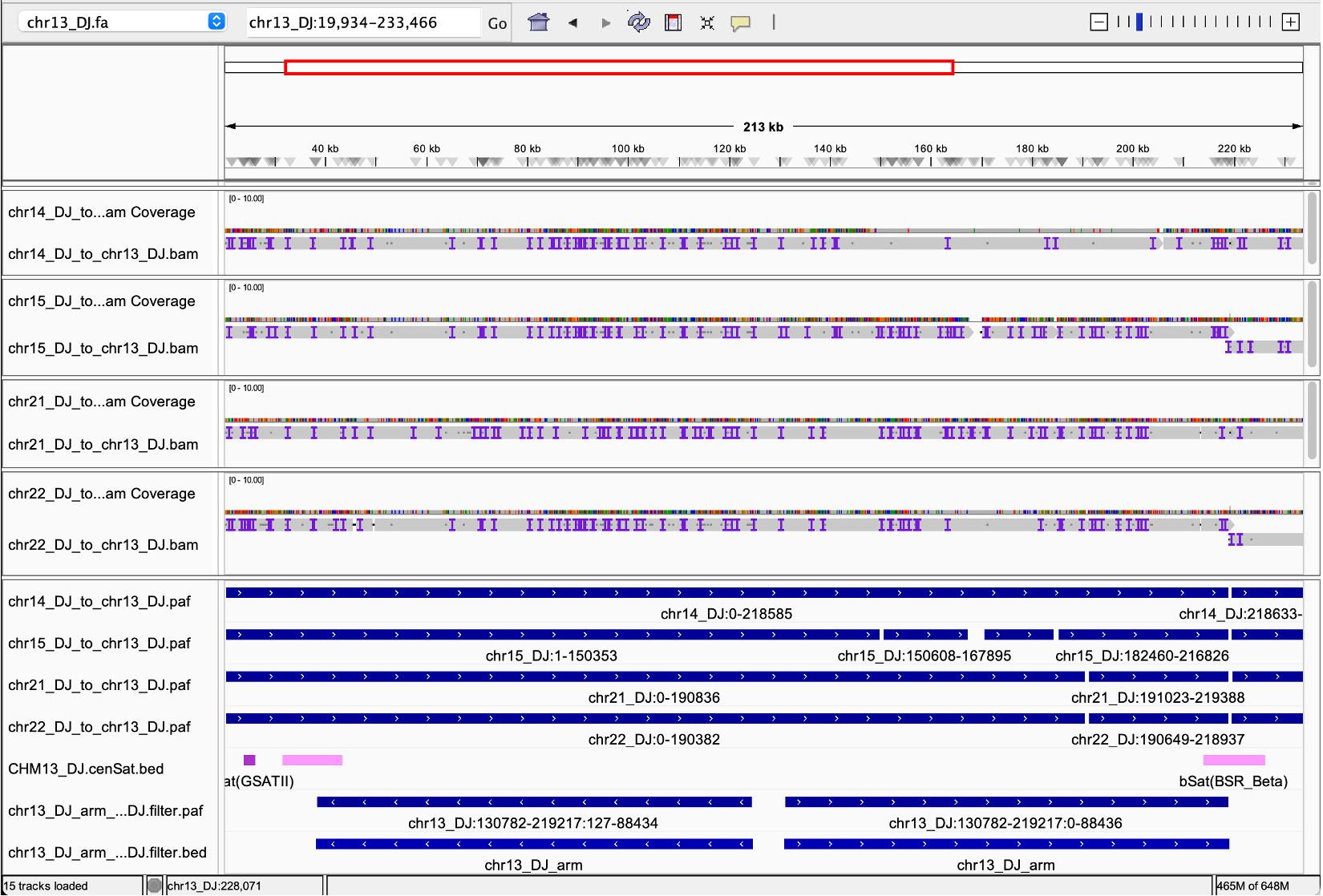
B

##### Figure S2. Target DJ region in CHM13, related to Figure 1B.

(A) Self-identity dotplot of the five DJs drawn with ModDotplot v0.9.0. All DJ units on the five acrocentric chromosomes were compared through a self-identity plot. Each DJ unit was separated by an 80 kbp gap for visualization purposes. The DJs are ordered from Chr. 13 to Chr. 22.

(B) Variations in each DJ unit are enriched at the bSat on the right side, between the DJ arm and DJ flank, and the second DJ arm (right side).

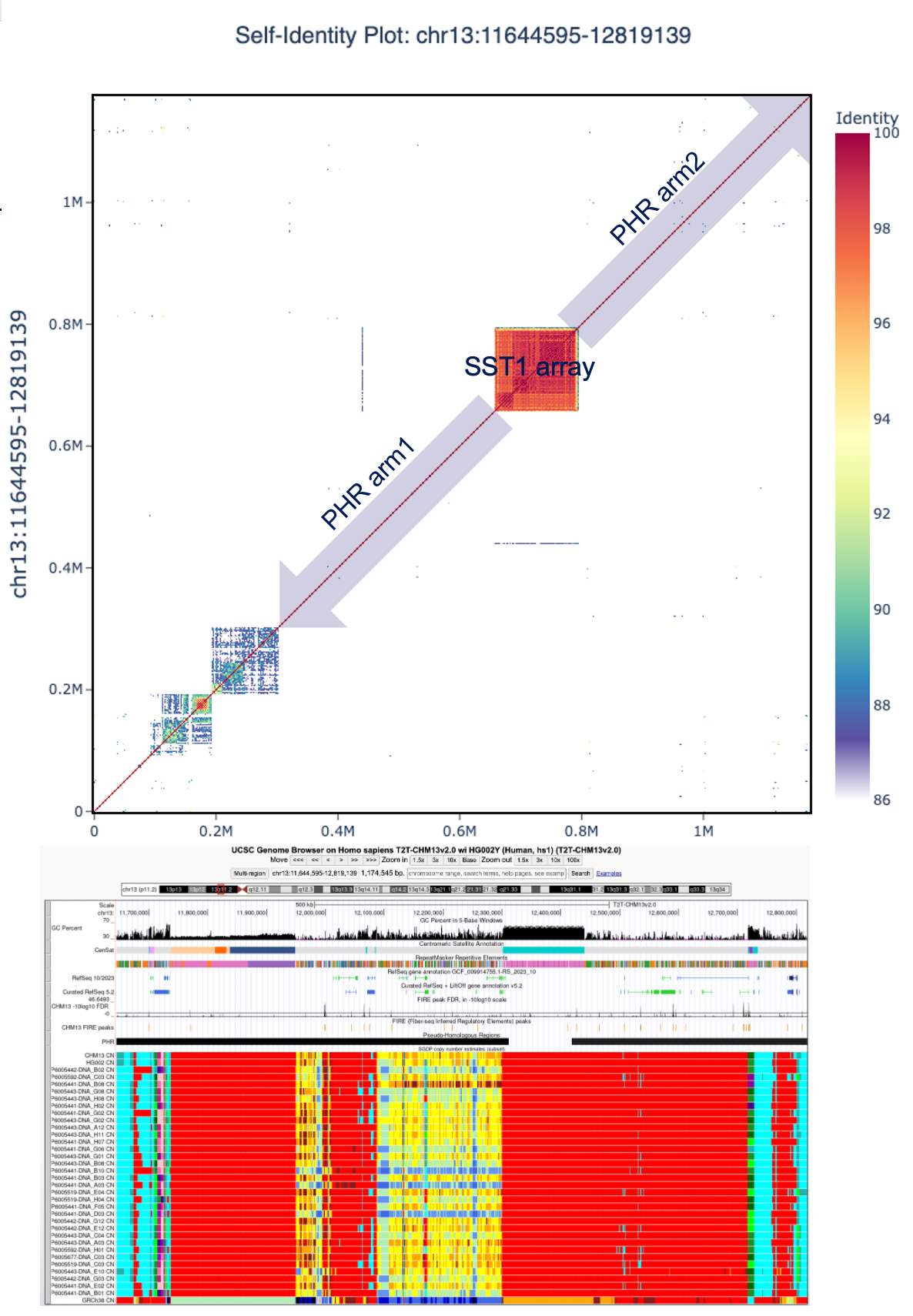

##### Figure S3. Target PHR region in CHM13, related to Figure 1C.

The target was chosen for the longest form as present on Chr. 13. The PHR showed inconsistent copy numbers from the SGDP copy number variation panel, thus we tried 3 regions: the entire PHR, PHR arm1 and PHR arm2 as shown in the plot. After evaluating the distributions, we chose PHR arm1 as the target for PHR.

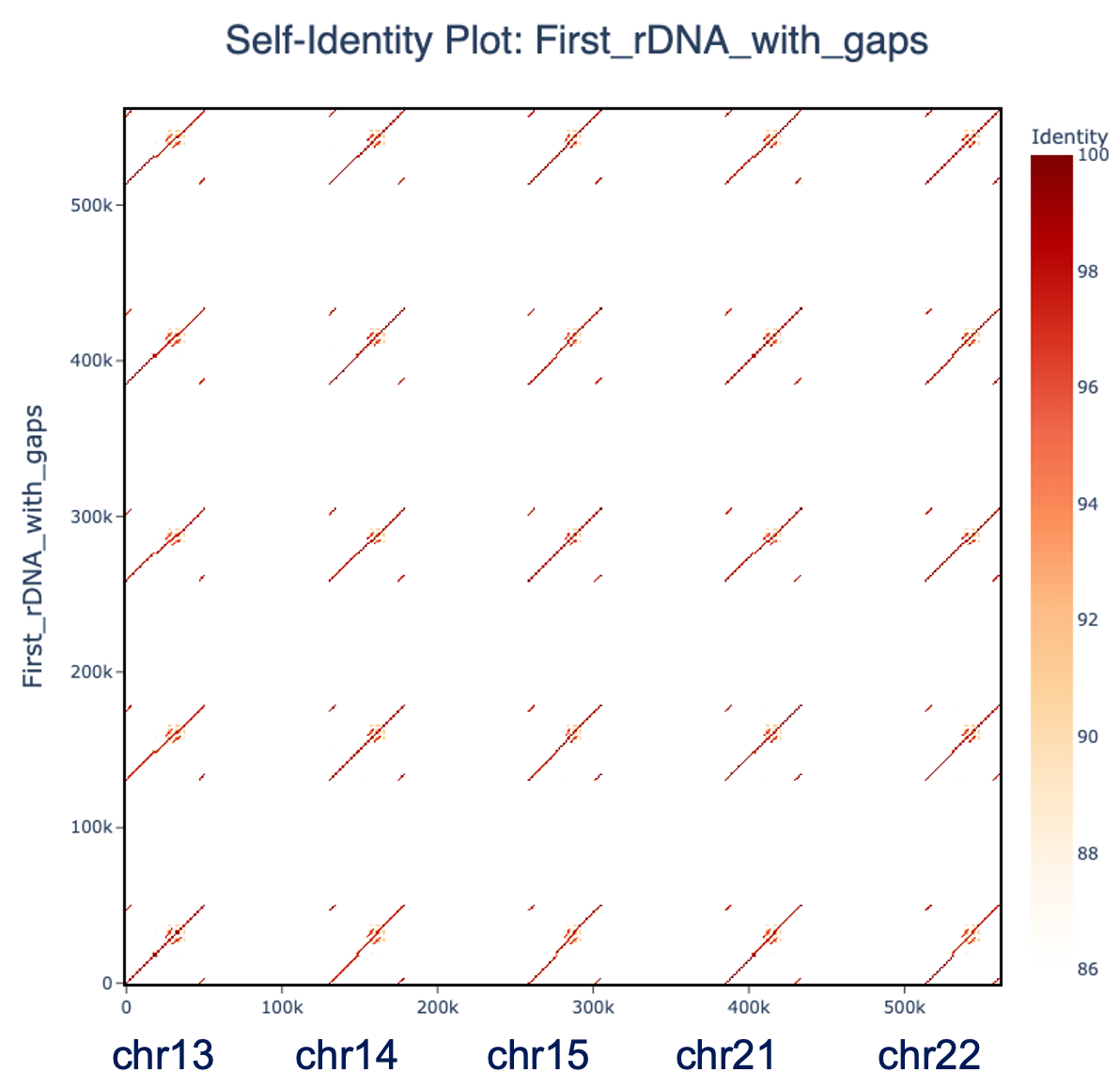

##### Figure S4. Self-Identity plot for the first units of the rDNAs in each chromosome, related to Figure 1C.

The unit in Chr. 13 had the longest form of rDNA.

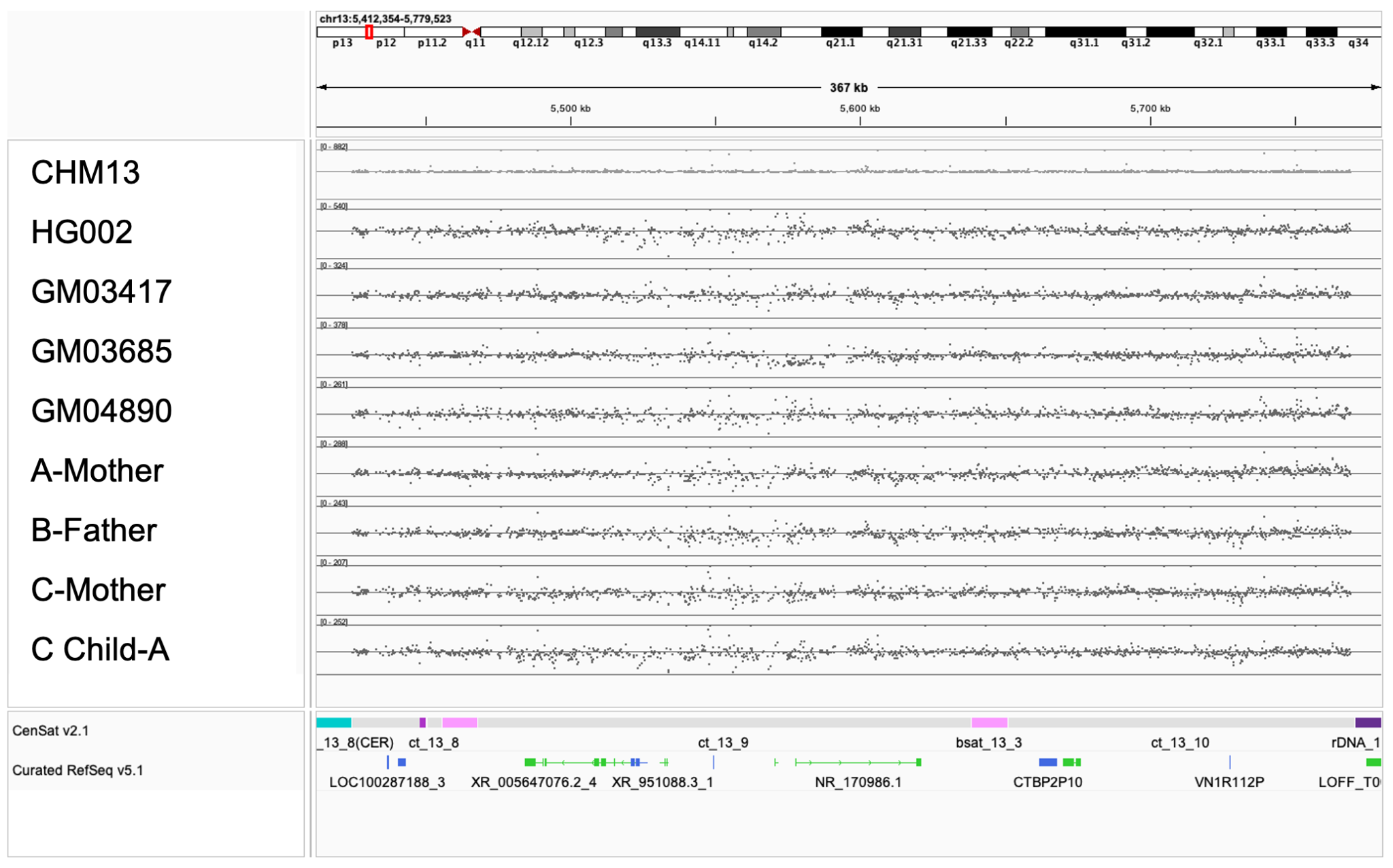

##### Figure S5. DJ target k-mers in the control samples and the 4 predicted ROB samples found in the RPC, related to Figure 1H.

The k-mer multiplicity is shown as found in the read set on the position of the k-mer as it appears on CHM13 Chr. 13 DJ region. The line across indicates the median of the DJ target k-mer multiplicity. It matched the estimated 10 copies for CHM13 and HG002 and 8 copies for the ROB samples based on the 2 copy peak observed in the k-mer histogram.

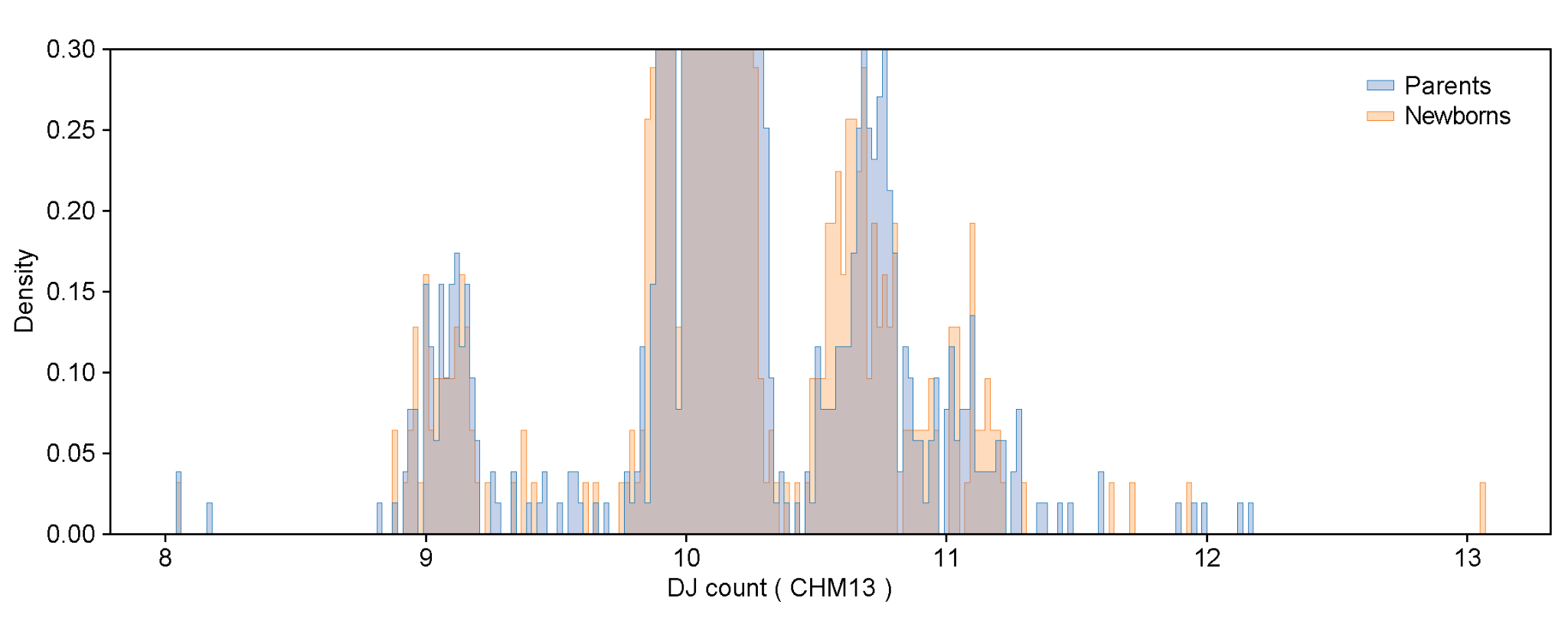

##### Figure S6. Zoomed-in density distribution of RPC samples: parents vs. newborns; related to Figure 2B.

This panel shows a magnified view of the top panel in Figure 2B.

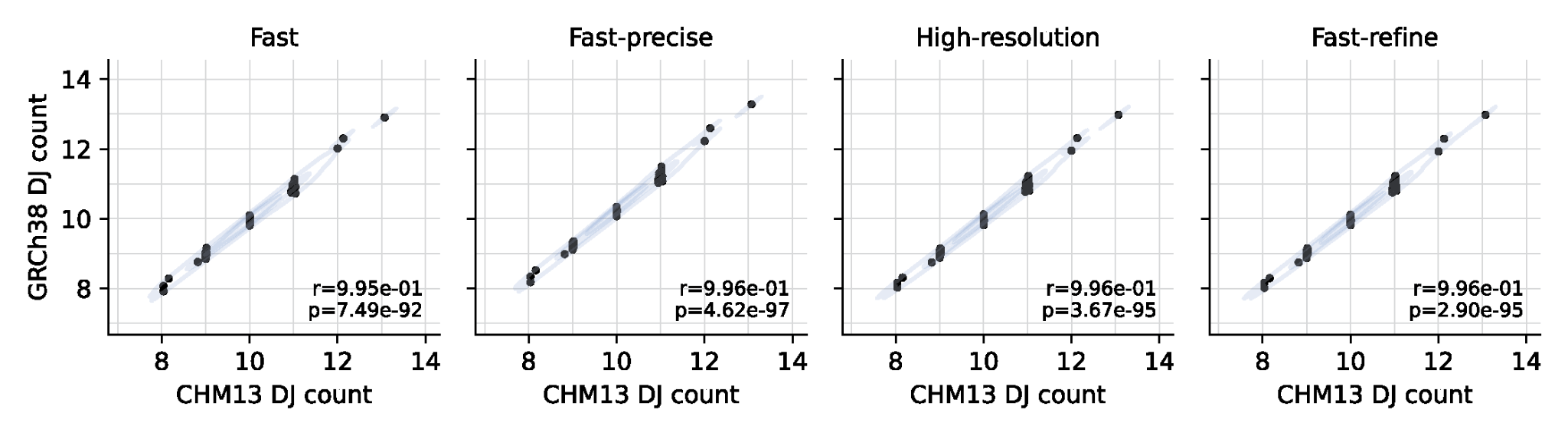

##### Figure S7. Correlation of DJ copy number estimates between GRCh38 mapping-based methods and the CHM13 mapping-based method; related to Figure 2D.

Pearson correlation analysis was performed between DJ copy number estimates derived from four GRCh38 mapping-based methods and those obtained using the CHM13 mapping-based method.

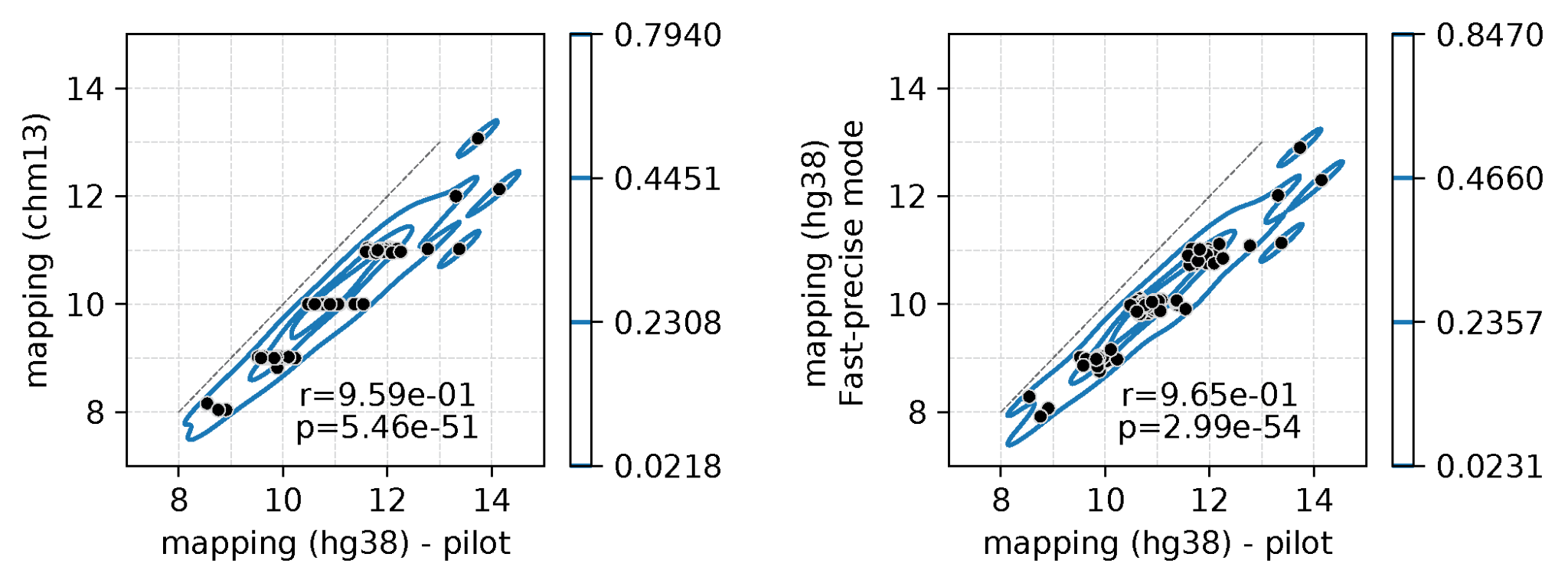

##### Figure S8. Comparison of mapping approaches using 92 subsampled RPC samples, related to Figure 3A.

Scatter plots comparing pilot-Fast mode with CHM13 mapping, and Fast mode with pilot-Fast mode. Correlations were estimated using Pearson’s correlation coefficient.

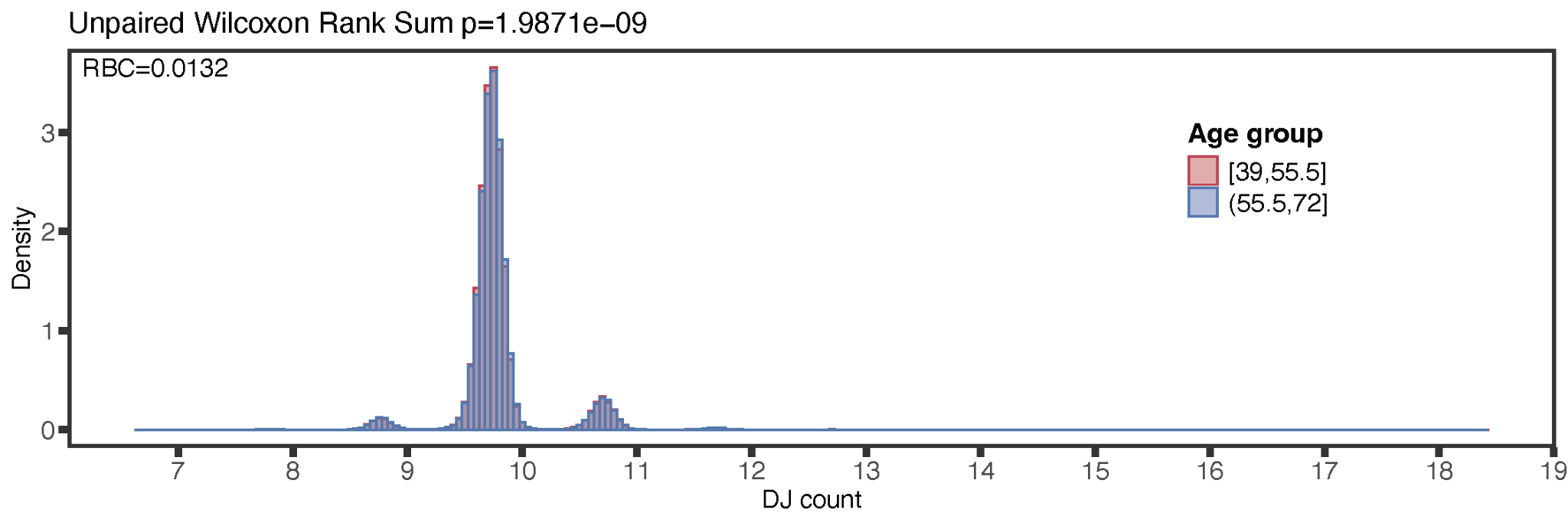

##### Figure S9. Density plot of DJ by two age groups in the UK Biobank, related to Figure 3B.

The two groups were split at 55.5 years. The results of the unpaired Wilcoxon Rank Sum test are shown. In the RBC (p = 0.0132), the older group tended to have higher DJ counts than the younger group.

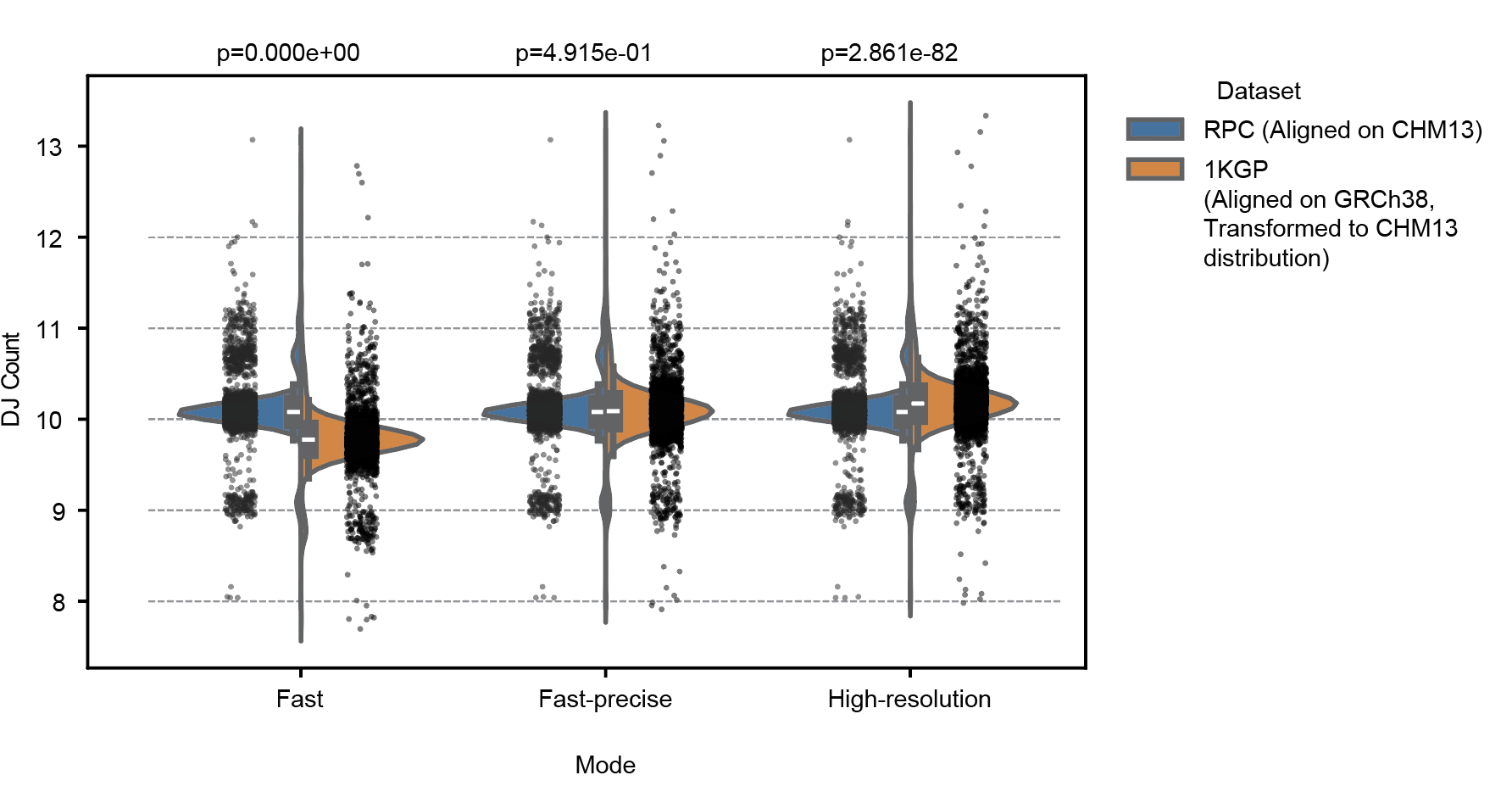

##### Figure S10. Comparison of DJ copy number distributions between the RPC cohort aligned to CHM13 and the transformed DJ copy numbers from the 1KGP cohort using the corresponding model, related to Figure 4A.

P-values were calculated using the Mann–Whitney U test.

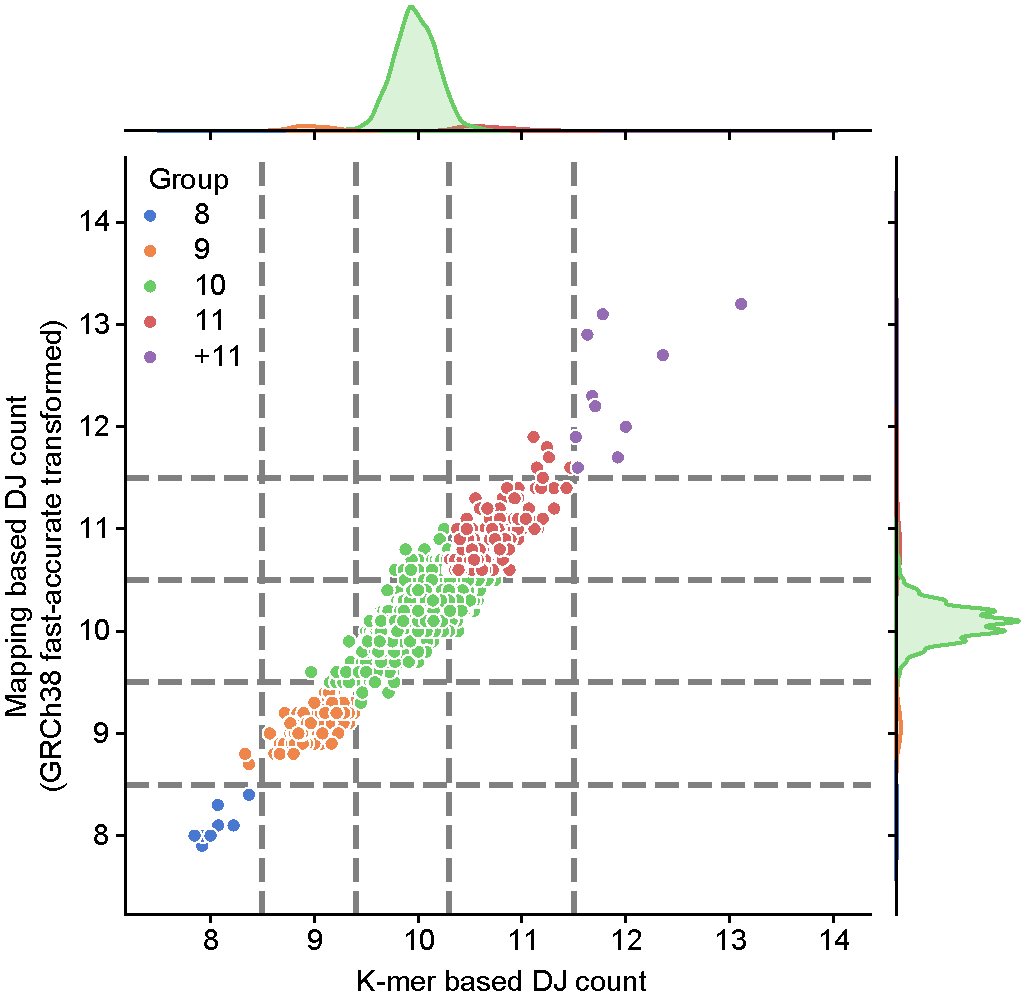

##### Figure S11. Grouping of 1KGP samples by DJ values, related to Figure 4B.

Scatter plot showing DJ values for 1KGP samples calculated using the *k*-mer-based method (x-axis) and the GRCh38 Fast-precise mode (transformed; y-axis). Samples are colored according to group.

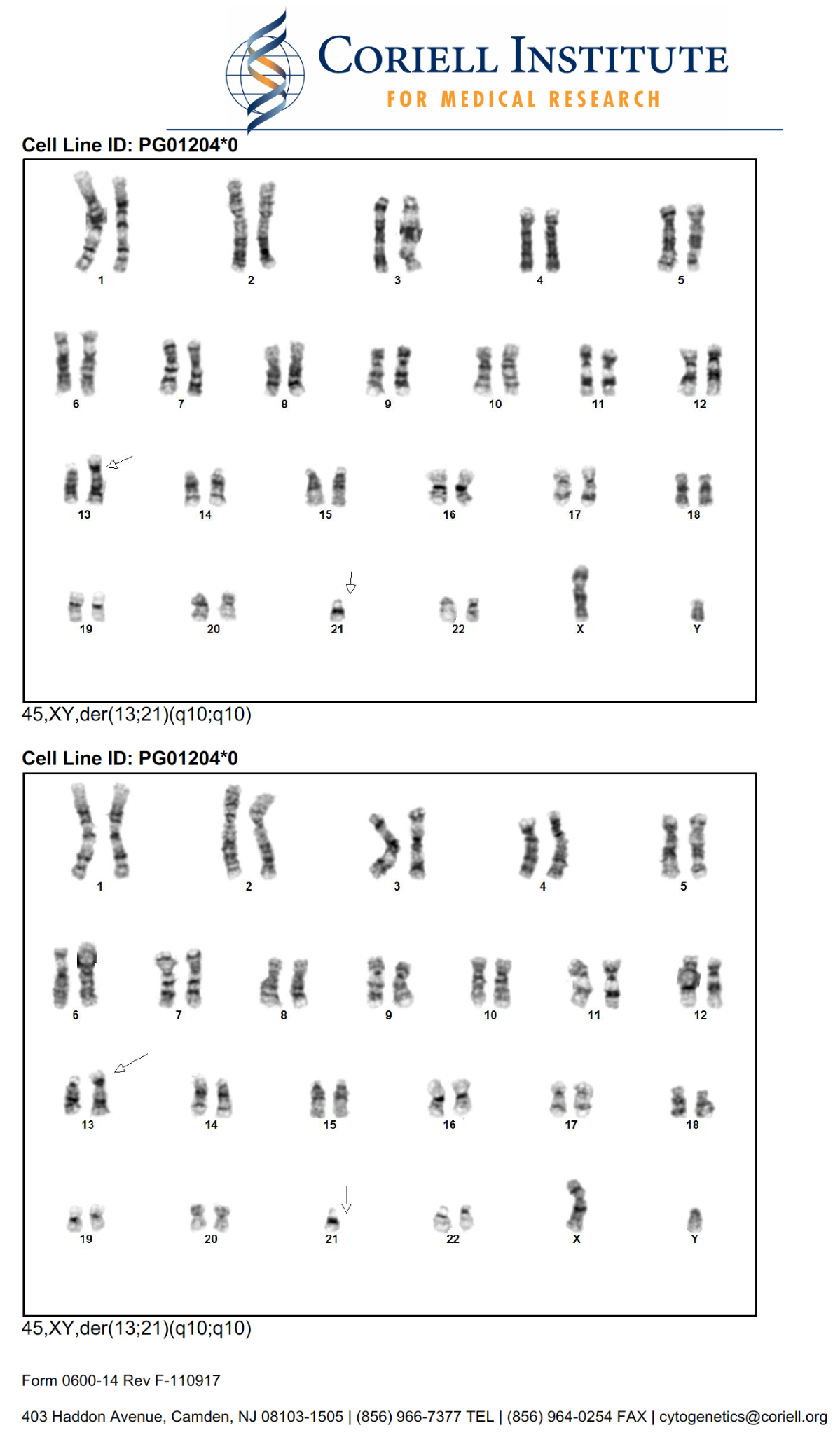

##### Figure S12. G-banding report from Coriell. G-banding was performed for HG011204 (Coriell case id: PG01204*0) at passage 6 on Mar. 20, 2024. Cell counts were 20, 5 cells were analyzed and karyotyped, related to Figure 4B.

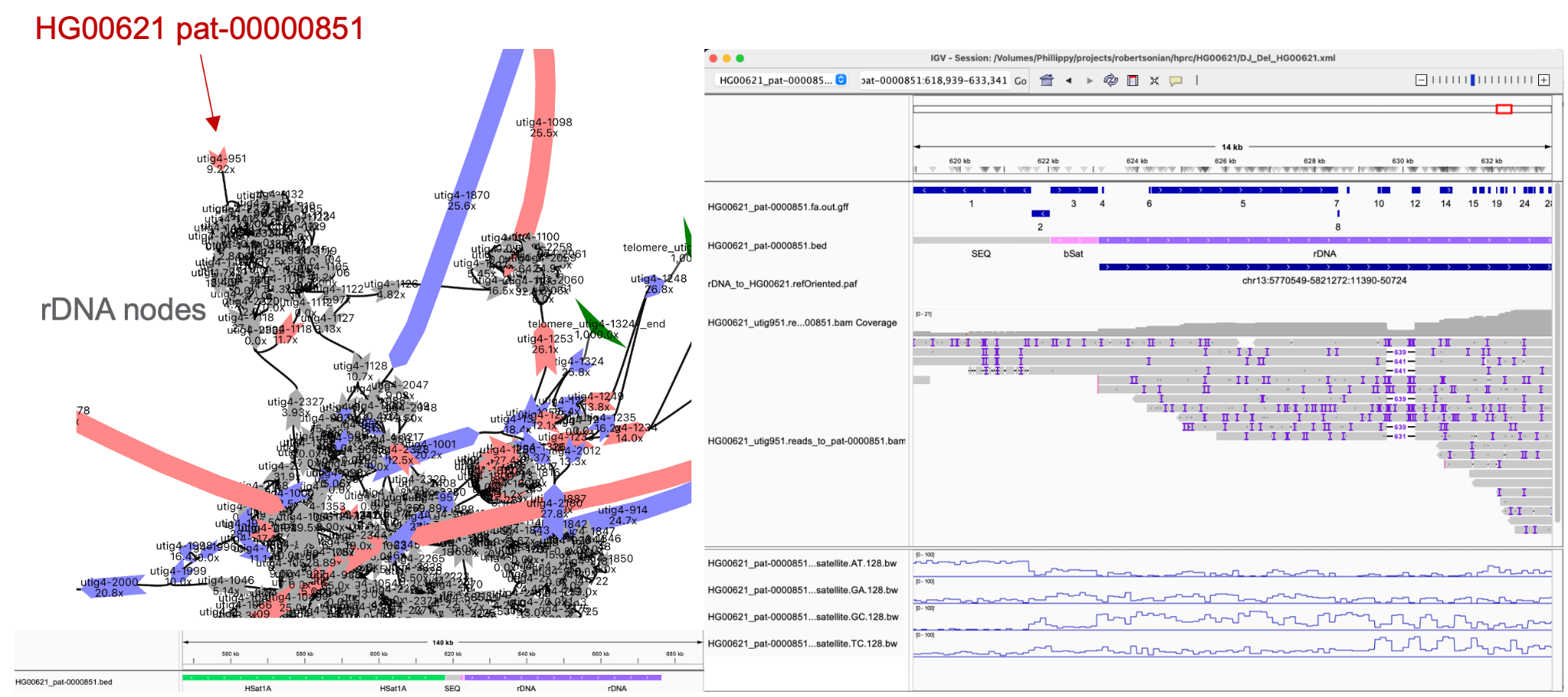

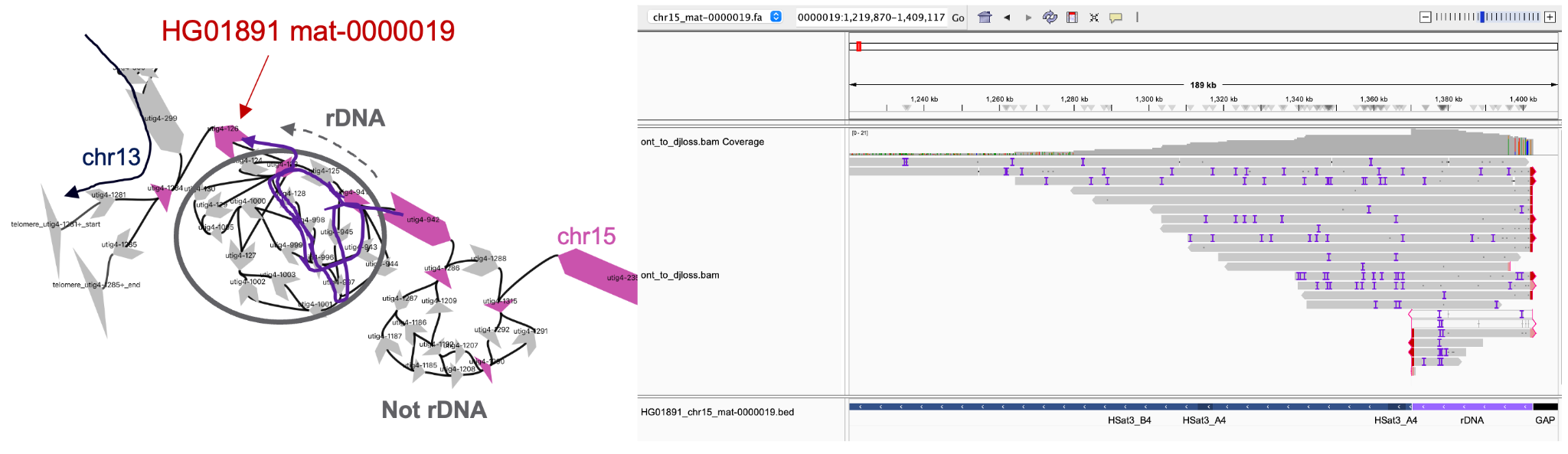

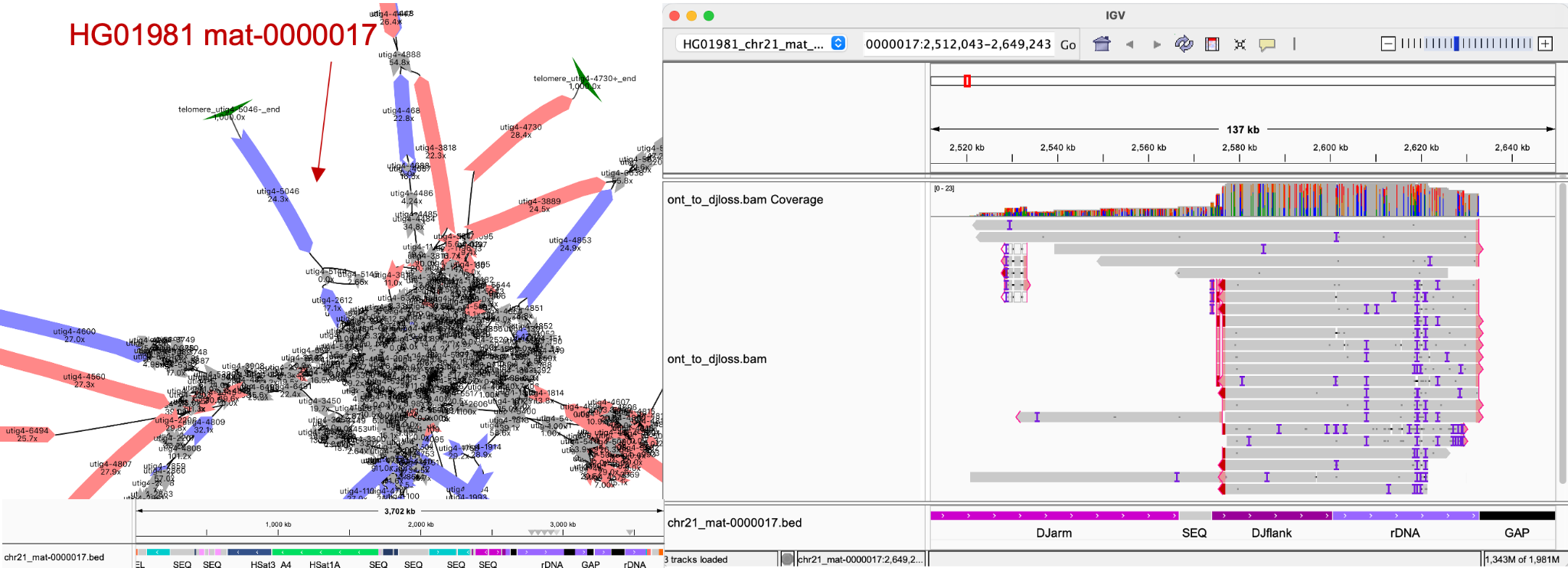

##### Figure S13. Nodes with truncated or missing DJs on the assembly graph and reads supporting the junction, related to Figure 4E.

The nodes marked with arrows correspond to the sequence shown in **Figure 4E**, connected to the rDNA node cluster.

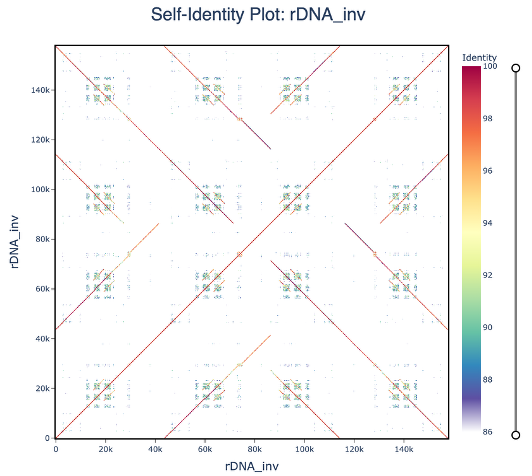

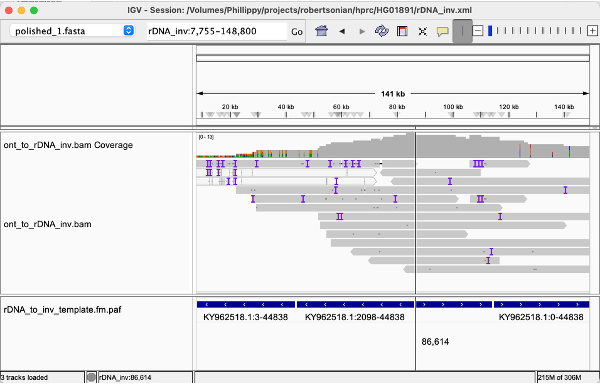

##### Figure S14. Inverted rDNA array found in HG01891, related to Figure 4E.

ONT-UL reads forming the inverted repeat were extracted and mapped to the rDNA consensus KY962518.1. A template has been constructed based on the read alignments, and polished with Flye v2.9.5. The self-identity plot on the top shows the inverted rDNA junction. Below shows the read alignments at the inversion breakpoint (vertical line).

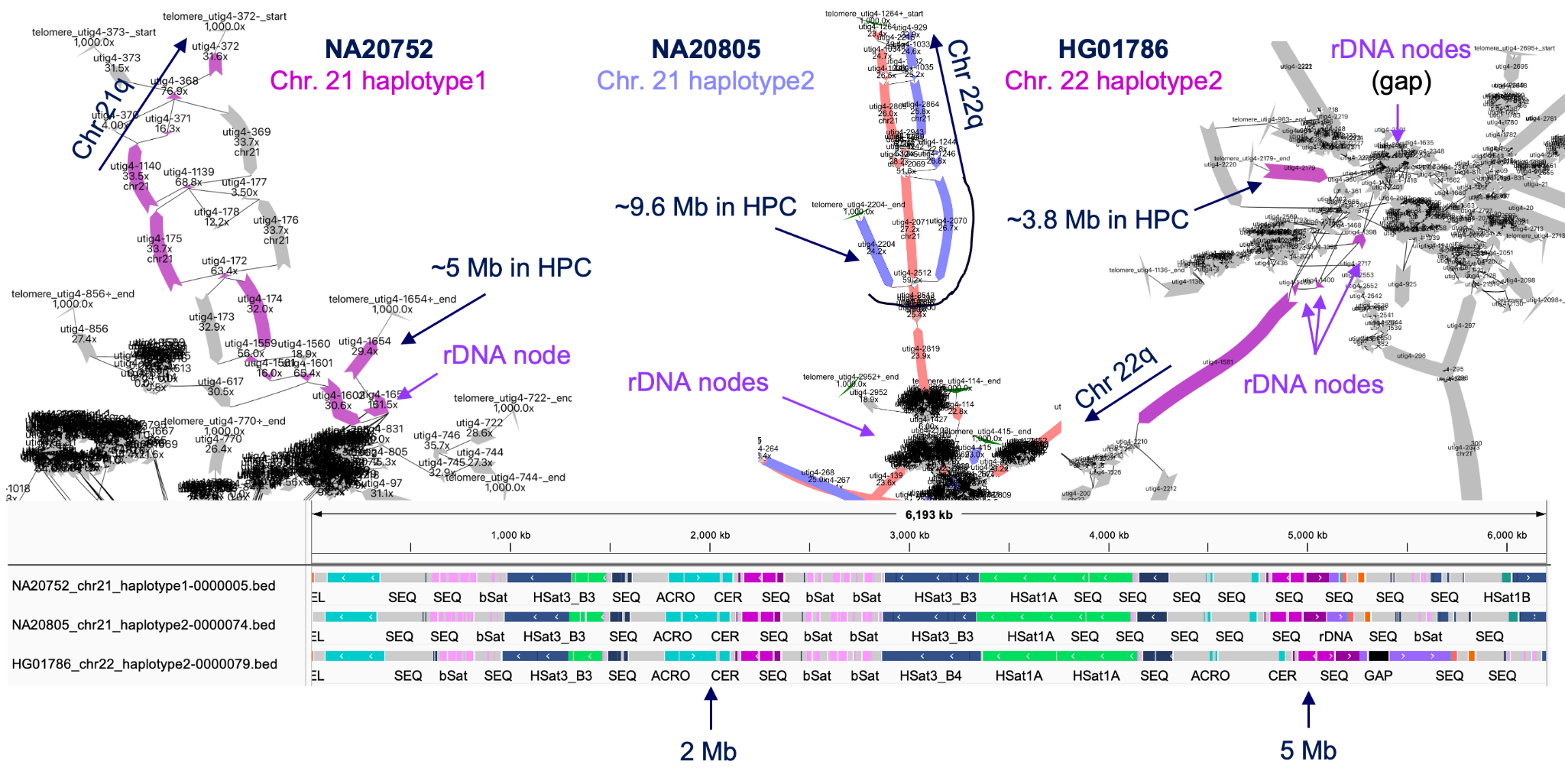

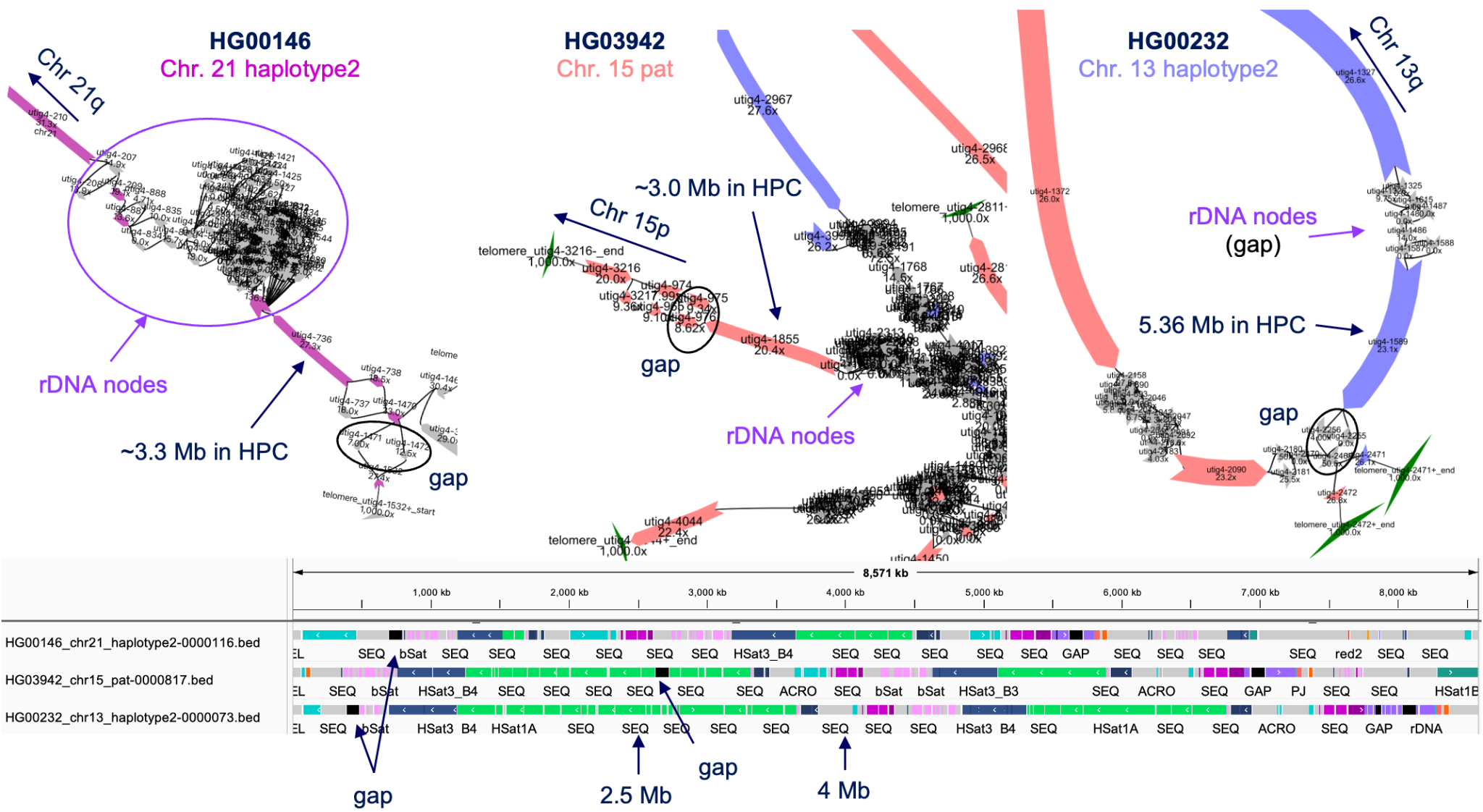

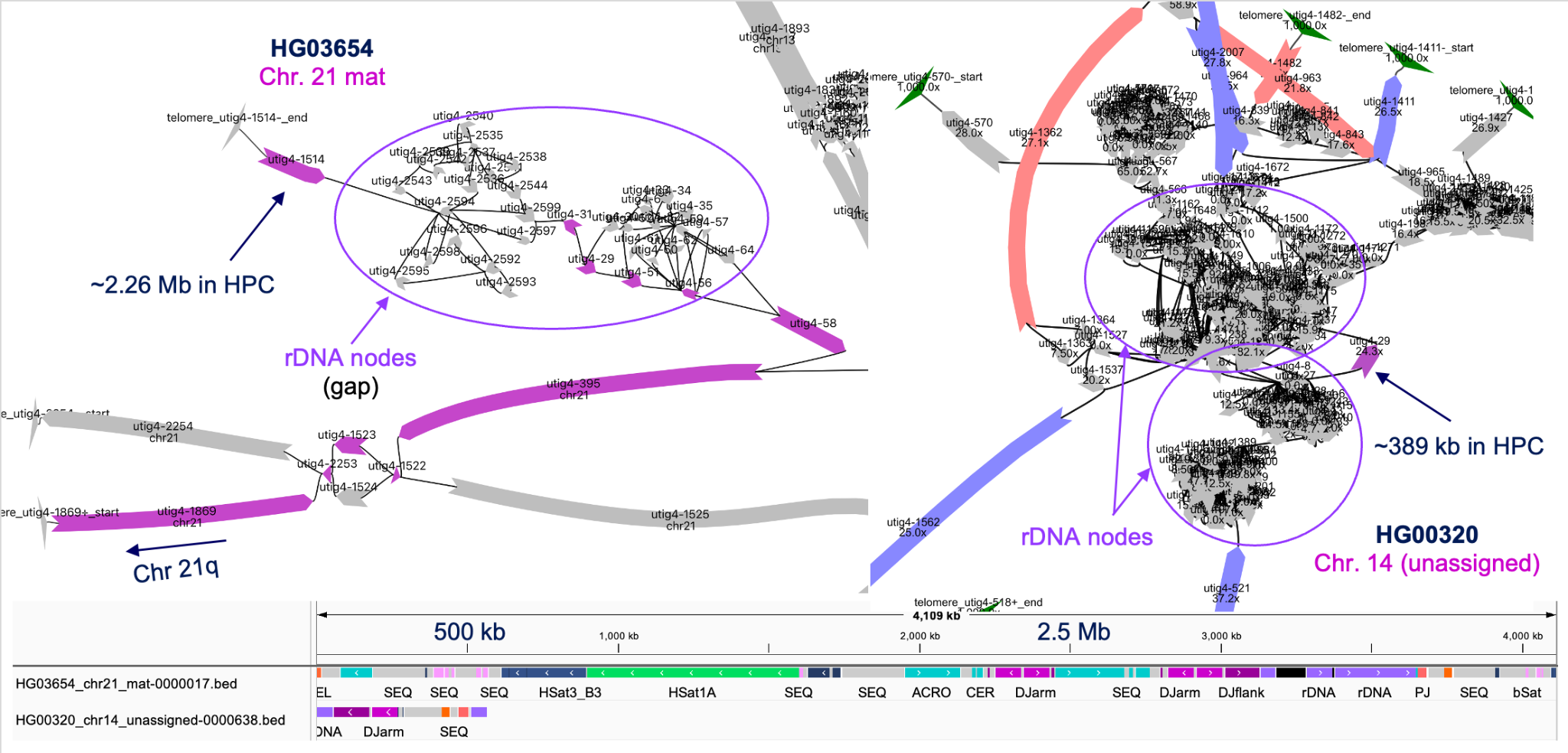

##### Figure S15. Nodes with the additional DJ copy on the assembly graph, related to Figure 4G.

All duplicated DJ blocks shown in **Figure 4G** are found on a single node (pointed in black arrow) in the Verkko assembly graph, indicating it was unambiguously resolved. The node size is shown in homopolymer compressed space (HPC), which is usually smaller than the final consensus. The IGV screen shot shows the consensus repeat annotation with the size in uncompressed space.

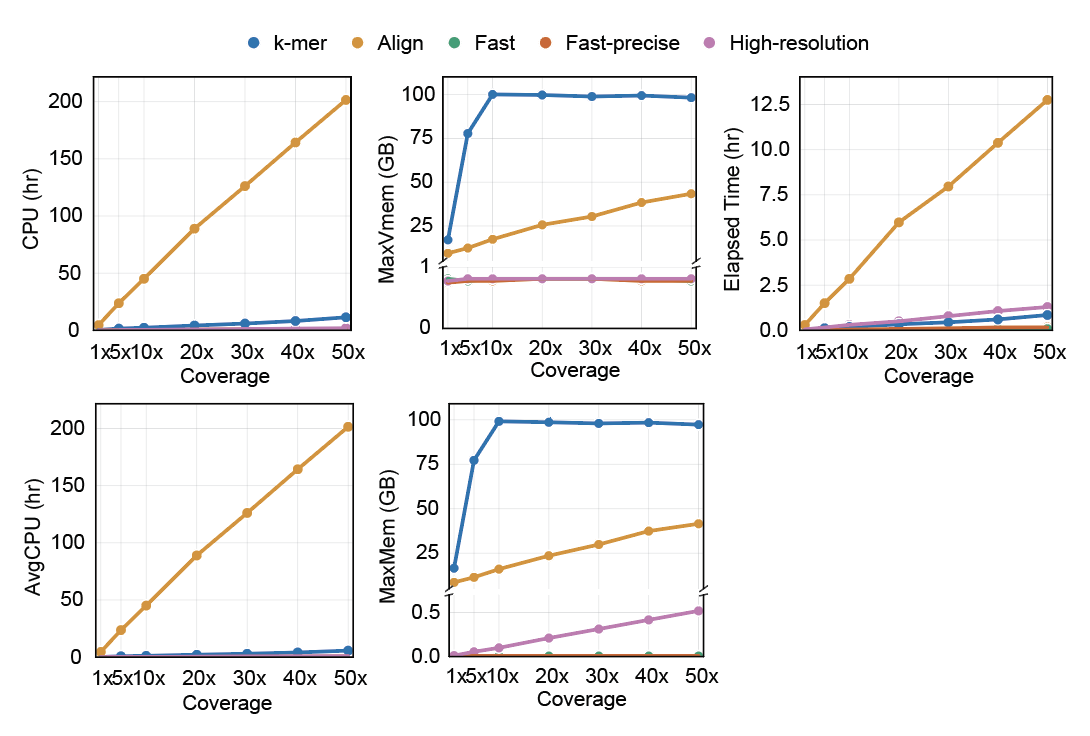

##### Figure S16. Computational resource usage for each DJ calculation method, related to Figure 5A-C.

Total CPU time in hour, maximum memory usage (MaxMem, GB), Elapsed time (wall-clock hours), AvgCPU (average CPU time per record in hours), and MaxVmem (maximum virtual memory usage in GB).

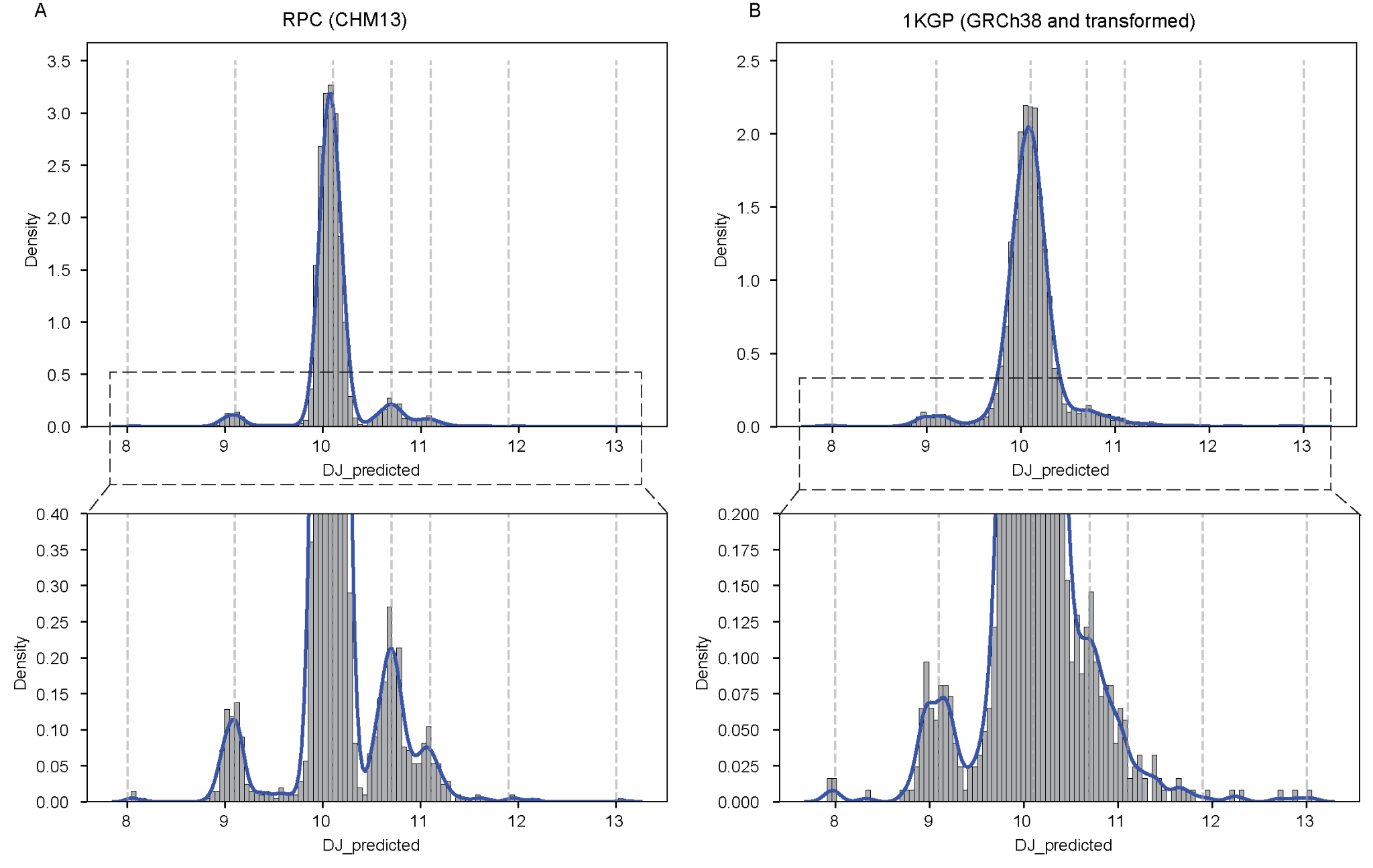

##### Figure S17. Density and kernel plots of DJ copy number distributions in the RPC cohort (CHM13-aligned) and the 1KGP cohort transformed using the Fast-precise model, related to Figure 5E-F.

Dotted vertical lines indicate the fixed component means used for Gaussian mixture model (GMM) fitting.

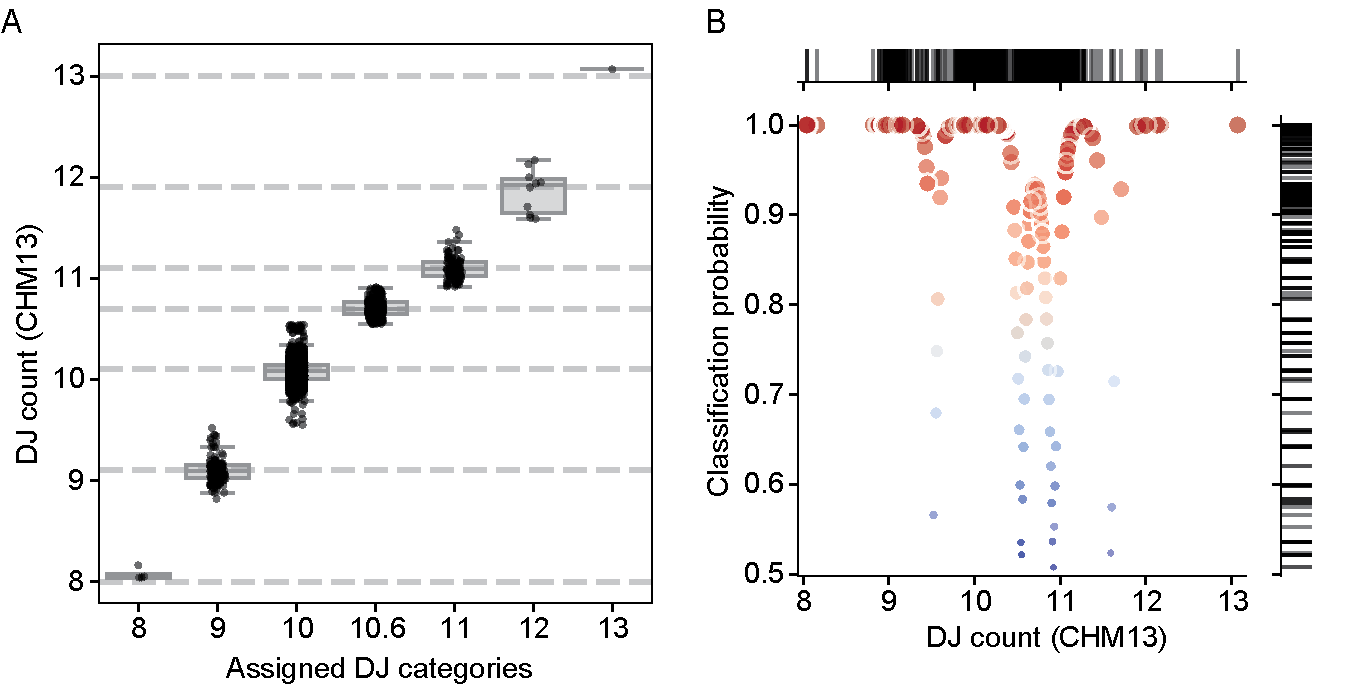

##### Figure S18. Classification results for the RPC cohort (CHM13-aligned) using a Gaussian mixture model (GMM) trained on DJ copy numbers from the 1KGP cohort (GRCh38-aligned) transformed with the Fast-precise model.

(A) DJ copy numbers derived from CHM13 mapping, grouped by each sample’s assigned classification bin. (B) DJ copy numbers and corresponding classification probabilities. Dot color and size represent classification probability, with redder and larger dots indicating higher probability. The color scale corresponds to the y-axis values, with higher values shown in red and lower values shown in blue.

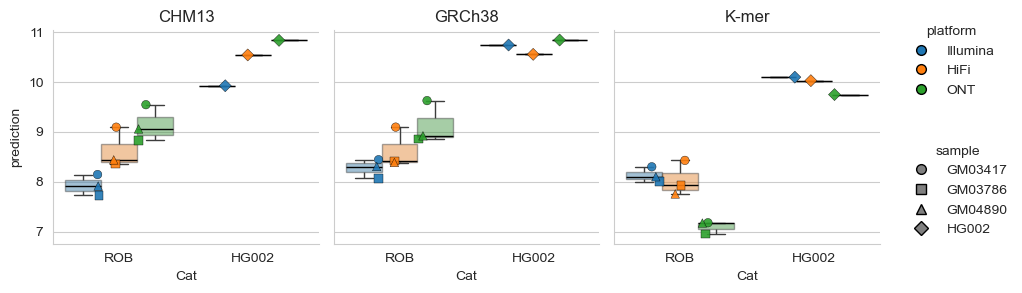

##### Figure S19. Comparison of DJ estimates across sequencing platforms and methods using long-reads and short-reads, related to Figure 5I.

DJ copy-number estimates by sequencing platform and method (mapping-based using GRCh38 and CHM13, and *k*-mer–based). For GRCh38, we used fast-precise mode and applied a transformation model to normalize DJ counts.

### Supplementary Tables

##### Table S1. Target unit for copy-number estimate and masked coordinates on CHM13, related to Figure 1C.

| Chr | Start | End | Remarks | CenSat Annotation |
| --- | --- | --- | --- | --- |
| chr13 | 5424523 | 5770548 | DJ_target | ct_13_7 to ct_13_10 |
| chr13 | 5467957 | 5638362 | ct_13_9_target | ct_13_9 |
| chr13 | 5770548 | 5817416 | rDNA_target |  |
| chr13 | 11644595 | 12819139 | PHR_target | ct_13_31 to ct_13_37 |
| chr13 | 11644595 | 12301366 | PHR_arm1_target | ct_13_33 to ct_13_35 |
| chr13 | 12440010 | 12819139 | PHR_arm2_target | ct_13_36 |
| chr1 | 228022744 | 228024984 | 5S_target |  |
| chrY | 0 | 2458320 | PAR1_masked_forXY |  |
| chrY | 62122809 | 62460029 | PAR2_masked_forXY |  |
| chrY | 0 | 62460029 | EntireY_masked_forXX |  |
| chr14 | 1755893 | 2099537 | chr14_DJ_masked |  |
| chr15 | 2164544 | 2506442 | chr15_DJ_masked |  |
| chr21 | 2762174 | 3108298 | chr21_DJ_masked |  |
| chr22 | 4448073 | 4793794 | chr22_DJ_masked |  |
| chr13 | 5817416 | 9348041 | chr13_rDNA_masked |  |
| chr14 | 2099537 | 2817811 | chr14_rDNA_masked |  |
| chr15 | 2506442 | 4707485 | chr15_rDNA_masked |  |
| chr21 | 3108298 | 5612715 | chr21_rDNA_masked |  |
| chr22 | 4793794 | 5720650 | chr22_rDNA_masked |  |
| chr14 | 6582691 | 7707094 | chr14_PHR_masked |  |
| chr21 | 8706141 | 9832742 | chr21_PHR_masked |  |
| chr1 | 227743723 | 228022744 | chr1_5S_array_masked |  |

##### Table S2. Target region used for the pilot "fast" mode across the UK Biobank samples, related to Figure 1D. Target DJ length of 136,405 bps was used for normalization.

| **Sequence** | **Start** | **End** |
| --- | --- | --- |
| chr17_GL000205v2_random | 82043 | 180103 |
| chr21 | 5499029 | 5539854 |
| chr21 | 5572242 | 5608781 |
| chr21 | 6212188 | 6249305 |
| chr21 | 6282205 | 6377069 |
| chr21 | 6630182 | 6706438 |
| chr21 | 6738884 | 6739084 |
| chr21 | 6984220 | 7034805 |
| chr21 | 7067322 | 7104007 |
| chr21 | 7377866 | 7416911 |
| chr21 | 7449274 | 7485785 |
| chr21 | 7652619 | 7690698 |
| chrUn_GL000195v1 | 26146 | 64537 |
| chrUn_GL000195v1 | 98567 | 182895 |

##### Table S3. Target DJ region on GRCh38, related to Figure 1D. Target size used for normalization is 346,026 bp.

| **Sequence** | **Start** | **End** | **Direction** | **Alignment block length** | **Idy (%)** | **Coverage (%)** | **Final_Filter** |
| --- | --- | --- | --- | --- | --- | --- | --- |
| chr21 | 5,499,028 | 5,796,009 | + | 302,892 | 76.3 | 69.2 | Included |
| chr21 | 6,211,371 | 6,377,258 | - | 171,633 | 95.5 | 49.5 | Included |
| chr21 | 6,984,219 | 7,149,527 | + | 171,119 | 95.2 | 49.4 | Included |
| chr21 | 7,550,890 | 7,692,975 | - | 142,679 | 98.6 | 41.2 | Included |
| chr21 | 7,377,865 | 7,500,890 | + | 128,765 | 94.1 | 37.1 | Included |
| chr21 | 6,630,181 | 6,739,085 | + | 114,258 | 94.0 | 33.0 | Included |
| chr21 | 8,099,839 | 8,202,044 | + | 102,369 | 99.2 | 29.5 | Included |
| chr21 | 8,310,971 | 8,385,090 | + | 74,247 | 99.3 | 21.4 | Included |
| chr21 | 5,849,945 | 5,899,543 | + | 49,680 | 99.2 | 14.3 | Included |
| chr21 | 5,889,728 | 5,916,593 | + | 26,909 | 99.3 | 7.8 | Included |
| chr17_GL000205v2_random | 38,600 | 182,663 | + | 144,191 | 99.0 | 41.6 | Included |
| chr22_KI270733v1_random | 0 | 118,281 | + | 118,549 | 99.2 | 34.2 | Included |
| chrUn_GL000195v1 | 0 | 182,896 | - | 183,113 | 98.8 | 52.8 | Included |
| chrUn_GL000220v1 | 0 | 101,548 | + | 101,669 | 99.2 | 29.4 | Included |
| chr4 | 190,051,896 | 190,061,962 | - | 10,232 | 81.9 | 2.8 | Filtered out |
| chr4 | 190,051,911 | 190,061,986 | + | 10,280 | 81.9 | 2.8 | Filtered out |
| chr10 | 133,651,258 | 133,661,200 | + | 10,139 | 83.4 | 2.8 | Filtered out |
| chr10 | 133,651,243 | 133,661,162 | - | 10,076 | 83.3 | 2.8 | Filtered out |

##### Table S4. Different modes for calculating DJ counts from GRCh38 alignments, related to Figures 1D, 5A-D.

| **Mode Name** | **Dup / Supplementary Read Removal** | **Target Calculate** | **Background Coverage** | **R2 with CHM13 Value** | ***p*-value of R2** | **RSS after fitting** |
| --- | --- | --- | --- | --- | --- | --- |
| Fast | No | Read counts | Median(autosome(read count/ length)) | 9.95E-01 | 7.49E-92 | 0.71 |
| Fast-precise | Yes | Read counts | Read counts of whole autosome / whole autosome length | **9.96E-01** | **4.62E-97** | **0.55** |
| High-resolution | Yes | Sum (per-base depth) | Median(autosome coverage) | **9.96E-01** | 3.67E-95 | 0.61 |

##### Table S5. Pipeline confirms 8 DJs in 3 known ROB cell lines and two control samples, where the DJ was karyotyped before.

|  |  | **DJ count values** | | |  |  |
| --- | --- | --- | --- | --- | --- | --- |
| **Sample** | **Sex** | **T2T-CHM13** | **GRCh38** | **k-mer** | **Remarks** | **Age** |
| CHM13 | XX | 10.3 | 10.1 | 10.2 | CHM13hTERT cell line, karyotypically normal | na |
| HG002 | XY | 10.1 | 10.7 | 10.1 | GM24385 cell line, karyotypically normal | na |
| GM03417 | XX | 8.3 | 8.4 | 8.3 | Clinical features of Down syndrome. Mosaic; 32% of cells are balanced 45,XX,t(14;21);  *Cellular heterogeneity with regard to the number of copies of Chr. 21, with 3 copies of Chr. 21 being relatively rare (~10%) | 15 |
| GM03786 | XX | 8.0 | 8.1 | 8.0 | Clinically normal | 19 |
| GM04890 | XX | 8.1 | 8.3 | 8.1 | Clinically normal; has had 5 miscarriages | 32 |

##### Exact DJ count values produced from each method are shown. Related to Figure 1F-H. Remarks are from Coriell. *Reported from de Lima et al., Nature, 2025

##### Table S6. Copy number estimates of the PHRs across the RPC cohort (n=4,187) and UK Biobank WGS cohort (n=490,374), related to Table 2.

| **Cohort** | **Num. of PHRs** | **Num. of samples** | **%** | **XX** | **%** | **XY** | **%** |
| --- | --- | --- | --- | --- | --- | --- | --- |
| RPC | <4 | 3 | 0.07% | 0 | 0.00% | 3 | 0.15% |
|  | 4 | 261 | 6.23% | 126 | 5.93% | 135 | 6.54% |
|  | 5 | 1,645 | 39.29% | 828 | 39.00% | 817 | 39.58% |
|  | 6 | 2,189 | 52.28% | 1,127 | 53.09% | 1,062 | 51.45% |
|  | 7 | 85 | 2.03% | 39 | 1.84% | 46 | 2.23% |
|  | >7 | 4 | 0.10% | 3 | 0.14% | 1 | 0.05% |
|  | Total | 4,187 | 100.00% | 2,123 | 100.00% | 2,064 | 100.00% |
| RPC  (Parents only) | <4 | 1 | 0.04% | 0 | 0.00% | 1 | 0.08% |
|  | 4 | 143 | 5.47% | 67 | 5.14% | 76 | 5.81% |
|  | 5 | 931 | 35.64% | 453 | 34.74% | 478 | 36.54% |
|  | 6 | 1,477 | 56.55% | 755 | 57.90% | 722 | 55.20% |
|  | 7 | 59 | 2.26% | 28 | 2.15% | 31 | 2.37% |
|  | >7 | 1 | 0.04% | 1 | 0.08% | 0 | 0.00% |
|  | Total | 2,612 | 1 | 1,304 | 100.00% | 1,308 | 100.00% |
| UK Biobank | <4 | 234 | 0.05% | 128 | 0.05% | 106 | 0.05% |
|  | 4 | 33,964 | 6.93% | 18,195 | 6.84% | 15,769 | 7.03% |
|  | 5 | 185,625 | 37.85% | 100,601 | 37.82% | 85,024 | 37.89% |
|  | 6 | 262,723 | 53.58% | 142,803 | 53.69% | 119,920 | 53.45% |
|  | 7 | 7,631 | 1.56% | 4,164 | 1.57% | 3,467 | 1.55% |
|  | >7 | 197 | 0.04% | 104 | 0.04% | 93 | 0.04% |
|  | Total | 490,374 | 100.00% | 265,995 | 100.00% | 224,379 | 100.00% |

##### Table S7. DJ copy numbers identified in families of the UK Biobank, related to Figure 3G.

| **Mother** | **Father** | **Offspring** | **N** |
| --- | --- | --- | --- |
| 8 | 10 | 10 | 2 |
| 9 | 10 | 9 | 16 |
| 9 | 10 | 10 | 8 |
| 9 | 11 | 10 | 3 |
| 9 | >11 | 10 | 1 |
| 10 | 9 | 9 | 15 |
| 10 | 9 | 10 | 10 |
| 10 | 10 | 9 | 3 |
| 10 | 10 | 10 | 845 |
| 10 | 10 | 11 | 2 |
| 10 | 11 | 10 | 35 |
| 10 | 11 | 11 | 27 |
| 10 | >11 | 11 | 1 |
| 10 | >11 | >11 | 1 |
| 11 | 9 | 9 | 1 |
| 11 | 9 | 10 | 1 |
| 11 | 9 | 11 | 1 |
| 11 | 10 | 10 | 25 |
| 11 | 10 | 11 | 31 |
| 11 | 11 | 10 | 2 |
| 11 | 11 | 11 | 2 |
| 11 | 11 | >11 | 4 |
| 11 | >11 | 11 | 1 |
| 11 | >11 | >11 | 1 |
| >11 | 10 | 10 | 2 |

##### Table S8. Copy number estimates of the DJs across the 1KGP samples (n=3,202), related to Figure 4B.

| **Cohort** | **Num. of DJs** | **Num. of samples** | **%** | **XX** | **%** | **XY** | **%** |
| --- | --- | --- | --- | --- | --- | --- | --- |
| 1KGP | 8 | 8 | 0.25% | 3 | 0.19% | 5 | 0.31% |
|  | 9 | 114 | 3.56% | 52 | 3.24% | 62 | 3.88% |
|  | 10 | 2,925 | 91.35% | 1,470 | 91.65% | 1,455 | 91.05% |
|  | 11 | 145 | 4.53% | 73 | 4.55% | 72 | 4.51% |
|  | >11 | 10 | 0.31% | 6 | 0.37% | 4 | 0.25% |
|  | Total | 3,202 | 100.00% | 1,604 | 100.00% | 1,598 | 100.00% |
| 1KGP  (Parents only) | **8** | **8** | **0.25%** | **2** | **0.16%** | **2** | **0.16%** |
|  | 9 | 114 | 3.56% | 44 | 3.47% | 50 | 4.06% |
|  | 10 | 2,925 | 91.35% | 1,156 | 91.17% | 1,123 | 91.08% |
|  | 11 | 145 | 4.53% | 62 | 4.89% | 54 | 4.38% |
|  | >11 | 10 | 0.31% | 4 | 0.32% | 4 | 0.32% |
|  | Total | 3,202 | 100.00% | 1,268 | 100.00% | 1,233 | 100.00% |

##### Table S9. ROB candidates and their family members found in 1KGP, related to Figure 4B.

| **Sample** | **Sex** | **Group** | **GRCh38** | ***k*-mer** | **Relationship** |
| --- | --- | --- | --- | --- | --- |
| HG01204 | XY | 8 | 7.99 | 7.85 | Father |
| HG01205 | XX | 10 | 10.48 | 10.58 | Mother |
| HG01206 | XY | 8 | 7.91 | 7.92 | Son |
| HG00650 | XY | 10 | 9.98 | 9.94 | Father |
| HG00651 | XX | 8 | 7.95 | 7.93 | Mother |
| HG00652 | XX | 8 | 8.06 | 8.07 | Daughter |
| NA18868 | XY | 8 | 8.01 | 8.00 | Father |
| NA18867 | XX | 10 | 10.39 | 9.93 | Mother |
| NA18869 | XY | 10 | 10.00 | 9.85 | Son |
| HG02687 | XY | 10 | 10.06 | 9.71 | Father |
| HG02688 | XX | 8 | 8.33 | 8.07 | Mother |
| HG02689 | XY | 9 | 8.94 | 9.10 | Son |
| NA12812 | XY | 9 | 8.93 | 9.07 | Father |
| NA12813 | XX | 9 | 9.38 | 9.11 | Mother |
| NA12801 | XY | 8 | 8.15 | 8.22 | Son |
| NA11891 | XY | 8 | 8.38 | 8.37 | Father |
| NA11892 | XX | 11 | 10.69 | 10.64 | Mother |
| NA10865 | XY | 10 | 10.22 | 10.00 | Son |

Samples with likely 8 copies are highlighted with a green background. NA12801 was the only sample with 8 DJs that did not match any of the parents. NA11891 is binned as 8 copies with lower confidence, staying on the higher end from the rest of the individuals binned as 8 DJs.

##### Table S10. Target sequences used for acrocentric repeat annotation, related to Figure 4C,D,E,G.

| **Target (sat)** | **Region** | **Direction** | **Color (col)** | **LengthFilter (len)** |
| --- | --- | --- | --- | --- |
| HSat1A | 13_1A_3.peak_1.3013bp-consensus | + | 0,222,96 | 7000 |
| HSat1B | chr13:10169300-10215148 | + | 27,153,139 | 7000 |
| HSat3_A4 | chr13:5000117-5152650 | - | 37,59,100 | 2000 |
| HSat3_A5 | 15_3_12.peak_1.1794bp-consensus | + | 51,130,165 | 2000 |
| HSat3_B3 | 13_3_3.peak_2.5390bp-consensus | + | 51,81,137 | 7000 |
| HSat3_B4 | chr13:5783-61195 | - | 51,81,137 | 2000 |
| HSat3Y | chrY:58654600-58660873 | - | 51,0,153 | 7000 |
| aSat | chr13:15547594-17498291 | - | 153,0,0 | 7000 |
| bSat | chr13:4973578-4985238 | + | 250,153,255 | 7000 |
| ACRO | chr13:5338347-5358151 | + | 204,204 | 7000 |
| CER | chr13:5369401-5424523 | - | 204,204 | 7000 |
| CENY | chrY:10699757-10745416 | - | 153,0,0 | 7000 |
| SST1 | chr13:12301367-12440010 | - | 204,204 | 7000 |
| DJarm | chr13:5555305-5643740 | + | 204,0,204 | 7000 |
| DJflank | chr13:5650824-5770548 | + | 153,0,153 | 7000 |
| rDNA | KY962518, rotated to begin at the 18S | + | 153,102,255 | 7000 |
| PHR_LOC | chr13:9442748-9474060 | + | 255,102,0 | 7000 |
| PJ | chr13:9356167-9389035 | + | 255,102,102 | 8000 |
| P5-AZFb | chrY:18878839-18989510 | + | 255,153,0 | 8000 |
| teal2 | chrY:23213314-23328994 | + | 128,128 | 8000 |
| P3-AZFc | chrY:23327300-23566501 | + | 60,84,164 | 8000 |
| blue2 | chrY:23328993-23582346 | + | 60,84,164 | 8000 |
| red2 | chrY:24037533-24167130 | + | 231,35,35 | 8000 |
| green2 | chrY:24167430-24482113 | + | 11,130,67 | 8000 |
| yellow1 | chrY:24482112-25159839 | + | 255,153,0 | 8000 |
| gray1 | chrY:25389078-25503792 | + | 125,112,96 | 8000 |
| SEQ | . | . | 204,204,204 | 0 |
| TEL | . | . | 255,102,51 | 0 |
| GAP | . | . | 0,0,0 | 0 |

For HSat1A, HSat3_A5 and HSat3_B3, the original consensus sequences used for CHM13v2.0 HSat annotation were obtained from https://github.com/altemose/HSatReview (Altemose et al., Science, 2022). For the rest, the annotated region from the CHM13v2.0 CenSatv2.0 track was used to extract the target sequences. SEQ indicates none of the other target sequences were annotated as a placeholder. Tel, Telomere. GAP, N-base gaps. RGB color codes were given for visualization.

##### Table S11. Copy number estimates of the DJs in 1KGP samples identified as 9 and 11 DJs among the HPRCr2 samples, related to Figure 4C,D,E,G.

|  | **GRCh38** | | | | ***k*-mer** | | |  |  |  |  |
| --- | --- | --- | --- | --- | --- | --- | --- | --- | --- | --- | --- |
| **Sample** | **Read Count** | **Back-ground** | **DJ raw** | **DJ pred.** | **DJ medCov** | **PeakCP2** | **Cnt.** | **Pop** | **Set** | **Assembly** | **Status** |
| HG00658 | 353197 | 0.22 | 9.20 | 9.0 | 123 | 28 | 8.8 | CHS | 698rel | 9 short-arms resolved, 10th ending in mat Chr. 14 rDNA tangle (no outgoing edge) | Unresolved |
| HG01891 | 353221 | 0.22 | 9.20 | 9.0 | 124 | 28 | 8.9 | ACB | 698rel | chr15 mat-0000019 | Resolved |
| HG03521 | 367793 | 0.23 | 9.39 | 9.2 | 129 | 29 | 8.9 | ESN | 2504 | 9 short-arms resolved, 10th assembled down to rDNA. Assembly graph has only 9 outgoing non-q-arm paths from rDNA | Unresolved |
| HG00621 | 347037 | 0.22 | 9.20 | 9.0 | 121 | 27 | 9.0 | CHS | 698rel | pat-0000851 | Junction resolved |
| HG02818 | 384595 | 0.23 | 9.48 | 9.3 | 126 | 28 | 9.0 | GWD | 698rel | *hifiasm ver., DJ too fragmented | Unresolved |
| HG01981 | 382218 | 0.23 | 9.63 | 9.4 | 128 | 28 | 9.1 | PEL | 698rel | chr21 mat-0000017 | Resolved |
| HG03942 | 435356 | 0.23 | 11.13 | 10.9 | 148 | 27 | 11.0 | BEB | 698rel | chr15 pat-0000817 | Resolved |
| HG00320 | 506539 | 0.27 | 11.03 | 10.8 | 174 | 32 | 10.9 | FIN | 2504 | chr14 unassigned-0000638 | partially resolved |
| HG01943 | 443821 | 0.22 | 11.49 | 11.2 | 151 | 28 | 10.8 | PEL | 698rel | mat-0000010 has DJ arms on both sides, but with a gap | Unresolved |
| HG01786 | 672642 | 0.34 | 11.32 | 11.1 | 230 | 43 | 10.7 | IBS | 2504 | chr22 haplotype2-0000079 | Resolved |
| HG02071 | 420087 | 0.22 | 11.06 | 10.8 | 154 | 29 | 10.6 | KHV | 698rel | DJ on multiple small contigs | Unresolved |
| HG03654 | 431992 | 0.23 | 10.81 | 10.6 | 143 | 27 | 10.6 | PJL | 698rel | chr21 mat-0000017 | Resolved |
| NA20752 | 411396 | 0.21 | 11.10 | 10.8 | 142 | 27 | 10.5 | TSI | 2504 | chr21 haplotype1-0000005 | Resolved |
| HG00232 | 699307 | 0.36 | 11.27 | 11.0 | 246 | 47 | 10.5 | GBR | 2504 | chr13 haplotype2-0000073 | Resolved |
| HG00146 | 540155 | 0.29 | 10.84 | 10.6 | 187 | 36 | 10.4 | GBR | 2504 | chr21 haplotype2-0000116 | Resolved |
| NA20805 | 438208 | 0.23 | 10.87 | 10.6 | 150 | 29 | 10.3 | TSI | 2504 | chr21 haplotype2-0000074 | Resolved |

These samples were targeted for investigating the assemblies.
