## Supplemental list of authors for "Biobank-scale genotyping of Robertsonian translocations reveals hidden structural variation on the human acrocentric chromosomes"

Derek Albracht^1^, Ivan A. Alexandrov^2^, Jamie Allen^3^, Alawi A. Alsheikh-Ali^4^, Nicolas Altemose^5^, Casey Andrews^6^, Dmitry Antipov^7^, Lucinda Antonacci-Fulton^1^, Alexander Arguello^8^, Mobin Asri^9^, Marcelo Ayllon^10^, Jennifer R. Balacco^11^, Floris P. Barthel^12^, Edward A. Belter Jr^1^, Halle D. Bender^9^, Andrew P. Blair^9^, Davide Bolognini^13^, Katherine E. Bonini^14^, Christina Boucher^15^, Guillaume Bourque^16,17,18^, Silvia Buonaiuto^19^, Shuo Cao^19^, Andrew Carroll^20^, Ann M. Mc Cartney^21^, Monika Cechova^9^, Mark J.P. Chaisson^22^, Pi-Chuan Chang^20^, Xian Chang^9^, Jitender Cheema^3^, Haoyu Cheng^23^, Claudio Ciofi^24^, Hiram Clawson^9^, Sarah Cody^1^, Vincenza Colonna^19^, Holland C. Conwell^25^, Robert Cook-Deegan^26^, Mark Diekhans^9^, Maria Angela Diroma^24^, Daniel Doerr^27,28,29^, Zheng Dong^6^, Danilo Dubocanin^5^, Richard Durbin^30,31^, Jana Ebler^27,32^, Evan E. Eichler^10,33^, Jordan M. Eizenga^9^, Parsa Eskandar^9^, Eddie Ferro^15^, Anna-Sophie Fiston-Lavier^34,35^, Sarah M. Ford^25^, Willard W. Ford^36^, Giulio Formenti^11^, Adam Frankish^3^, Mallory A. Freeberg^3^, Qichen Fu^6^, Stephanie M. Fullerton^37^, Robert S. Fulton^1^, Shenghan Gao^38^, Yan Gao^39^, Gage H. Garcia^10^, Obed A. Garcia^40^, Joshua M.V. Gardner^9^, Shilpa Garg^41^, Erik Garrison^19^, Nanibaa' A. Garrison^42,43,44^, John E. Garza^1^, Margarita Geleta^45,46^, Mohammadmersad Ghorbani^47^, Tina A. Graves-Lindsay^1^, Richard E. Green^25^, Carol W. Greider^48^, Cristian Groza^49^, Bida Gu^22^, Andrea Guarracino^12,19^, Melissa Gymrek^50^, Maximilian Haeussler^9^, Leanne Haggerty^3^, Ira M. Hall^51,52^, Nancy F. Hansen^7^, Yue Hao^12^, Mohammad Amiruddin Hashmi^4^, David Haussler^9^, Prajna Hebbar^9^, Peter Heringer^27,28,29^, Glenn Hickey^9^, Todd L. Hillaker^9^, S. Nakib Hossain^3^, Neng Huang^39,53^, Sarah E. Hunt^3^, Toby Hunt^3^, Alexander G. Ioannidis^5,9,46^, Nafiseh Jafarzadeh^9^, Nivesh Jain^11^, Erich D. Jarvis^11,33^, Maryam Jehangir^12^, Juan Jiang^6^, Eimear E. Kenny^14^, Juhyun Kim^7^, Bonhwang Koo^11^, Sergey Koren^7^, Milinn Kremitzki^1,6^, Charles H. Langley^54^, Ben Langmead^55^, Heather A. Lawson^6^, Daofeng Li^6^, Heng Li^39,53^, Ronghan Li^6^, Wen-Wei Liao^51,52^, Jiadong Lin^10^, Tianjie Liu^6^, Glennis A. Logsdon^38^, Ryan Lorig-Roach^9^, Jonathan LoTempio Jr^21,56^, Hailey Loucks^9^, Jane E. Loveland^3^, Jianguo Lu^57^, Shuangjia Lu^51,52^, Julian K. Lucas^9^, Walfred Ma^22^, Juan F. Macias-Velasco^1,6,58^, Kateryna D. Makova^59^, Maximillian G. Marin^39,53^, Christopher Markovic^1^, Tobias Marschall^27,32^, Franco L. Marsico^19^, Fergal J. Martin^3^, Mira Mastoras^9^, Capucine Mayoud^34^, Brandy McNulty^9^, Jack A. Medico^11^, Julian M. Menendez^9^, Karen H. Miga^9^, Anna Minkina^60^, Matthew W. Mitchell^61^, Saswat K. Mohanty^62^, Younes Mokrab^47,63,64^, Jean Monlong^65^, Shabir Moosa^47^, Avelina Moreno-Ochando^66,67^, Shinichi Morishita^68^, Jonathan M. Mudge^3^, Katherine M. Munson^10^, Njagi Mwaniki^69^, Nasna Nassir^4^, Chiara Natali^24^, Shloka Negi^9^, Lingbin Ni^10^, Adam M. Novak^9^, Faith Okamoto^9^, Keisuke K. Oshima^38^, Pilar N. Ossorio^70,71^, Chie Owa^68^, Sadye Paez^11^, Benedict Paten^9^, Clelia Peano^13,72^, Adam M. Phillippy^7,55,73,74^, Brandon D. Pickett^7^, Laura Pignata^19^, Nadia Pisanti^69^, David Porubsky^10,75^, Pjotr Prins^19^, Timofey Prodanov^27,32^, Anandi Radhakrishnan^9^, T. Rhyker Ranallo-Benavidez^12^, Brian J. Raney^9^, Mikko Rautiainen^76^, Alessandro Raveane^13^, Andreas Rechtsteiner^48^, Luyao Ren^10,33^, Arang Rhie^7^, Fedor Ryabov^77,78^, Samuel Sacco^25^, Farnaz Salehi^19^, Michael C. Schatz^55,79^, Laura B. Scheinfeldt^80^, Aarushi Sehgal^36^, William E. Seligmann^25^, Mahsa Shabani^81^, Kishwar Shafin^20^, Shadi Shahatit^34^, Ruhollah Shemirani^14^, Vikram S. Shivakumar^55^, Swati Sinha^3^, Jouni Sirén^9^, Linnéa Smeds^62^, Steven J. Solar^7^, Marco Sollitto^11,24^, Nicole Soranzo^13,30,82^, Andrew B. Stergachis^10,60^, Marie-Marthe Suner^3^, Yoshihiko Suzuki^68^, Arda Söylev^27,32^, Ahmad Abou Tayoun^83,84^, Jack A.S. Tierney^3^, Chad Tomlinson^1^, Francesca Floriana Tricomi^3^, Mohammed Uddin^4,85^, Matteo Tommaso Ungaro^25,86^, Rahul Varki^15^, Flavia Villani^19^, Ivo Violich^9^, Mitchell R. Vollger^87^, Brian P. Walenz^7^, Charles Wang^88^, Lisa E. Wang^14^, Ting Wang^1,6,58^, Aaron M. Wenger^89^, Conor V. Whelan^11^, Zilan Xin^6^, Zheng Xu^6^, Kai Ye^90^, DongAhn Yoo^10^, Wenjin Zhang^6^, Ying Zhou^39^, Xiaoyu Zhuo^6^, Giulia Zunino^13^

### Affiliations

^1^ McDonnell Genome Institute, Washington University School of Medicine, St. Louis, MO 63108, USA

^2^ Department of Human Molecular Genetics and Biochemistry, Faculty of Medical and Health Sciences, Tel Aviv University, Tel Aviv 69978, Israel

^3^ European Molecular Biology Laboratory, European Bioinformatics Institute (EMBL-EBI), Wellcome Genome Campus, Hinxton, Cambridge CB10 1SD, UK

^4^ Center for Applied and Translational Genomics (CATG), Mohammed Bin Rashid University of Medicine and Health Sciences, Dubai Health, Dubai, UAE

^5^ Department of Genetics, Stanford University, Palo Alto, CA 94304 USA

^6^ Department of Genetics, Washington University School of Medicine, St. Louis, MO 63110, USA

^7^ Genome Informatics Section, Center for Genomics and Data Science Research, National Human Genome Research Institute, National Institutes of Health, Bethesda, MD 20892, USA

^8^ Division of Genome Sciences, National Human Genome Research Institute, Bethesda, MD 20871 USA

^9^ UC Santa Cruz Genomics Institute, University of California, Santa Cruz, CA 95060, USA

^10^ Department of Genome Sciences, University of Washington School of Medicine, Seattle, WA 98195, USA

^11^ The Vertebrate Genome Laboratory, The Rockefeller University, New York, NY 10065, USA

^12^ Bioinnovation and Genome Sciences, The Translational Genomics Research Institute (TGen), Phoenix, AZ 85004, USA

^13^ Human Technopole, Milan, Italy

^14^ Institute for Genomic Health, Icahn School of Medicine at Mount Sinai, New York, NY 10029, USA

^15^ Department of Computer and Information Science and Engineering, University of Florida, Gainesville, FL 32611, USA

^16^ Canadian Center for Computational Genomics, McGill University, Montréal, QC H3A 0G1, Canada

^17^ Department of Human Genetics, McGill University, Montréal, QC H3A 0G1, Canada

^18^ Victor Phillip Dahdaleh Institute of Genomic Medicine, Montréal, QC H3A 0G1, Canada

^19^ Department of Genetics, Genomics and Informatics, University of Tennessee Health Science Center, Memphis, TN 38163, USA

^20^ Google LLC, Mountain View, CA 94043, USA

^21^ Institute of Clinical and Translational Sciences, University of California, Irvine, CA 92697, USA

^22^ Quantitative and Computational Biology, University of Southern California, Los Angeles, CA 90089, USA

^23^ Department of Biomedical Informatics and Data Science, Yale School of Medicine, New Haven, CT 06510, USA

^24^ Department of Biology, University of Florence, Sesto Fiorentino, FI 50019, Italy

^25^ Department of Ecology and Evolutionary Biology, University of California, Santa Cruz, CA 95060, USA

^26^ Arizona State University, Consortium for Science, Policy & Outcomes, Washington, DC 20006, USA

^27^ Center for Digital Medicine, Heinrich Heine University Düsseldorf, Düsseldorf, NRW, DE

^28^ Department for Endocrinology and Diabetology at the Medical Faculty and University Hospital Düsseldorf, Heinrich Heine University Düsseldorf, Düsseldorf, NRW, DE

^29^ Paul-Langerhans-Group Computational Diabetology, German Diabetes Center (DDZ) and Leibniz Institute for Diabetes Research, Düsseldorf, NRW, DE

^30^ Wellcome Sanger Institute, Genome Campus, Hinxton, CB10 1RQ, UK

^31^ Department of Genetics, University of Cambridge, Cambridge, CB2 3EH, UK

^32^ Institute for Medical Biometry and Bioinformatics, Medical Faculty and University Hospital Düsseldorf, Heinrich Heine University, Düsseldorf, NRW, DE

^33^ Howard Hughes Medical Institute, Chevy Chase, MD 20815, USA

^34^ ISEM, Univ Montpellier, CNRS, IRD, Montpellier, FR

^35^ Institut Universitaire de France, Paris, FR

^36^ Department of Computer Science and Engineering, University of California San Diego, La Jolla, CA 92093, USA

^37^ Department of Bioethics & Humanities, University of Washington School of Medicine, Seattle, WA 98195, USA

^38^ Department of Genetics, Epigenetics Institute, Perelman School of Medicine, University of Pennsylvania, Philadelphia, PA 19104, USA

^39^ Department of Data Science, Dana-Farber Cancer Institute, Boston, MA 02215, USA

^40^ Department of Anthropology, University of Kansas, Lawrence, KS 66045, USA

^41^ School of Health Sciences, University of Manchester, Manchester M13 9PL, UK

^42^ Traditional, ancestral and unceded territory of the Gabrielino/Tongva peoples, Institute for Society & Genetics, University of California, Los Angeles, Los Angeles, CA 90095, USA

^43^ Traditional, ancestral and unceded territory of the Gabrielino/Tongva peoples, Institute for Precision Health, David Geffen School of Medicine, University of California, Los Angeles, Los Angeles, CA 90095, USA

^44^ Traditional, ancestral and unceded territory of the Gabrielino/Tongva peoples, Division of General Internal Medicine & Health Services Research, David Geffen School of Medicine, University of California, Los Angeles, Los Angeles, CA 90095, USA

^45^ Department of Electrical Engineering and Computer Science, University of California, Berkeley, Berkeley, CA 94720, USA

^46^ Department of Biomedical Data Science, Stanford University School of Medicine, Stanford, CA 94305, USA

^47^ Medical and Population Genomics Lab, Sidra Medicine, Doha, Qatar

^48^ Department of Molecular Cell and Developmental Biology, University of California, Santa Cruz, CA, USA

^49^ Montreal Heart Institute, Montréal, QC, Canada

^50^ Department of Pediatrics, University of California San Diego, La Jolla, CA 92093, USA

^51^ Center for Genomic Health, Yale University School of Medicine, New Haven, CT 06510, USA

^52^ Department of Genetics, Yale University School of Medicine, New Haven, CT 06510, USA

^53^ Department of Biomedical Informatics, Harvard Medical School, Boston, MA 02115, USA

^54^ Department of Evolution and Ecology and the Center for Population Biology, University of California, One Shields, Davis, CA 95616, USA

^55^ Department of Computer Science, Johns Hopkins University, Baltimore, MD 21218, USA

^56^ Department of Pediatrics, Division of Genetics, School of Medicine, University of California, Irvine, CA 92697, USA

^57^ Sun Yat-sen University, Guangzhou, China

^58^ Edison Family Center for Genome Sciences & Systems Biology, Washington University School of Medicine, St. Louis, MO 63110, USA

^59^ Department of Biology and Center for Medical Genomics, Penn State University, University Park, PA 16802, USA

^60^ Division of Medical Genetics, Department of Medicine, University of Washington School of Medicine, Seattle, WA 98195, USA

^61^ The Jackson Laboratory for Genomic Medicine, Farmington, CT 06032, USA

^62^ Department of Biology, Penn State University, University Park, PA 16802, USA

^63^ Department of Biomedical Science, College of Health Sciences, Qatar University, Doha, Qatar

^64^ Department of Genetic Medicine, Weill Cornell Medicine-Qatar, Doha, Qatar

^65^ IRSD - Digestive Health Research Institute, University of Toulouse, INSERM, INRAE, ENVT, UPS, Toulouse, FR

^66^ MATCH biosystems, S.L., Elche, Spain

^67^ Universidad Miguel Hernández de Elche, Elche, Spain

^68^ Department of Computational Biology and Medical Sciences, The University of Tokyo, Kashiwa, Chiba 277-8561, Japan

^69^ Department of Computer Science, University of Pisa, Pisa, Italy

^70^ Law School, University of Wisconsin-Madison, Madison, WI 53706, USA

^71^ Morgridge Institute for Research, Madison, WI 53715, USA

^72^ Institute of Genetics and Biomedical Research, UoS of Milan, National Research Council, Milan, Italy

^73^ Department of Biomedical Engineering, Johns Hopkins University, Baltimore, MD 21218, USA

^74^ Department of Genetic Medicine, Johns Hopkins University School of Medicine, Baltimore, MD 21205, USA

^75^ Genome Biology Unit, European Molecular Biology Laboratory (EMBL), Heidelberg, DE

^76^ Institute for Molecular Medicine Finland, Helsinki Institute of Life Science, University of Helsinki, Helsinki, Finland

^77^ The Center for Bio- and Medical Technologies, Moscow, RUS

^78^ Centre for Biomedical Research and Technology, HSE University, Moscow, RUS

^79^ Department of Biology, Johns Hopkins University, Baltimore, MD 21218, USA

^80^ Coriell Institute for Medical Research, Camden, NJ 08103, USA

^81^ University of Amsterdam, Amsterdam, Netherlands

^82^ School of Clinical Medicine, University of Cambridge, Cambridge, CB2 0SP, UK

^83^ Center for Genomic Discovery, Mohammed Bin Rashid University, Dubai Health, UAE

^84^ Dubai Health Genomic Medicine Center, Dubai Health, UAE

^85^ GenomeArc Inc, Mississauga, ON, Canada

^86^ Department of Biology and Biotechnologies “Charles Darwin”, University of Rome “La Sapienza”, Rome 00185, IT

^87^ Department of Human Genetics and Utah Center for Genetic Discovery, University of Utah, Salt Lake City, UT, USA

^88^ Center for Genomics, Loma Linda University School of Medicine, Loma Linda, CA 92350, USA

^89^ PacBio, Menlo Park, CA 94025, USA

^90^ The first affiliated hospital of Xi’an Jiaotong University, Xi'an Jiaotong University, Xi’an, Shaanxi, 710049, China
